# Impact and mitigation of sampling bias to determine viral spread: evaluating discrete phylogeography through CTMC modeling and structured coalescent model approximations

**DOI:** 10.1101/2022.07.07.498932

**Authors:** Maylis Layan, Nicola F. Müller, Simon Dellicour, Nicola De Maio, Hervé Bourhy, Simon Cauchemez, Guy Baele

## Abstract

Bayesian phylogeographic inference is a powerful tool in molecular epidemiological studies that enables reconstructing the origin and subsequent geographic spread of pathogens. Such inference is, however, potentially affected by geographic sampling bias. Here, we investigated the impact of sampling bias on the spatiotemporal reconstruction of viral epidemics using Bayesian discrete phylogeographic models and explored different operational strategies to mitigate this impact. We considered the continuous-time Markov chain (CTMC) model and two structured coalescent approximations (BASTA and MASCOT). For each approach, we compared the estimated and simulated spatiotemporal histories in biased and unbiased conditions based on simulated epidemics of rabies virus (RABV) in dogs in Morocco. While the reconstructed spatiotemporal histories were impacted by sampling bias for the three approaches, BASTA and MASCOT reconstructions were also biased when employing unbiased samples. Increasing the number of analyzed genomes led to more robust estimates at low sampling bias for CTMC. Alternative sampling strategies that maximize the spatiotemporal coverage greatly improved the inference at intermediate sampling bias for CTMC, and to a lesser extent, for BASTA and MASCOT. In contrast, allowing for time-varying population sizes in MASCOT resulted in robust inference. We further applied these approaches to two empirical datasets: a RABV dataset from the Philippines and a SARS-CoV-2 dataset describing its early spread across the world. In conclusion, sampling biases are ubiquitous in phylogeographic analyses but may be accommodated by increasing sample size, balancing spatial and temporal composition in the samples, and informing structured coalescent models with reliable case count data.

## Introduction

Over the past decade, Bayesian discrete phylogeographic inference has greatly benefited viral epidemiological studies in unraveling the origin and subsequent spread of viral epidemics (Faria et al. 2019; Lemey et al. 2020; Lu et al. 2021), the spatial processes driving viral spread (Müller et al. 2021), and environmental and human-related factors associated with viral spread (Lemey et al. 2014; Dudas et al. 2017; He et al. 2022). BEAST is a popular Bayesian phylodynamics software package commonly used in the analysis of time-stamped viral molecular sequences. It offers different discrete phylogeography approaches: a popular and computationally efficient discrete phylogeographic inference approach that makes use of continuous-time Markov chain (CTMC) modeling (Lemey et al. 2009), also known as the discrete trait analysis or DTA, and the structured coalescent model under its exact and approximated forms (Vaughan et al. 2014; De Maio et al. 2015; Müller et al. 2018). CTMC models migration between discrete locations in the same way as nucleotide substitutions are modeled. In other words, geographical locations are modeled as a neutral trait that evolves on top of the tree from the root to the tips. As such, CTMC modeling does not explicitly model the branching process that gave rise to the tree. In contrast, the structured coalescent model - which is an extension of the coalescent model to a structured population - is a tree-generating model that explicitly models how lineages coalesce within and migrate between subpopulations from present to past. Two computationally efficient approximations of the structured coalescent model are available in BEAST2: the Bayesian structured coalescent approximation (BASTA) (De Maio et al. 2015) and the marginal approximation of the structured coalescent (MASCOT) (Müller et al. 2018). Currently, they both assume constant prevalence through time for each deme/population, while the CTMC approach does not (Lemey et al. 2009).

Bayesian discrete phylogeography approaches are complementary to mathematical modeling and epidemiological studies, and particularly informative when epidemiological data are scarce. In such contexts, viral genetic sequences are expected to compensate for the lack of epidemiological data. However, genetic samples may constitute a biased snapshot of the underlying viral spread, especially when isolated through passive surveillance systems. The impact of such sampling bias on discrete phylogeographic inference has been discussed and examined ever since. Indeed, CTMC estimates were suspected to be biased towards the most sampled location (Lemey et al. 2009) and, later, sampling heterogeneity was shown to inform the posterior, and more specifically the migration parameters, which is not the case for BASTA (De Maio et al. 2015). In BASTA, sampling evenness is not informative as such and the estimated migration rates are more correlated to the true values under simulated biased and unbiased conditions compared to CTMC (De Maio et al. 2015). As a result, BASTA has been argued to be more robust to sampling bias (De Maio et al. 2015). Nevertheless, the structured coalescent model is known to be sensitive to unsampled locations, known as ghost demes (Beerli 2004; Ewing and Rodrigo 2006; De Maio et al. 2015). In parallel, several studies tested alternative strategies to mitigate the potential effects of sampling bias, mostly focusing on CTMC as it was shown to be potentially less robust to sampling bias compared to the structured coalescent model. Downsampling that was tested early on but was limited to large datasets (Lemey et al. 2014; Yang et al. 2019) rapidly became a prerequisite in any SARS-CoV-2 data analysis study due to the large number of available sequences and the high sampling heterogeneity between countries (Hodcroft et al. 2021). However, Magee and Scotch (2018) showed that inference accuracy rapidly plateaus when using up to 25-50% of the sequence data available (Magee and Scotch 2018). Other studies aimed at improving inference accuracy by integrating additional reliable epidemiological data. For example, CTMC was extended to incorporate information on the recent migration events using individual travel records (Lemey et al. 2020; Hong et al. 2021). More recently, a simulation study focused on quantifying the impact of sampling bias on the predicted location of internal nodes, the prediction of migration events that lead to large local spread as well as on the estimation of migration rates in a maximum likelihood framework (Liu et al.). The authors showed that prediction accuracy actually depends on multiple factors: the underlying migration rate, the magnitude of sampling bias and the magnitude of traveler sampling. Importantly, they observed a lower relative accuracy with biased samples and when samples overrepresent travelers. Concerning the structured coalescent model, Müller et al. informed the deme population sizes with reliable case count data from the 2014 Ebola epidemic in Sierra Leone using MASCOT (Müller et al. 2019). This allows modeling time-varying population sizes instead of assuming constant population sizes over time. Sampling bias is also a concern in continuous phylogeography analyses in which other mitigation approaches were tested. Recently, Dellicour et al. downsampled SARS-CoV-2 genomic records from New York City based on hospitalisations rather than case counts to analyze representative samples irrespective of testing effort and strategy (Dellicour et al. 2021), Kalkauskas et al. incorporated sequence-free samples from unsampled areas (Kalkauskas et al. 2021), and Guindon and De Maio explicitly modeled sampling strategy in the data likelihood (Guindon and De Maio 2021).

Whereas numerous studies tested strategies to deal with sampling bias, the impact of sampling bias on discrete phylogeographic reconstructions remains insufficiently characterized. Here, we compare the performance of the different phylogeographic methods using simulated viral epidemics using a stochastic metapopulation model, based on rabies virus (RABV) epidemics in dogs in Morocco. We investigated the impact of sampling bias on the spatiotemporal reconstruction of these viral epidemics using CTMC, BASTA, and MASCOT, with the latter two assuming populations to stay constant over time. Next, we explored different approaches to mitigate sampling bias, maximizing the spatial and/or temporal coverage of the sample, and informing the deme sizes under MASCOT with the true (time-varying) case count data per location. The latter is to test to what degree biases originating from assuming constant population sizes over time can be mitigated by allowing them to vary over time. Finally, we applied the three algorithms to two empirical datasets: a dataset of RABV sequences isolated in the Philippines islands between 2004 and 2010, and a global dataset of SARS-CoV-2 genomes of the early spread of the pandemic.

## Results

### Simulation framework

We simulate RABV epidemics across three or seven locations using a stochastic metapopulation model (Figure 1A) whose connectivity matrix is parameterized using human population mobility that we estimated by fitting the radiation model of (Simini et al. 2012) to human population density data from (WorldPop) (Figure 1B). As each location is associated to a specific deme/population, we refer to the two simulation frameworks as the three or seven demes framework for the remainder of the text. We simulate 50 epidemics that start with the introduction of a single case and lead to at least 60,000 cases over a 30-year period (Figure 1C). On top of the transmission chains, we simulate viral genomes for each case and then sub-sample starting one year after the introduction of the index case either 150 or 500 sequences in a biased or unbiased fashion (Figure 1D). We then perform Bayesian discrete phylogeographic analysis on the geolocated and time-stamped sequence alignments before comparing the true and reconstructed evolutionary and migration histories for each discrete phylogeographic approach. Importantly, the vast majority of samples in the three demes framework contain at least one sequence of each deme which is not the case for the seven demes framework for which sampling bias often leads to unsampled locations, also called “ghost” demes.

**Figure 1.**
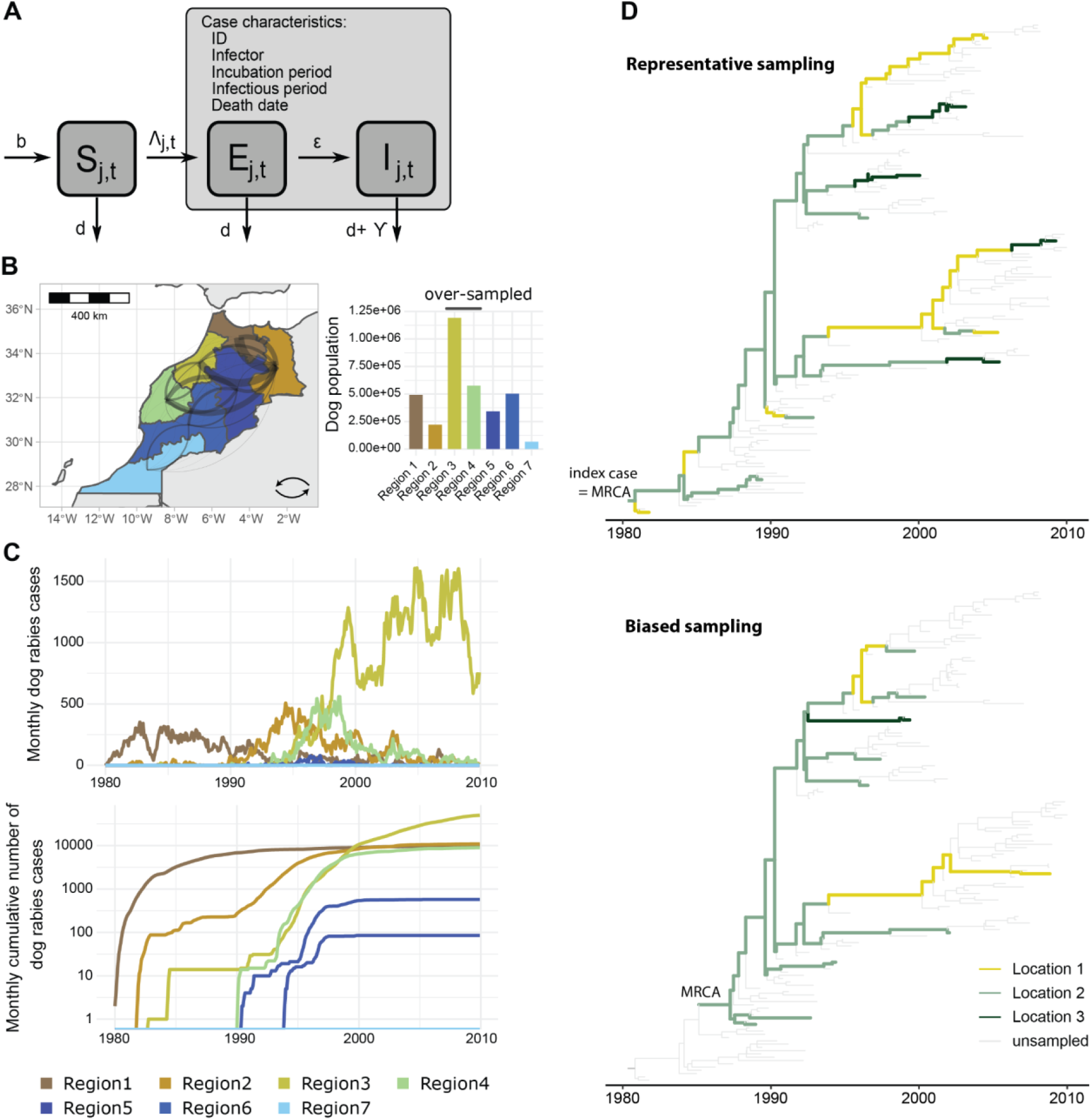
Rabies virus (RABV) epidemic simulation framework. We simulate realistic epidemics by emulating the scenario of RABV spread in dog populations in Morocco. **A:** Metapopulation model of rabies spread in dogs. In each geographical location *j*, the dog population is divided into three compartments: susceptible, exposed but yet not infectious, and infectious individuals. Individuals are born at rate *b* and die from natural causes at rate *γ*. The rate of infection corresponds to the per-capita force of infection *∧*_*j,t*_ that aggregates the force of infection from infectors in location j and all the other locations. Individuals become infectious at rate *ε*. We identify all infected individuals and simulate their infector, incubation period, infectious period and date of death. **B:** Connectivity between the seven arbitrary Moroccan regions estimated by the radiation model and estimated dog population size per region. Curvature indicates flux direction. **C:** Example for one simulation of the prevalence (first row) and cumulative number (second row) of rabid cases per month and location. **D:** Graphical illustration of the potential impact of sampling bias on the reconstruction of the phylogenetic relationships between viral samples over an epidemic, assuming no intra-host evolution.

### Robust estimation of the phylogeny and genetic parameters with respect to sampling bias

While the focus of our simulation study is on reconstructing the spatial spread, we first assess the potential impact of sampling bias on estimating the phylogeny itself, as well as the evolutionary parameters (Figure 2A). The phylogeny of the simulated pathogen is not impacted by sampling bias when using CTMC, BASTA, and MASCOT (Figure 2A and S1). In addition, the average evolutionary rate (Figure 2C), the stationary nucleotide frequencies (Figures S2-5), and the ratio of transition-transversion rates (Figure S6) are all well estimated at any level of sampling bias.

**Figure 2.**
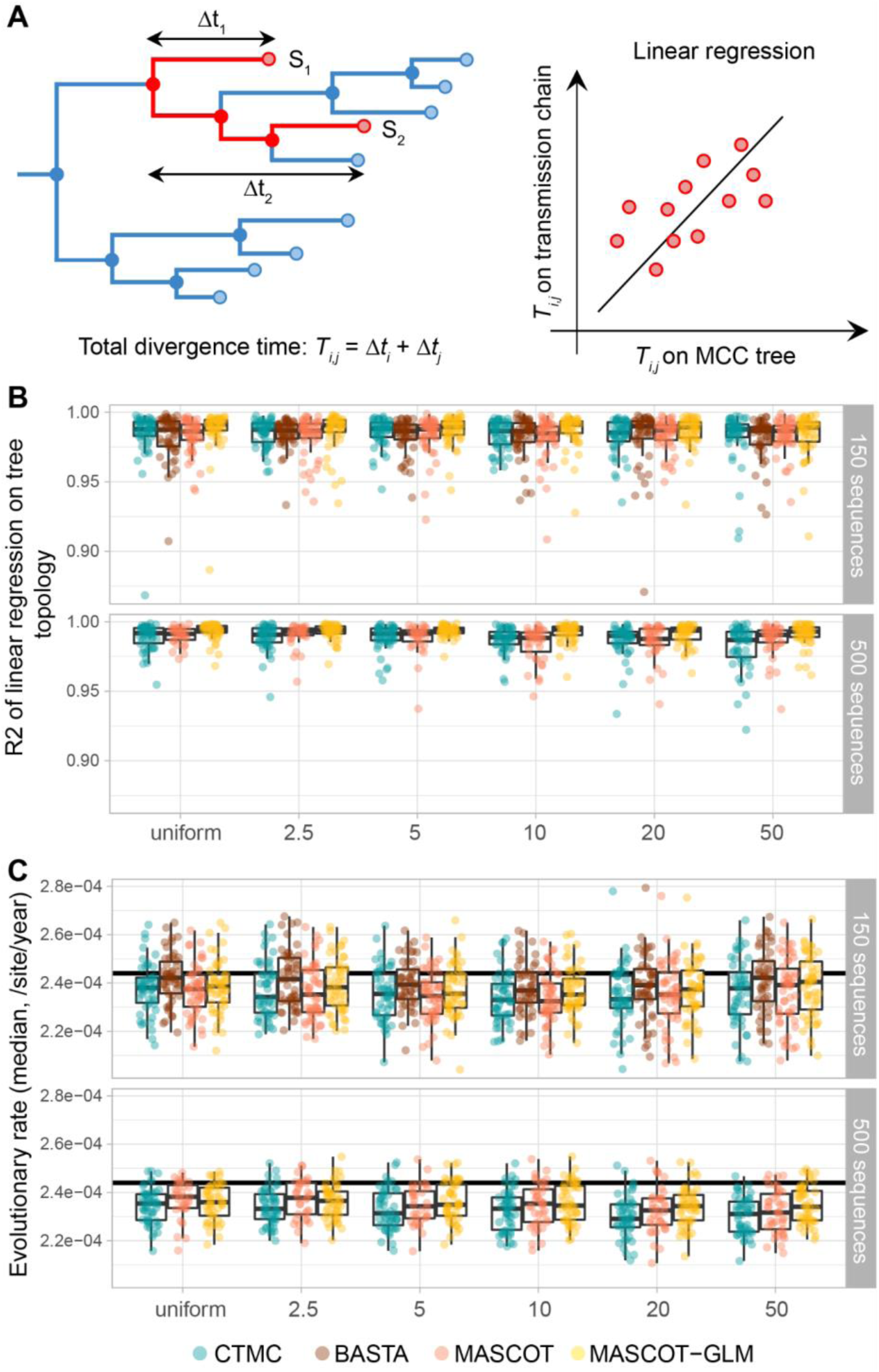
Estimation of genetic and phylogenetic parameters under spatially-biased sampling conditions. **A:** Comparison of the simulated transmission chain and the estimated maximum clade credibility (MCC) tree topologies. For the estimated and simulated topologies, we computed the total divergence time between every pair of sampled tips. We compared the two using linear regression. **B:** Pearson’s determination coefficient of the pairwise divergence time between the simulated transmission chain and the MCC tree. **C:** Estimation of the evolutionary rate. Each dot corresponds to the median estimate of the evolutionary rate in one simulation (*n* = 50 per sampling protocol and sample size).

### Spatiotemporal history reconstruction in (un)biased conditions

As the inferred spatiotemporal histories of lineages cannot be compared in a unique simple way between the different approaches, we use four types of summary statistics: i) the total migration counts - corresponding to Markov jumps in the case of CTMC and their equivalent for BASTA and MASCOT - that account for multiple migration events along the tree branches (Figure 3), ii) the lineage migration counts (Figure S7). iii) the lineage introduction dates into the sampled locations (Figure 4), and iv) the root location (Figure 5). We evaluate the performance of the phylogeographic models using five metrics: the correlation between true and estimated values, the proportion of estimated parameters for which the true value is in the 95% highest posterior interval (HPD) that we refer to as the calibration, the mean relative bias, the mean relative 95% HPD width, and the weighted interval score (WIS). The WIS is a generalization of the absolute error accounting for estimation uncertainty (Bracher et al. 2021). The smaller the WIS, the better the inference. It is widely used to evaluate epidemic forecasts and favors estimates that are slightly biased but with a narrow confidence interval compared to estimates without bias but very large uncertainty (Bracher et al. 2021).

**Figure 3.**
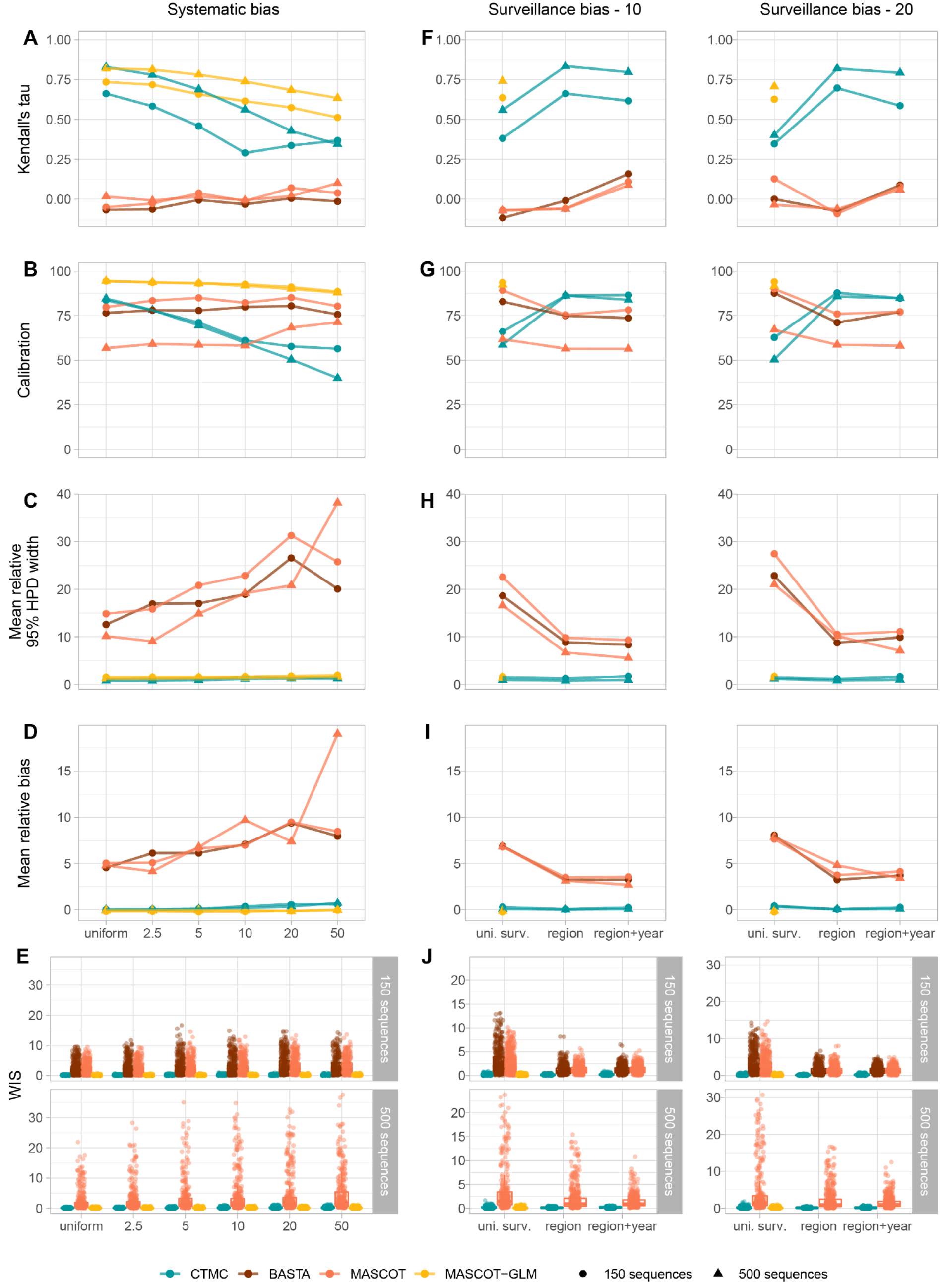
Impact and mitigation of spatial bias on the estimation of the total migration counts. **A-E:** Impact of the increasing levels of spatial bias on the correlation, the calibration, the mean relative 95% highest posterior density (HPD) width, the mean relative bias, and the WIS between the simulated and the estimated total migration counts. **F-J**: Mitigation of the impact of spatial bias on the correlation, the calibration, the mean relative 95% HPD width, the mean relative bias, and the WIS between the simulated and estimated total migration counts by using alternative sampling strategies. In the left and right columns, samples are drawn from biobanks with an underlying bias of 10 and 20, respectively. Overall, the algorithms correctly estimate the total migration counts when the correlation and the calibration are high (close to 1 and 100, respectively) and when the mean relative 95% HPD width, the mean relative bias, and the WIS are close to zero. Finally, the mean relative bias and the mean relative 95% HPD width are not defined when the true value is null. We removed 612 out of 3,600 and 380 out of 3,600 simulated migration events in the small and large samples, respectively, due to null true values.

**Figure 4.**
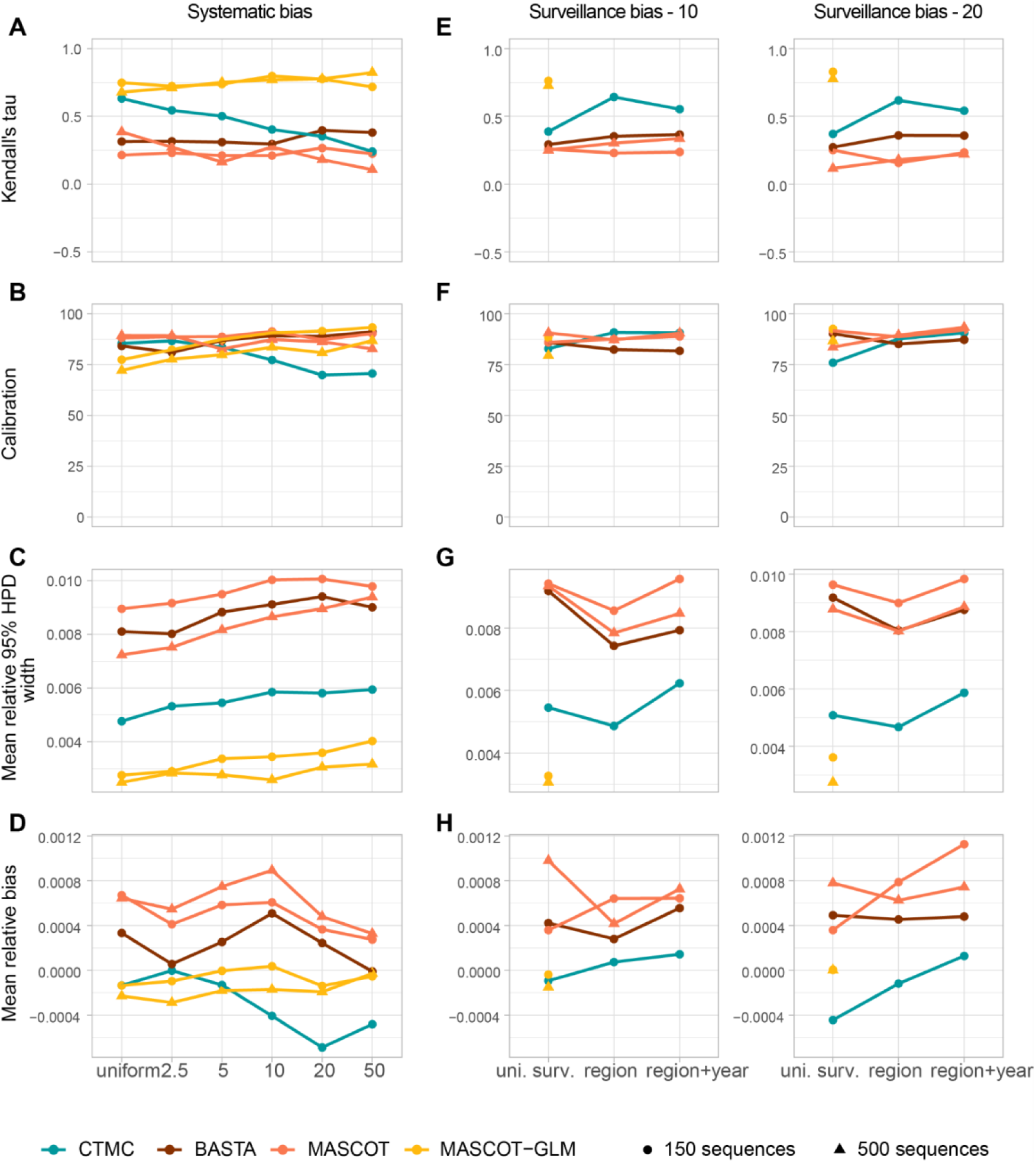
Impact and mitigation of spatial bias on the estimation of the lineage introduction dates. **A-D:** Impact of the increasing levels of spatial bias on the correlation, the calibration, the mean relative 95% highest posterior density (HPD) width, and the mean relative bias between the simulated and the estimated introduction dates. Uniform samples are representative of the simulated spatiotemporal dynamics of the virus. Samples 2.5, 5, 10, 20 and 50 samples biased towards Regions 3 and 4. Samples 2.5 and 5 correspond to low levels of bias, samples 10 and 20 to intermediate levels of bias and sample 50 to high levels of bias. **E-H:** Mitigation of the impact of spatial bias on the correlation, the calibration, the mean relative 95% HPD width, and the mean relative bias between the simulated and estimated introduction dates by using alternative sampling strategies. In the left and right columns, samples are drawn from biobanks with an underlying bias of 10 and 20, respectively.

**Figure 5.**
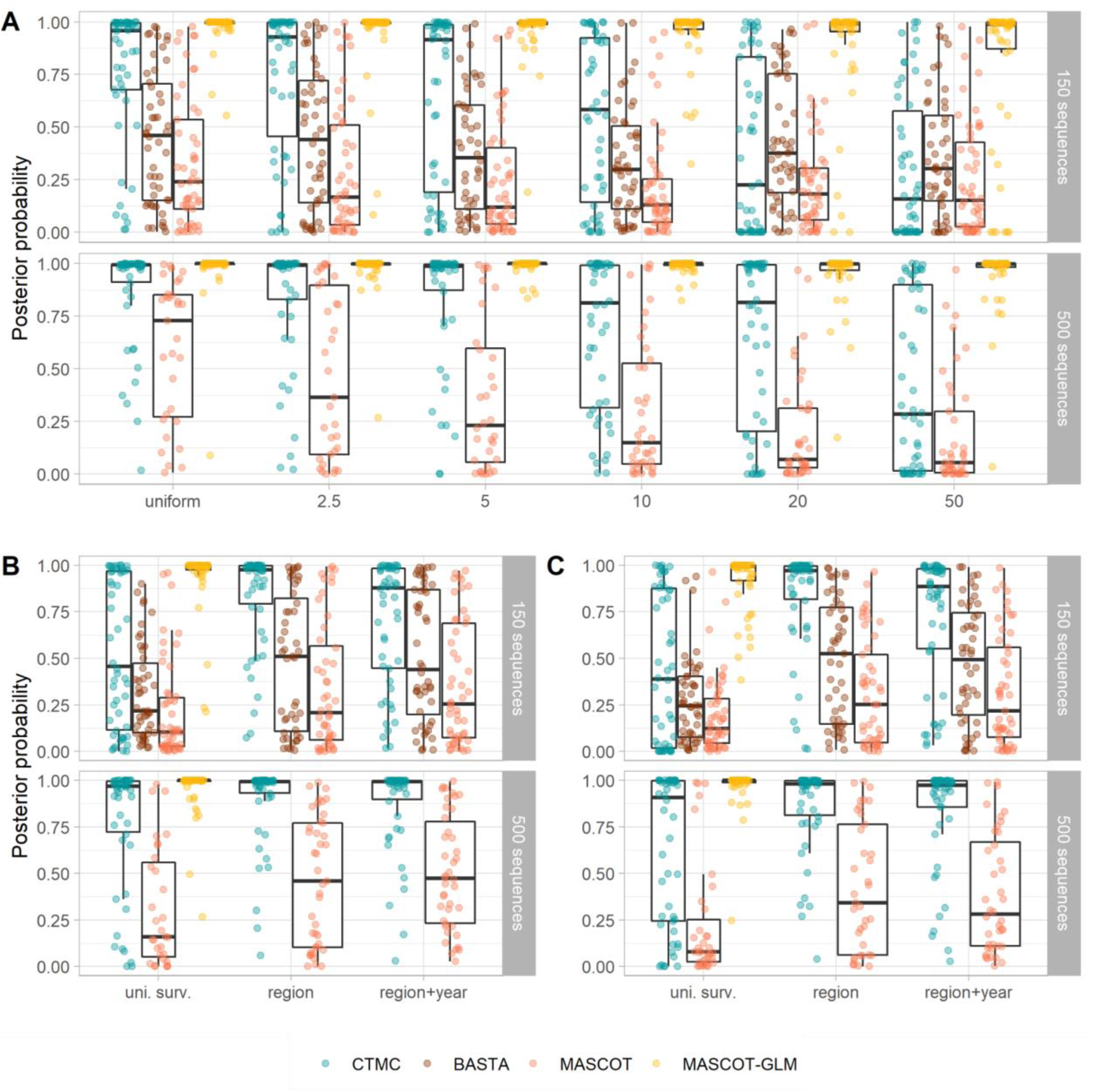
Impact and mitigation of spatial bias on the estimation of the root location. **A:** Posterior probability of estimating the true root state for increasing levels of biais. **B-C:** Mitigation of the effects of spatial biais using alternative sampling strategies under a surveillance bias of 10 and 20, respectively. Each dot corresponds to the median root state posterior probability in one simulation (*n* = 50 per sampling protocol and sample size).

First, we assess the reconstruction of the spatial process in the absence of sampling bias. In the unbiased/representative (uniform) scenario, CTMC correctly estimates the four types of parameters. Indeed, the correlation between the true and estimated parameter values is high, and the WIS is close to zero. BASTA and MASCOT show no correlation for the total migration counts on uniform samples and higher WIS compared to CTMC (Figures 3A and 3E) indicating biased median estimates and higher uncertainty around the point estimate. The correlation is over 0.5 when we consider the lineage migration counts under the three and seven demes frameworks. This suggests that BASTA and MASCOT only partly recover the global migration process in the absence of sampling bias (Figures S7 and S15). Overall, CTMC outperforms BASTA and MASCOT when the sampling is representative of the true underlying transmission process, as BASTA and MASCOT only recover the big picture of the migration process.

Secondly, we evaluate how phylogeographic algorithms perform under increasing levels of bias. While CTMC satisfyingly estimates the total migration counts in the absence of sampling bias, the correlation and the calibration drop rapidly with bias and the mean relative 95% HPD width tends to decrease suggesting that bias strongly impacts CTMC estimates (Figure 3A-C). Nevertheless, the WIS and the mean relative bias remain smaller than those of BASTA and MASCOT, even at high levels of bias. Consequently, CTMC leads to median estimates that are closer to the true values but with 95% HPDs that are too narrow. It leads to a biased picture of the geographical process with some transition events that are drastically underestimated (Figure S7). BASTA and MASCOT less accurately estimate the total migration counts with no correlation between simulated and estimated values. They are also less confident with an average 95% HPD width that is ten to thirty times higher compared to CTMC. This uncertainty is exacerbated in large samples analyzed with MASCOT in the seven demes framework, for which almost 30% (87 out of 300) of the chains have low ESS values often due to bimodal structured coalescent posterior density. Additionally, BASTA and MASCOT partly recover the global migration process (lineage migration counts) even at high levels of bias since correlation and calibration are not impacted by bias (Figure S8). When we consider transmission dynamics between three demes, BASTA and MASCOT yield higher correlation levels than in the seven demes scenario (Figures S15 and S16). Overall, the WIS indicate better performance of CTMC over BASTA and MASCOT.

When it comes to the estimation of lineage introduction dates, BASTA seems to outcompete CTMC and MASCOT under the three demes framework (Figure S17) but not under the seven demes framework (Figure 4A-E). In the three demes framework, the uncertainty around the median estimate remains high for BASTA and MASCOT, and the correlation and the calibration are barely affected by bias for BASTA, contrary to CTMC and MASCOT. In the seven demes framework, correlation is low for both BASTA and MASCOT but not affected by bias. CTMC performs poorly with a sharp decrease in both correlation and calibration in the three demes framework, and a slighter decrease in the seven demes framework. It also tends to estimate more ancient lineage introduction dates compared to BASTA and MASCOT in both frameworks. Of note, samples of 500 sequences displayed a higher correlation than samples of 150 sequences at low and intermediate levels of bias for CTMC (conditions 2.5, 5 and 10 in Figures 4A and S8A).

Finally, we analyze the potential impact of sampling bias on root location estimation (Figure 5A and S18A). Of note, the location probability of the true root location is very heterogeneous among the 50 simulated epidemics when there is no or little sampling bias, notably for the two approximations of the structured coalescent model. Root location prediction by CTMC is affected by sampling bias, notably in the three demes framework (Figure S18), which is in agreement with previous findings (De Maio et al. 2015). As for the other parameters, BASTA and MASCOT perform less well compared to CTMC, at any level of bias. On the other hand, sampling bias moderately worsens their estimates. They also perform relatively better in the three demes framework.

### Sample balancing mitigates the impact of sampling bias

We test alternative sampling strategies in order to mitigate the impact of sampling bias. Large and biased samples of 5,000 sequences that we refer to as biobanks were generated, then discrete phylogeographic analyses were carried out on sub-samples of 150 or 500 sequences, which aimed at reproducing real life situations. For example, researchers may have access to numerous viral specimens from biobanks but cannot analyze all of them due to computational limitations, potential underlying biased sampling that may lead to spurious results, or financial limitations.

Similar to the analyses on systematically biased samples, the estimation of the total migration counts (Figure 3F-J), lineage migration counts (Figure S6F-J), lineage introduction dates (Figures 4E-H), and root location posterior probabilities (Figure 5B-C) is strongly impacted in biased sub-samples (uniform surv.) for the three approaches. By maximizing the spatial (region) or the spatiotemporal coverage (region+year), the correlation of lineage migration counts increased substantially for the CTMC even when the underlying sampling bias was high (weight=20, i.e. sequences from oversampled regions are 20 times more likely to be samples), and to a lesser extent for BASTA and MASCOT. Calibration remained high for BASTA and MASCOT, as shown earlier (Figures 3G and 4F) while it considerably improved for CTMC. Estimates of the lineage migration counts by BASTA and MASCOT are improved in the region and region+year conditions compared to the uniform surv. condition, illustrated by a decreased mean relative 95% HPD width and decreased WIS. Still, performance remained lower than for CTMC. In the three demes framework, we obtain even stronger improvements in terms of correlation and decreased mean relative bias for BASTA and MASCOT (Figure S15). Overall, subsampling strategies that maximize the spatial or spatiotemporal coverage considerably improved the inference of the geographical spread by the CTMC, and improved inference under BASTA and MASCOT to a lesser extent.

### True incidence data as a predictor of the time-varying deme sizes mitigate sampling bias in MASCOT

Due to the lack of statistical power (data not shown), we have forced all deme sizes to be equal in BASTA and MASCOT and to be constant over time, the latter being currently the default assumption of both structured coalescent models. This hypothesis is potentially impactful given that deme sizes are directly related to the migration history in the structured coalescent model (De Maio et al. 2015; Müller et al. 2018). To relax this assumption and allow for time-varying effective population sizes, we next use the monthly incidence data from our simulations as a predictor of the deme sizes over time in the generalized linear model (GLM) extension of MASCOT and denote the resulting model as MASCOT-GLM. This approach is only available for MASCOT, we can therefore not perform the same analysis for BASTA.

By accommodating for the time variations of deme sizes, the correlation, mean relative 95% HPD width, mean relative bias, and WIS are markedly improved with MASCOT-GLM compared to BASTA and MASCOT for the total migration counts (Figure 3A-E), lineage migration counts (Figure S7A-E), and lineage introduction dates (Figure 4A-E) even under high levels of sampling bias in the systematically biased conditions (scenarios 5, 10, 20, and 50) and in the biased sub-samples (uniform surv.). In addition to the strong correlation between simulated and estimated values, the uncertainty around the true value and the bias (mean relative bias and WIS) are low compared to BASTA and MASCOT with constant population sizes.

### Analysis of the spread of RABV in the Philippines

As a case study to compare the performance of the three algorithms, we analyze the spread of RABV in dog populations between six Philippine islands (Luzon, Catanduanes, Oriental Mindoro, Cebu, Negros Oriental, and Mindanao) using 233 sequences of the RABV glycoprotein gene isolated between 2004 and 2010 (Saito et al. 2013; Tohma et al. 2014). Discrete phylogeography is particularly adapted here to model transmission in animal populations across an archipelago. In this dataset, sampling is highly heterogeneous across the different islands: Luzon represents up to 65% of the total dataset while Oriental Mindoro is represented only by a single sequence (Figure S19). This heterogeneity is very unlikely to be representative of the underlying transmission but rather due to case underreporting outside Luzon.

Previous studies on RABV in the Philippines suggested that although the circulating lineages likely circulate independently in the main islands (Saito et al. 2013; Tohma et al. 2014), inter-island transmission events can lead to sustained circulation in previously rabies-free islands (Tohma et al. 2016). Importantly, the patterns of spatial spread were evaluated using the CTMC at a finer spatial scale (Tohma et al. 2014). Here, the CTMC model also predicts a highly spatially-structured phylogeny with few migration events between islands. It reconstructs four island-specific clades located in Catanduanes, Luzon, Mindanao, and Negros Oriental with high node and location posterior support (Figure 6A). BASTA and MASCOT also predict the Catanduanes, Mindanao, and Negros Oriental clades with high node and location posterior support (Figure 6B-C). However, the migration history of the Luzon clade is more uncertain with potential intense migrations between Luzon and Oriental Mindoro islands, the most and least sampled islands, respectively. As shown in the simulations, CTMC might be overconfident compared to BASTA and MASCOT but the uncertainty of the two approximations of the structured coalescent model might be related to the pseudo-ghost demes, i.e. locations for which very few sequences are available. As we don’t have information regarding the number of cases over time, we could not apply MASCOT-GLM to this dataset.

**Figure 6.**
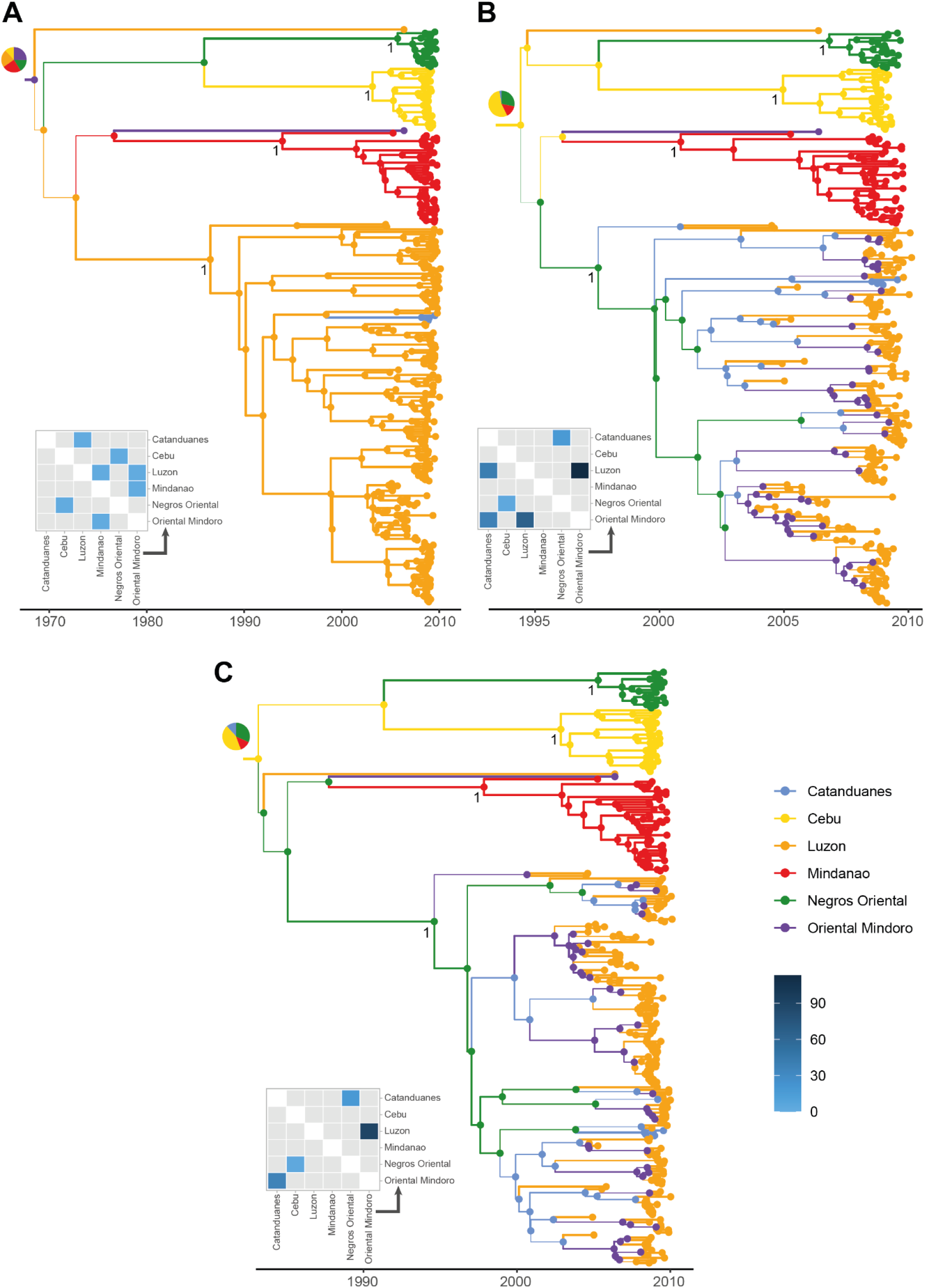
Maximum clade credibility (MCC) trees and median total migration counts estimated on the rabies dataset. **A-C:** MCC trees and median number of total migration counts estimated on the rabies dataset by CTMC, BASTA, and MASCOT, respectively. Branch width is proportional to the maximal ancestral location probability predicted by the algorithms, and branches are colored by the maximal ancestral location. Posterior support of the Negros Oriental, Catanduanes, Mindanao, and Luzon island lineages are reported. Pie charts displayed at root nodes represent the posterior probability distribution of the root location. Median estimates of the total migration counts are reported as heatmaps. Gray tiles correspond to transitions associated with a migration rate that is not statistically supported, i.e., with a Bayes factor lower than 3.

### Analysis of the early spread of SARS-CoV-2 across the world

In the context of zoonotic diseases, surveillance systems mostly rely on disease monitoring in human populations. Thus, there are typically no reliable estimates of the number of new cases in wild animal populations and, depending on the country and the species considered, domestic animal populations. However, for pathogens infecting the human population, such estimates are typically widely available, as is the case for SARS-CoV-2 and as has also been shown used previously when studying Dengue virus, HIV and West Nile virus (Gill et al. 2016; Dellicour, Lequime, et al. 2020).

To compare the phylogeographic reconstructions of the four algorithms tested above, we analyze a dataset of SARS-CoV-2 genomic sequences from the early stage of the pandemic (Lemey et al. 2020). In the original study, the initial wave of SARS-CoV-2 infections was investigated using a novel travel history-aware extension of the CTMC model, which we here refer to as CTMC-TRAVEL.

We again use the CTMC model, BASTA and MASCOT as well as MASCOT-GLM to analyze the dataset. MASCOT-GLM is informed using the seven-day moving average of case count data either from Our World In Data (Ritchie et al. 2020) or from the (World Health Organization (WHO)). MASCOT-GLM is then referred to as MASCOT-WID and MASCOT-WHO, respectively (Figure S20). Due to the low number of mutations accumulated in the SARS-CoV-2 genome at the start of the pandemic, the posterior support of internal nodes for each algorithm is low and the tree topology very uncertain (Morel et al. 2021). Besides, we do not intend to reconstruct the origins of SARS-CoV-2 which in any case cannot be addressed solely with phylogeographic analyses (Pipes et al. 2021). That is why our comparison focuses on the posterior support of four clades originally identified by (Lemey et al. 2020): clades A, A.1, B.1, and B.4. Whereas clades A.1, B.1, and B.4 are predicted with high posterior support by all algorithms, clade A is predicted with a satisfying posterior support only by CTMC (Figure S21). In general, CTMC and MASCOT-WHO predictions are closer to the original predictions than the other algorithms, in terms of tree topology (Figure S21A) and of total migration counts (Figure 7). As previously shown, BASTA and MASCOT lead to more uncertain ancestral migration histories with the extreme case of BASTA for which the posterior evolutionary rate and the structured coalescent density are bimodal. We report two maximum clade credibility (MCC) trees for BASTA, corresponding to the two modes of the evolutionary rate and structured coalescent density (Figures S21B-C and S22). The first mode of BASTA infers a tree topology and a migration history that are similar to CTMC and CTMC-TRAVEL. For example, the predicted location of the MRCA of the B.4 lineage is China for CTMC-TRAVEL, CTMC, and the 1st mode of BASTA, whereas it is located in Oceania by MASCOT and the 2nd mode of BASTA (Table S1). For the latter two reconstructions, most of the ancestral branches were not inferred to occur in China and, similarly to the RABV dataset, these approaches predict the least sampled locations (Africa and Oceania) to play a major role in the transmission process.

**Figure 7.**
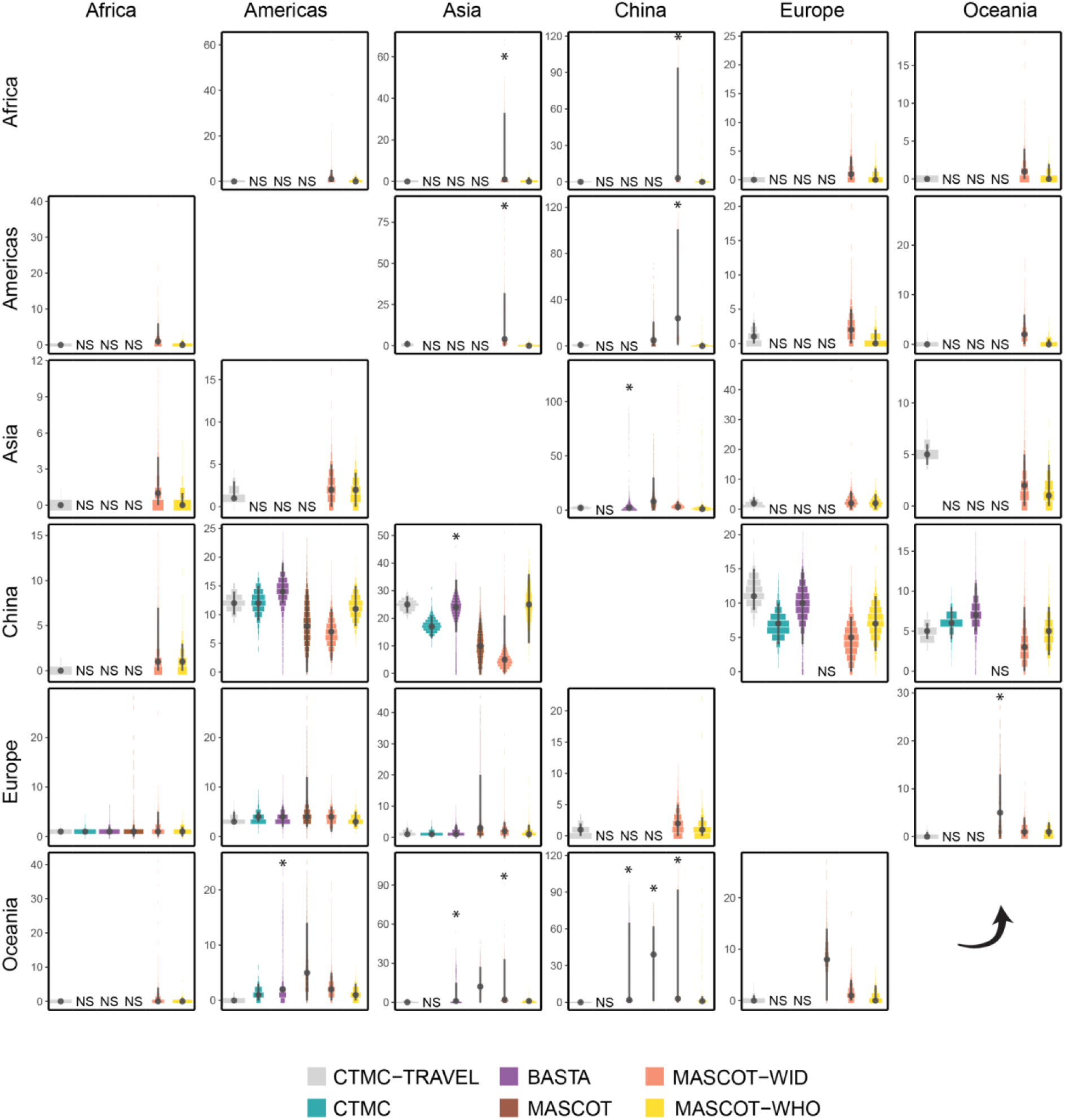
Posterior distributions of the total migration counts estimated on the SARS-CoV-2 data. Source locations are displayed by rows and destination locations by columns. For CTMC, BASTA and MASCOT, posterior distributions of the total migration counts with a Bayes factor (BF) < 3 are not depicted but marked as non-significant (NS). We identify bimodal marginal posterior distributions with a (*) and we report for each posterior distribution the median and 95% HPD. We normalize the width of the violin plots so that the cumulative density is equal to one.

We next incorporated incidence data from either Our World In Data or the WHO into MASCOT-GLM. Interestingly, the reconstructions differed strongly between the two datasets for incidence. While MASCOT-WID predictions are uncertain with multimodal total migration counts (Figure 7) and do not reflect the original spread from China (Figure S21E), MASCOT-WHO estimated migration counts that are close to the estimates of CTMC-TRAVEL (Figure 7) and its MCC tree is in agreement with the origin of the pandemic (Figure S21E). Importantly, the two datasets differ strongly in how well early cases are covered (Figure S20), with the WHO dataset being more representative of the incidence over time. Overall and as also suggested by our simulations, while the structured coalescent model, in principle, allows to mitigate sampling biases, it can itself be highly biased when the wrong population dynamics are assumed.

## Discussion

Sampling bias is a key challenge in phylodynamic inference (Frost et al. 2015), as in discrete phylogeography. In its early developments, the evaluation of the impact of sampling bias on Bayesian discrete phylogeography models was restricted by the availability of whole genomes (Lemey et al. 2009). The SARS-CoV-2 pandemic has led to a paradigm shift as genomic surveillance became part of routine surveillance systems around the world (Hodcroft et al. 2021). Here, we evaluated the impact of sampling bias on discrete phylogeography inference using simulated and real data to provide insightful knowledge on how sampling bias affects such inference and how it could be mitigated.

### Inference performance in absence of sampling bias

In our simulation study, genetic parameters (i.e., average evolutionary rate, stationary nucleotide frequencies, ratio of transition-transversion rates) are correctly estimated and tree topologies match the corresponding simulated transmission chains for all approaches. In addition, CTMC leads to high correlation between simulated and estimated spatiotemporal parameters as well as low relative and absolute error in absence of sampling bias. Overall, CTMC reconstructs the spatiotemporal histories well and its estimates are more accurate in large samples. BASTA and MASCOT do not correctly infer the spatiotemporal parameters in the seven demes framework but correlation between simulated and estimated total migration counts is slightly improved in the three demes framework while remaining lower than CTMC. This could result from three different causes. First, we assumed that all deme sizes are equal and constant over time in BASTA and MASCOT, the former to avoid overparameterization and the latter being the only available assumption in current implementations. Such a parameterization is more appropriate in the case of endemic circulation with limited time-varying dynamics such as local extinctions. However, large variations in time and local extinctions occur in our simulations meaning that we had to assume incorrect population dynamics in BASTA and MASCOT. This is confirmed by the better performance of MASCOT-GLM in uniform samples that accommodates for the true population dynamics. Secondly, we would expect BASTA and MASCOT to perform better on “even” samples that contain approximately as many sequences of each sampled location (De Maio et al. 2015). In our simulation study, uniform sampling does not imply an even representation of sampled locations. Indeed, locations where the virus has not circulated much are less represented. Such an effect is more pronounced in the seven demes framework than the three demes framework and we effectively observe poorer performances of BASTA and MASCOT in the seven demes framework. Finally, the structured coalescent model is known to be sensitive to ghost demes, i.e. unsampled locations (Beerli 2004; Ewing and Rodrigo 2006; De Maio et al. 2015). As we considered the sampling process to be naive of the number of affected locations, locations where the virus has not circulated much may remain unsampled. This is true for the seven demes framework only for which we observe poorer performance of BASTA and MASCOT compared to the three demes framework. However, the impact of ghost deme inclusion and potential misspecification on the estimation of the migration patterns remains unclear. While two studies showed that accounting for ghost demes in the structured coalescent model improves the inference of deme size (Beerli 2004; Ewing and Rodrigo 2006), Ewing and Rodriguo also showed that adding just a few sequences from the ghost deme leads to the overestimation of the migration rate (Ewing and Rodrigo 2006).

### Inference performance under sampling bias

We show that CTMC, BASTA, and MASCOT are impacted by spatial sampling bias in different ways. CTMC performance is dramatically impaired with increasing levels of sampling bias. This is directly linked to the geographical sampling frequencies that inform the likelihood of CTMC (De Maio et al. 2015). It also tends to be overconfident, and this overconfidence worsens with stronger sampling bias as previously shown (De Maio et al. 2015). However, the impact of sampling bias can be mitigated by either using large samples at low levels of sampling bias or controlling for sampling bias by balancing sample composition (region and region+year subsamples) at intermediate levels of sampling bias. These results were well-replicated in a simpler framework of transmission between three locations which rules out the confounding effect of the simulation complexity and unsampled locations on our results (see section 4 of the Supplementary Materials).

BASTA and MASCOT do not accurately estimate the total migration counts nor the lineage introduction dates in biased and unbiased conditions. Nevertheless, the overall migration process evaluated by the lineage migration counts is relatively well captured with a correlation around 0.5 that is not impacted by sampling bias contrary to CTMC in both the three demes and seven demes scenarios. We show that the approximations of the structured coalescent model are generally less confident than CTMC which is in agreement with a previous study (De Maio et al. 2015) and their uncertainty around median estimates increases with sampling bias. We also show that sample composition impacts the inference of BASTA and MASCOT in the three demes framework since correlation levels are strongly improved and bias and uncertainty are reduced for all spatiotemporal parameters in “even” samples (region and region+year), despite the underlying surveillance bias. Still, BASTA and MASCOT estimates display lower correlation with the simulated values, higher uncertainty and higher relative and absolute bias compared to CTMC. In the seven demes framework, the results are less clear which may be due to the presence of ghost demes. Interestingly, BASTA seems to outperform CTMC and MASCOT in the inference of the lineage introduction dates in the three demes framework. This result was however not replicated in the seven demes framework.

While structured coalescent methods potentially allow mitigating sampling biases as previously shown (De Maio et al. 2015), assuming incorrect population dynamics very likely introduces biases. These models currently assume constant population sizes which leads to biases in the case of true underlying complex population dynamics, which we salvaged by modeling these population dynamics more accurately, using a GLM approach, whenever the required data to do so were available. Indeed, using incidence data to inform population dynamics in MASCOT counteracts the impact of sampling bias even at high levels. This result also underlines that sampling frequencies do not inform the structured coalescent model when population dynamics are known (De Maio et al. 2015). It also shows that the inclusion of ghost demes is not necessary when the true population dynamics are used. Overall, our results showcase the importance of considering the assumptions of population dynamics on the ancestral state reconstruction in structured coalescent model approximations.

### Analysis of empirical RABV and SARS-CoV-2 datasets

We further compare the approaches on real datasets of RABV and SARS-CoV-2. As dog case counts were not available for RABV, we compare only CTMC, BASTA, and MASCOT. CTMC predicts a highly spatially-structured migration process whereas BASTA and MASCOT predict a non-parsimonious scenario. We observe similar results for the SARS-CoV-2 dataset. As we have set equal deme sizes in BASTA and MASCOT but a single tip is sampled for Oriental Mindoro in the RABV dataset and Africa in the SARS-CoV-2 dataset, the two algorithms compensate for location underrepresentation by estimating high backwards-in-time migration rates to the underrepresented location (Oriental Mindoro and Africa). Our results are in line with previous studies reporting strong differences between CTMC and the structured coalescent model on real datasets (De Maio et al. 2015; Dudas et al. 2018). However, there is also evidence in the literature of a good agreement between the two types of models (Faria et al. 2017; Brynildsrud et al. 2018; Yang et al. 2019; Mavian et al. 2020). Such similarities can result from sample composition (at least ten sequences per location in (Yang et al. 2019)), the parameters used for comparison (probability of clade ancestral location in (Faria et al. 2017)), prior information (information on the root location in (Brynildsrud et al. 2018)), or the underlying transmission dynamics. Besides, these studies focused on the overall migration process which corresponds to the lineage migration counts in our simulation study and we showed that the overall migration process is roughly estimated at any level of bias. In brief, we show on real datasets that singletons may be inferred as drivers of the migration process in an unparsimonious way by structured coalescent model approximations. This result supplements a previous study on the impact of the inclusion of few ghost deme sequences on the inference of migration rates (Ewing and Rodrigo 2006), however their impact remains unclear and deserves close consideration.

Interestingly, the posterior density of the structured coalescent model in BASTA is bimodal for the SARS-CoV-2 dataset. Its major mode corresponds to a past migration history close to CTMC-TRAVEL and our expectations of SARS-COV-2 spread at the start of the pandemic, whereas the minor mode corresponds to the non-parsimonious scenario. Such bimodality was not observed for MASCOT in the SARS-CoV-2 analysis. This difference in estimation is not unexpected since the two structured coalescent model approximations are different. However, it is not clear which characteristics of the two algorithms would lead to different behaviors. Another possibility relies on the choice of operators that determine how well the two approximations explore the parameter and tree space in which case MASCOT should lead to a bimodal posterior density on the long run.

### Practical implications for the analysis of empirical datasets

Computation time is an important consideration in real-life situations. CTMC is a fast algorithm that can handle many sequences while facing little convergence issues, which made it the predominant approach. For example, CTMC and its extensions have been extensively used during the SARS-CoV-2 pandemic (Candido et al. 2020; Dellicour, Durkin, et al. 2020; Lemey et al. 2020; Alteri et al. 2021; Butera et al. 2021; Dellicour et al. 2021; Kaleta et al. 2022; Perez et al. 2022). In general, researchers analyzed large datasets whose composition was corrected or reflected case counts (Candido et al. 2020; Lemey et al. 2020) or the number of hospitalizations per geographical location (Dellicour et al. 2021). According to our results, even though the pool of available sequences is not representative of the underlying transmission process, CTMC inference should be little impacted when using even sub-samples of the available sequences. However, we did not test sampling strategies based on case counts in our simulations.

With BASTA and MASCOT, computation time can become rapidly cumbersome and even impractical - also as a result of poor mixing of structured coalescent model parameters - when the number of sequences and locations increase. In such cases, these approaches are not able to discriminate which migration routes are the most important in the migration process leading to bimodal structured coalescent posterior densities, as observed for MASCOT on large samples of 500 sequences in the seven demes framework and for BASTA on the SARS-CoV-2 dataset. Repeating these problematic analyses with different starting values did not redeem these issues. Other studies have reported similar issues (Richardson et al. 2018). However, these problematic inferences can potentially be overcome by informing structured coalescent models with additional covariate data on viral population size dynamics. Indeed, as a result of adding such data, MASCOT-GLM not only outperformed the other approaches at estimating spatiotemporal parameters but also displayed improved mixing as expected with GLM approaches which improves the computational burden. However, such improvements depend on the availability and informativeness of the case count data used, notably on the early viral population size dynamics. This is illustrated in our analysis of the SARS-CoV-2 data for which the addition of WHO data led to improved chain mixing and past migration inference compared to the Our World in Data data, knowing that the dynamics are rather similar in the two datasets but they go back to January, 4th 2020 for the WHO data and to January, 23rd 2020 for the Our World in Data data.

### Limitations

We acknowledge several limitations of our study. First, BASTA and MASCOT are expected to perform better on even samples, a condition that we did not directly test. In the representative (uniform) samples, location frequencies inform CTMC and thus it would be expected to be favored over BASTA and MASCOT. Still, we show that MASCOT and BASTA perform better on even (region and region+year) samples in the three demes framework even if they are derived from biased large biobanks. This result suggests that BASTA and MASCOT perform better on even samples with no ghost demes. Second, our sub-sampling procedure in the simulation analysis could leave some locations unsampled, which can be considered as an extreme case of sampling bias. While this happened in only a few highly biased samples in the three demes framework, it is very common in the seven demes framework even in absence of sampling bias. It is difficult to determine whether the poor performance of MASCOT and BASTA in absence of bias in the seven demes framework compared to the three demes framework is due to ghost demes or is simply due to the higher number of locations. Additionally, we cannot rule out that the effects of sampling bias we observe are due to unsampled locations/unspecified ghost demes rather than unrepresentative sampling. We did not include unsampled locations as ghost demes in such conditions. However, this is unlikely to improve migration rate estimation (Ewing and Rodrigo 2006). Third, the impact of sampling bias certainly depends on the underlying overall migration rate as shown by (Liu et al.), an impact that we did not investigate here.

Another limitation concerns the incorporation of epidemiological data in phylogeographic models. Here, deme sizes in MASCOT-GLM are informed by case count data but this kind of data may not be readily available (Grubaugh et al. 2019) and is known to be often biased due to varying testing effort and strategy, as well as differential testing behaviors by age (Buckee et al. 2021). It is difficult to predict how MASCOT-GLM would perform if parameterized with biased case counts, a case that we did not address in our simulations. The comparison between the WHO and WID cases data, however, suggests that biased coverage of the true case load could bias such inference. If case count data are not reliable, one could use hospitalization data instead (Dellicour et al. 2021). Further, a similar approach is available under the CTMC framework but we did not test it here. This framework consists in modeling the migration process with CTMC and the overall population dynamics with the GLM extension (Gill et al. 2016) of the skygrid coalescent model (Gill et al. 2013). In this extension, case count over all locations could be used as a predictor of the viral population size over time. Yet, such an approach assumes a panmictic population and remains rare.

Finally, it is difficult to generalize our results in regards to the number of demes. Our choice of the number of locations was influenced by the RABV scenario in Morocco. While a scenario with three demes was doable, the one with seven demes turned out to be difficult to analyze, notably due to computational burden (Supplementary Materials). More research and development is needed for datasets with a large number of locations (> 15) and it currently seems unlikely that such analyses are possible at all with BASTA and MASCOT.

### Perspectives

In conclusion, sampling bias can be tackled at different levels of data generation and analysis in phylogeographic analyses: sample constitution, inference model choice, and data integration (e.g. through an integrated GLM). Other studies also assess the impact of sampling bias in *post hoc* analyses (Chaillon et al. 2020; Vrancken et al. 2020) or explicitly model sampling patterns (Guindon and De Maio 2021). Although the exploration of the impact of sampling bias has increased over the recent years and more robust methodologies have been developed, many aspects remain unclear, among which the impact of unsampled locations, biased epidemiological data incorporation, or the relative performances on even versus representative samples. Whenever possible, we would advise to opt for an even sampling strategy across geographical locations, compare the inferences of the different approaches or compare the inferences over multiple sub-samples when analyzing real datasets. These considerations are all the more important in a world of ever-growing genome sequence generation and concern not only human viral diseases but also zoonoses and epizooties.

## Material and Methods

### Simulation study

#### Simulation of viral transmission chains using a metapopulation model

In order to address the impact of spatial sampling bias on discrete phylogeographic inference, we performed a detailed simulation study. Sampling bias concerns all diseases, but it is even more challenging to address in the context of zoonotic diseases for which most of the transmission process is unobserved. We grounded our study in the context of dog rabies in North Africa where transmission processes are relatively well-documented. It was notably shown that rabies transmission relies on human movement over long distances. We simulated rabies epidemics in dog populations according to realistic scenarios using a stochastic, discrete-time and spatially-explicit model implemented in R using the Rccp package (Eddelbuettel and Balamuta 2018). We divided the Moroccan dog population into three or seven subpopulations corresponding to arbitrary regions (see the section below on the parametrization of the mobility matrix, Figure S22). We divided each subpopulation into three compartments: susceptible, exposed, and infectious individuals (Figure 1A). At each discrete time step, we drew newborns and dead individuals in the susceptible compartment from Poisson distributions with respective means the birth rate *b* and the death rate *d*. We defined the force of infection *Λ*_*i,t*_, i.e. the per-capita rate of infection of susceptible individuals in region *i* on day *t*, as:

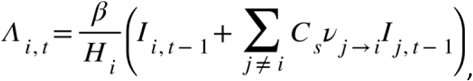

where *β* is the transmission rate of rabies scaled by *H*_*i*_, i.e. the human population size in region *i, ν* _*j* → *i*_ is the per-capita mobility rate of individuals moving from region *j* to region *i*, ***I***_*i, t* − 1_ is the number of infectious individuals in region *i* on day *t-1*, and ***C***_*s*_ is a scale factor (see below for more information). Exhaustive dog census data were not available and it is well known that human-mediated movement plays a major role in the spread of rabies in North Africa (Talbi et al. 2010; Dellicour et al. 2017), thus we assumed that dog populations were proportional to human populations (Table S2). We scaled the rabies transmission rate by population size to ensure that the force of infection is density-independent as previously documented on rabies (Morters et al. 2013). We used the scale factor *C*_*s*_ to monitor the proportion of inter-region infections. Its value was arbitrarily chosen so that 1% of infection events occurred between regions, and the basic reproduction ratio is approximately equal to 1.05 within and between regions. At each time step, we drew the number of newly exposed individuals in each region from Poisson distributions with a mean specified by the number of susceptible individuals in region *i* on day *t-1* (*S*_*i, t* − 1_) multiplied by the force of infection in region *i* on day t (*Λ*_*i, t*_). Once an individual *e*_*j, t*_ entered the exposed compartment, it was uniquely identified. The location of its infector was drawn from a multinomial distribution with the following probabilities:

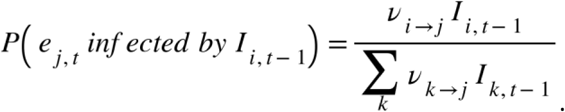

Once the location of the infector was drawn, the ID of the infector was randomly sampled from the set of infectors present in the location. All infectious individuals in each region had the same probability of infection. The incubation period of exposed individuals was drawn from a gamma distribution with shape 2 and rate 11.055 (Hampson et al. 2009) and its infectious period was drawn from a discretized gamma distribution adapted from Hampson et al. (Hampson et al. 2009) so that it could not exceed 15 days (World Health Organization (WHO) 2018). Finally, the life span was drawn from an exponential distribution with rate *d*. If natural death occurred before the end of the incubation or infectious periods, the individual was removed prematurely. Otherwise, the individual went through the exposed and infectious compartment before dying from rabies (Table S2).

We initiated all simulations with the introduction of a single index case in Region 3 (Figure S22). According to Darkaoui et al., there are on average 400 confirmed animal cases per year in Morocco (Darkaoui et al. 2017) which is certainly an underestimation (Broban et al. 2018). We assumed a 20% reporting rate of dog cases in Morocco (Taylor et al. 2017), and thus retained epidemics with at least 60,000 cases over a 30-year period (Figure 1C). We analyzed the results for 50 simulations.

#### Parametrization of the between-region mobility matrix

To avoid computational difficulties and over-parameterization of the different discrete phylogeographic models, we aggregated the fifteen official Moroccan regions retrieved from the GADM dataset (http://www.gadm.org) into three or seven locations (in two simulated scenarios, respectively) that are based on human demographics and ecological features (Figure S22). Dog mobility was defined across locations by fitting a radiation model to a raster of human population distribution (WorldPop) using the R package movement (Golding et al. 2015). In the radiation model, commuting is determined by the job seeking behavior modeled as an absorption and radiation process (Simini et al. 2012). The average commuting flux *T*_*i, j*_ from location *i* to location *j* with population *m*_*i*_ and *n*_*j*_, respectively is:

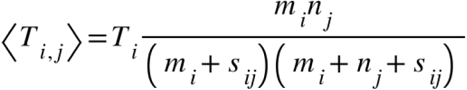

with *s*_*i, j*_ the total population in the circle of radius *r*_*i, j*_. centred at *i* (excluding the source and destination population).

We used a model of human mobility as it has been shown that humans play a major role in dog rabies spread and maintenance in North Africa, especially across long distances (Talbi et al. 2010; Dellicour et al. 2017). We preferred the radiation model over the gravity model for two reasons: the radiation model has been shown to outcompete the gravity model at local and large scales (Simini et al. 2012), and it presents the advantage of having no free parameter(s). In our study, we inferred the average daily number of commuters between raster cells of 20 km with more than 1,000 inhabitants per km^2^. The size of the cells corresponds approximately to the municipality level, and the density threshold corresponds to the urban density in Morocco. The number of commuters was then aggregated at the location level.

#### Evolutionary model of RABV genomes associated with cases

Simulation studies that analyze the accuracy of phylogeographical techniques often use the inference model as the simulation model (De Maio et al. 2015; Müller et al. 2017; Kalkauskas et al. 2021). Here, we took an epidemiological perspective by simulating rabies epidemics using a metapopulation model and by inferring the spatiotemporal history of rabies from RABV sequences and not from phylogenetic trees. After simulating rabies epidemics as described above, RABV genomes associated with each case were simulated according to the HKY model (Hasegawa et al. 1985). We simulated in R sequence evolution forwards-in-time along the transmission chains which were used in the same way as a phylogeny. We opted for a simple evolutionary process in which selection, gene partition, and site heterogeneity were not considered. Parameter values are listed in Table S2. The genome of the index case is a real canine rabies genome of 13 kb length isolated in Morocco in 2013 (GenBank Accession Number KF155001.1) (Marston et al. 2013).

#### Sampling schemes of viral sequences

The aim of the study is to determine the impact of sampling bias on phylogeographic inference and how alternative sampling schemes may mitigate the effects of such sampling bias. To address the former issue, we sampled either uniformly (uniform) or with a sampling bias favoring viral sequences from highly populated locations (Regions 3 and 4). In the latter scenario, sequences from Regions 1, 2, 5, 6, and 7 had a weight equal to one, whereas Regions 3 and 4 had a weight equal to 2.5, 5, 10, 20 or and 50 (biased-2.5, biased-5, biased-10, biased-20, and biased-50, respectively, Figure 1D). To mitigate the potential effects of sampling bias, we tested a different setup reproducing a surveillance system. In this setup, a biobank of 5,000 sequences were drawn from each epidemic with a weight of one for Regions 1, 2, 5, 6, and 7, and a weight of 10 or 20 for Region 3 and 4. Subsets of sequences were sampled from the biobank either uniformly (uniform surv.), by maximizing the spatial coverage (max per region), or by maximizing the spatiotemporal coverage (max per region and per year). For all sampling schemes, a large sample of 500 sequences and a nested sample of 150 sequences were drawn over the entire epidemic except for the first year, as we assumed that the spread of the virus would remain undetected at the start of the epidemic as observed in other settings (Townsend et al. 2013).

#### Discrete phylogeographic analysis in BEAST

##### Generation of BEAST XML files and phylogeography inference set up

Tailored XML template files for the BASTA and MASCOT structured coalescent models, as well as for the discrete trait analysis (CTMC) model, were edited using the lxml Python package to add sequence alignments along with their metadata. Bayesian phylogeographic analyses were performed using BEAST v1.10.5 (Suchard et al. 2018) for the CTMC model (Lemey et al. 2009), and BEAST v2.6.4 (Bouckaert et al. 2019) for MASCOT v2.2.1 (Müller et al. 2018) and BASTA v3.0.1 (De Maio et al. 2015), making use of the BEAGLE library v3.1.1 (Ayres et al. 2012). We assumed an HKY substitution model with a strict molecular clock. Population dynamics in the CTMC model followed a constant population size prior. We chose this prior since the model of population dynamics is not expected to impact migration history inference and the constant population size model is often chosen for the analysis of endemic diseases. For the BASTA and MASCOT structured coalescent models, all demes were set to have equal size due to numerical issues leading to a computation time of over 70 hours per million iterations (data not shown). For both models, asymmetric migration matrices were inferred and Bayesian stochastic search variable selection (BSSVS) was used to avoid over-parametrization. The detailed list of prior distributions is available in Table S3 for each inference framework.

If deme sizes are set to be equal in the structured coalescent model but the actual population dynamics vary through time, the model tends to explain population dynamics by migration dynamics. In our simulations, the incidence changed dramatically over time and location (Figure 1C), thus the inference by the structured coalescent model is expected to improve when accounting for time-varying population dynamics. To test this hypothesis, we used monthly incidence data from our simulations as a predictor of the deme sizes by using a GLM in MASCOT (Müller et al. 2019). We tested this alternative parametrization (MASCOT-GLM) in the following conditions: uniform, biased-2.5, biased-5, biased-10, biased-20, biased-50, uniform surv. 10, and uniform surv. 20.

These different BEAST analyses were run for at least 20 and 40 million steps, and sampled every 2,000 and 4,000 steps for small and large alignments, respectively. In total, 8,800 XML files were run for this study, for a total of an estimated 1,500 hours of computation on multi-core CPUs across different computing infrastructures (Table S4).

##### Analysis of phylogeographic inference output

For each BEAST analysis, adequate mixing was assessed based on the effective sample size (ESS) values of the continuous parameters. We calculated ESS values using a Python function adapted from Tracer v1.7.2 (Rambaut et al. 2018). When at least one continuous parameter had an ESS value below 200, chains were resumed to reach at most 120 million iterations. Analyses that exhibited ESS values lower than 200 at this point were discarded (Tables S5 and S6). Due to the higher computational burden of BASTA, the ESS cut-off was reduced to 100. We discarded a 10% burn-in in the selected chains. The combined posterior tree distributions were summarized into MCC trees using TreeAnnotator for BASTA and the CTMC, and the Python library dendropy (Sukumaran and Holder 2010) for MASCOT and MASCOT-GLM. Summary statistics, ESS values, Bayes factors (BFs) on migration rates (Lemey et al. 2009), root state probabilities, dates of lineage introduction, and lineage migration counts were calculated in Python before plotting the results in R using the ggplot2 package (Wickham 2016).

##### Performance analysis

To assess the accuracy of the phylogenetic reconstruction, the time to the most recent common ancestor (TMRCA) of every pair of sampled tips was computed on both the MCC tree and the simulated transmission chain, and these outcomes were subsequently compared using the Pearson correlation coefficient (Figure 1A). In addition, we evaluated the impact of sampling bias and alternative sampling strategies on the estimation of the total migration counts, lineage migration counts, and dates of first lineage introduction into each sampled location using five metrics:

- Kendall’s tau correlation: a rank-correlation measure that is less sensitive to outliers compared to Pearson’s correlation coefficient

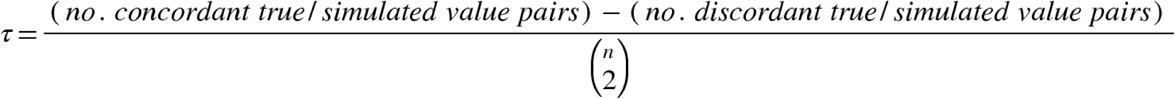
- calibration

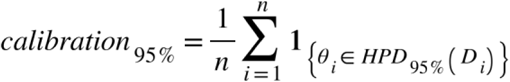
- mean relative bias

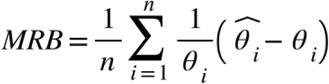
- mean relative 95% highest posterior density (HPD) width

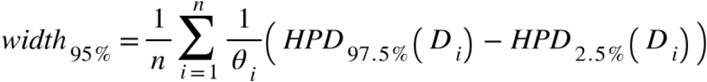
- weighted interval score (WIS): a generalization of the absolute error accounting for estimation uncertainty. We present the formula of the WIS and refer to the original article for further details, notably on the interval score (Bracher et al. 2021).

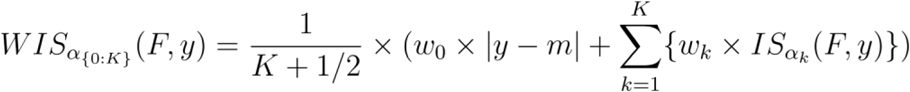

We denote *θ*_*i*_ the true value of the parameter, *D*_*i*_ the parameter posterior distribution, 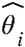 the median estimate, *HPD*_95%_the 95% HPD, *K* the number of prediction intervals included in the calculation of the WIS, *y* the observed outcome by forecast *F, m* the predictive median on the (1 − *α*_*k*_) × 100% prediction interval, 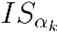 the interval score on the (1 − *α*_*k*_) × 100% prediction interval and *w*_*k*_ its weight. The mean relative bias and the mean relative 95% HPD width are defined when the true value is not zero. However, the total migration counts and the lineage migration counts for some pairs of locations can be null in our simulations whereas the algorithms infer a non-null median. These cases were not considered in the calculation of the mean relative bias and the mean relative 95% HPD width. We reported their numbers in the caption of the corresponding figures.

### Data analysis

#### RABV expansion in the Philippines

We extended our comparative analysis of the CTMC, BASTA, and MASCOT by analyzing a set of RABV genetic sequences using the three approaches. In total, 233 sequences corresponding to the RABV glycoprotein gene were sampled in the Philippines from 2004 and 2010 (Saito et al. 2013). In the original discrete phylogeographic analysis, the authors studied viral spread across 11 out of the 17 Philippines regions and showed that the genetic diversity was highly spatially-structured, notably at the island level (Tohma et al. 2014). Here, we evaluated spread across the six sampled islands (Luzon, Catanduanes, Oriental Mindoro, Cebu, Negros Oriental, and Mindanao) to compare the reconstructions on a highly structured dataset and limit the number of demes that considerably slow down BASTA and MASCOT. We assumed an HKY nucleotide substitution model with an among-site rate heterogeneity modeled by a discretized gamma distribution (Yang 1994), and an uncorrelated relaxed molecular clock with an underlying lognormal distribution (Drummond et al. 2006). For the CTMC, we assumed a constant size coalescent model for the viral demographics as in the original analysis. For MASCOT and BASTA, current implementations assume a constant population size model for the viral demographics within demes. A detailed description of the priors is reported in Table S7. For each algorithm, we combined three post-burnin independent chains of 50 million iterations each.

#### The early dynamics of SARS-CoV-2 worldwide spread

Tracking viral disease spread in animal populations faces many challenges, and to our knowledge, no reliable incidence data are available for zoonoses such as rabies. In this context, MASCOT-GLM cannot readily be used. We analyzed the early worldwide spread of SARS-CoV-2 to compare the inferences of the CTMC, BASTA, MASCOT, and MASCOT-GLM. Lemey et al. analyzed this dataset to characterize SARS-CoV-2 spread across 44 location states by incorporating individual travel histories of sampled individuals to help correct for sampling bias and unsampled locations (Lemey et al. 2020). By using the carefully obtained results of (Lemey et al. 2020) as a reference, we can evaluate how the four algorithms are impacted by sampling bias.

The dataset comprises 282 SARS-CoV-2 genomic sequences sampled in the five continents from December 24th, 2019 to March 4th, 2020. We assumed an HKY nucleotide substitution model with a proportion of invariant sites, an among-site rate heterogeneity modeled by a discretized gamma distribution, and a strict molecular clock. For the CTMC, we assumed an exponential growth model for the viral demographics. MASCOT and BASTA assume a constant size model for the viral demes demographics. Contrary to the original study, we analyzed migration between six discrete locations: Africa, Americas, Asia, China, Europe, and Oceania. For MASCOT-GLM, we used the daily number of confirmed cases at the continent level from Our World In Data (Ritchie et al. 2020), or from the (World Health Organization (WHO)) as a predictor of the deme sizes. The former is referred to as MASCOT-WID, and the latter as MASCOT-WHO. We smoothed the number of new confirmed cases using a seven-day moving average. The detailed description of the priors is reported in Table S8. We combined three post-burnin independent chains of 50 or 100 million iterations for each inference.

## Data Availability

All scripts used to simulate epidemics and perform the analyses presented in this article are available at https://github.com/mlayan/Sampling_bias including examples of output files. Output files which are not in the GitHub repository are available upon reasonable request to the authors. Supplementary data are available at *Molecular Biology and Evolution* online.

## Acknowledgments

We would like to thank Dr Alessio Andronico for his help on the implementation of the rabies simulation model in Rcpp. SD acknowledges support from the *Fonds National de la Recherche Scientifique* (F.R.S.-FNRS, Belgium; grant n°F.4515.22) and from the European Union Horizon 2020 project MOOD (grant agreement n°874850). SD and GB acknowledge support from the Research Foundation - Flanders (*Fonds voor Wetenschappelijk Onderzoek - Vlaanderen*, FWO, Belgium; grant n° G098321N). GB also acknowledges support from the Internal Funds KU Leuven (Grant No. C14/18/094) and the Research Foundation - Flanders (*Fonds voor Wetenschappelijk Onderzoek - Vlaanderen*, FWO, Belgium; G0E1420N). SC acknowledges financial support from the Laboratoire d’Excellence Integrative Biology of Emerging Infectious Diseases program (grant ANR-10-LABX-62-IBEID) and the INCEPTION project (grant PIA/ANR-16-CONV-0005).

## References

Alteri C, Cento V, Piralla A, Costabile V, Tallarita M, Colagrossi L, Renica S, Giardina F, Novazzi F, Gaiarsa S, et al. 2021. Genomic epidemiology of SARS-CoV-2 reveals multiple lineages and early spread of SARS-CoV-2 infections in Lombardy, Italy. Nat. Commun. 12:1–13.

Ayres DL, Darling A, Zwickl DJ, Beerli P, Holder MT, Lewis PO, Huelsenbeck JP, Ronquist F, Swofford DL, Cummings MP, et al. 2012. BEAGLE: An application programming interface and high-performance computing library for statistical phylogenetics. Syst. Biol. 61:170–173.

Beerli P. 2004. Effect of unsampled populations on the estimation of population sizes and migration rates between sampled populations. Mol. Ecol. 13:827–836.

Bouckaert R, Vaughan TG, Barido-Sottan J, Duchêne S, Fourment M, Gavryushkina A, Heled J, Jones G, Kühnert D, Maio N De, et al. 2019. BEAST 2.5 : An Advanced Software Platform for Bayesian Evolutionary Analysis. PLoS Comput. Biol. 15:e1006650.

Bracher J, Ray EL, Gneiting T, Reich NG. 2021. Evaluating epidemic forecasts in an interval format.Pitzer VE, editor. PLOS Comput. Biol. 17:e1008618.

Broban A, Tejiokem MC, Tiembré I, Druelles S, L’Azou M. 2018. Bolstering human rabies surveillance in Africa is crucial to eliminating canine-mediated rabies.Knobel D, editor. PLoS Negl. Trop. Dis. 12:e0006367.

Brynildsrud OB, Pepperell CS, Suffys P, Grandjean L, Monteserin J, Debech N, Bohlin J, Alfsnes K, Pettersson JOH, Kirkeleite I, et al. 2018. Global expansion of Mycobacterium tuberculosis lineage 4 shaped by colonial migration and local adaptation. Sci. Adv. 4:5869–5886.

Buckee C, Noor A, Sattenspiel L. 2021. Thinking clearly about social aspects of infectious disease transmission. Nature 595:205–213.

Butera Y, Mukantwari E, Artesi M, Umuringa J d’arc, O’Toole ÁN, Hill V, Rooke S, Hong SL, Dellicour S, Majyambere O, et al. 2021. Genomic sequencing of SARS-CoV-2 in Rwanda reveals the importance of incoming travelers on lineage diversity. Nat. Commun. 12:5705.

Candido DS, Claro IM, de Jesus JG, Souza WM, Moreira FRR, Dellicour S, Mellan TA, du Plessis L, Pereira RHM, Sales FCS, et al. 2020. Evolution and epidemic spread of SARS-CoV-2 in Brazil. Science (80-.). 369:1255–1260.

Chaillon A, Gianella S, Dellicour S, Rawlings SA, Schlub TE, de Oliveira MF, Ignacio C, Porrachia M, Vrancken B, Smith DM. 2020. HIV persists throughout deep tissues with repopulation from multiple anatomical sources. J. Clin. Invest. 130:1699–1712.

Darkaoui S, Cliquet F, Wasniewski M, Robardet E, Aboulfidaa N, Bouslikhane M, Fassi-Fihri O. 2017. A Century Spent Combating Rabies in Morocco (1911–2015): How Much Longer? Front. Vet. Sci. 4:1–16.

Dellicour S, Durkin K, Hong SL, Vanmechelen B, Martí-Carreras J, Gill MS, Meex C, Bontems S, André E, Gilbert M, et al. 2020. A phylodynamic workflow to rapidly gain insights into the dispersal history and dynamics of SARS-CoV-2 lineages. Mol. Biol. Evol.:1–6.

Dellicour S, Hong SL, Vrancken B, Chaillon A, Gill MS, Maurano MT, Ramaswami S, Zappile P, Marier C, Harkins GW, et al. 2021. Dispersal dynamics of SARS-CoV-2 lineages during the first epidemic wave in New York City. Lauring AS, editor. PLOS Pathog. 17:e1009571.

Dellicour S, Lequime S, Vrancken B, Gill MS, Bastide P, Gangavarapu K, Matteson NL, Tan Y, du Plessis L, Fisher AA, et al. 2020. Epidemiological hypothesis testing using a phylogeographic and phylodynamic framework. Nat. Commun. 11:5620.

Dellicour S, Rose R, Faria NR, Vieira LFP, Bourhy H, Gilbert M, Lemey P, Pybus OG. 2017. Using Viral Gene Sequences to Compare and Explain the Heterogeneous Spatial Dynamics of Virus Epidemics. Mol. Biol. Evol. 34:2563–2571.

Drummond AJ, Ho SYW, Phillips MJ, Rambaut A. 2006. Relaxed phylogenetics and dating with confidence. PLoS Biol. 4:699–710.

Dudas G, Carvalho LM, Bedford T, Tatem AJ, Baele G, Faria NR, Park DJ, Ladner JT, Arias A, Asogun D, et al. 2017. Virus genomes reveal factors that spread and sustained the Ebola epidemic. Nature 544:309–315.

Dudas G, Carvalho LM, Rambaut A, Bedford T. 2018. MERS-CoV spillover at the camel-human interface. Elife 7.

Eddelbuettel D, Balamuta JJ. 2018. Extending R with C++: A Brief Introduction to Rcpp. Am. Stat. 72:28–36.

Ewing G, Rodrigo A. 2006. Estimating Population Parameters using the Structured Serial Coalescent with Bayesian MCMC Inference when some Demes are Hidden. Evol. Bioinforma. 2:117693430600200.

Faria NR, Quick J, Claro IM, Thézé J, de Jesus JG, Giovanetti M, Kraemer MUG, Hill SC, Black A, da Costa AC, et al. 2017. Establishment and cryptic transmission of Zika virus in Brazil and the Americas. Nature 546:406–410.

Faria NR, Vidal N, Lourenco J, Raghwani J, Sigaloff KCE, Tatem AJ, Van De Vijver DAM, Pineda-Peña AC, Rose R, Wallis CL, et al. 2019. Distinct rates and patterns of spread of the major HIV-1 subtypes in Central and East Africa. PLoS Pathog. 15:1–23.

Frost SDW, Pybus OG, Gog JR, Viboud C, Bonhoeffer S, Bedford T. 2015. Eight challenges in phylodynamic inference. Epidemics 10:88–92.

Gill MS, Lemey P, Bennett SN, Biek R, Suchard MA. 2016. Understanding Past Population Dynamics: Bayesian Coalescent-Based Modeling with Covariates. Syst. Biol. 65:1041–1056.

Gill MS, Lemey P, Faria NR, Rambaut A, Shapiro B, Suchard MA. 2013. Improving bayesian population dynamics inference: A coalescent-based model for multiple loci. Mol. Biol. Evol. 30:713–724.

Golding N, Schofield A, Kraemer MUG. 2015. Movement: Functions for the analysis of movement data in disease modelling and mapping. R Packag. version 0.2.

Grubaugh ND, Saraf S, Gangavarapu K, Watts A, Tan AL, Oidtman RJ, Ladner JT, Oliveira G, Matteson NL, Kraemer MUG, et al. 2019. Travel Surveillance and Genomics Uncover a Hidden Zika Outbreak during the Waning Epidemic. Cell 178:1057–1071.e11.

Guindon S, De Maio N. 2021. Accounting for spatial sampling patterns in Bayesian phylogeography. Proc. Natl. Acad. Sci. 118:e2105273118.

Hampson K, Dushoff J, Cleaveland S, Haydon DT, Kaare M, Packer C, Dobson A. 2009. Transmission dynamics and prospects for the elimination of canine Rabies. PLoS Biol. 7:0462–0471.

Hasegawa M, Kishino H, Yano T aki. 1985. Dating of the human-ape splitting by a molecular clock of mitochondrial DNA. J. Mol. Evol. 22:160–174.

He W-T, Bollen N, Xu Y, Zhao J, Dellicour S, Yan Z, Gong W, Zhang C, Zhang L, Lu M, et al. 2022. Phylogeography Reveals Association between Swine Trade and the Spread of Porcine Epidemic Diarrhea Virus in China and across the World.Barlow M, editor. Mol. Biol. Evol. 39.

Hodcroft EB, De Maio N, Lanfear R, MacCannell DR, Minh BQ, Schmidt HA, Stamatakis A, Goldman N, Dessimoz C. 2021. Want to track pandemic variants faster? Fix the bioinformatics bottleneck. Nature 591:30–33.

Hong SL, Lemey P, Suchard MA, Baele G. 2021. Bayesian Phylogeographic Analysis Incorporating Predictors and Individual Travel Histories in BEAST. Curr. Protoc. 1:1–16.

Kaleta T, Kern L, Hong SL, Hölzer M, Kochs G, Beer J, Schnepf D, Schwemmle M, Bollen N, Kolb P, et al. 2022. Antibody escape and global spread of SARS-CoV-2 lineage A.27. Nat. Commun. 13:1152.

Kalkauskas A, Perron U, Sun Y, Goldman N, Baele G, Guindon S, De Maio N. 2021. Sampling bias and model choice in continuous phylogeography: Getting lost on a random walk.Kosakovsky Pond SL, editor. PLOS Comput. Biol. 17:e1008561.

Lemey P, Hong SL, Hill V, Baele G, Poletto C, Colizza V, O’Toole Á, McCrone JT, Andersen KG, Worobey M, et al. 2020. Accommodating individual travel history and unsampled diversity in Bayesian phylogeographic inference of SARS-CoV-2. Nat. Commun. 11:5110.

Lemey P, Rambaut A, Bedford T, Faria N, Bielejec F, Baele G, Russell CA, Smith DJ, Pybus OG, Brockmann D, et al. 2014. Unifying Viral Genetics and Human Transportation Data to Predict the Global Transmission Dynamics of Human Influenza H3N2. PLoS Pathog. 10.

Lemey P, Rambaut A, Drummond AJ, Suchard MA. 2009. Bayesian phylogeography finds its roots. PLoS Comput. Biol. 5.

Liu P, Song Y, Colijn C, Macpherson A. The impact of sampling bias on viral phylogeographic reconstruction. Available from: https://doi.org/10.1101/2022.05.12.22275024

Lu L, Sikkema RS, Velkers FC, Nieuwenhuijse DF, Fischer EAJ, Meijer PA, Bouwmeester-Vincken N, Rietveld A, Wegdam-Blans MCA, Tolsma P, et al. 2021. Adaptation, spread and transmission of SARS-CoV-2 in farmed minks and associated humans in the Netherlands. Nat. Commun. 12:6802.

Magee D, Scotch M. 2018. The effects of random taxa sampling schemes in Bayesian virus phylogeography. Infect. Genet. Evol. 64:225–230.

De Maio N, Wu CH, O’Reilly KM, Wilson D. 2015. New Routes to Phylogeography: A Bayesian Structured Coalescent Approximation. PLoS Genet. 11:1–22.

Marston DA, McElhinney LM, Ellis RJ, Horton DL, Wise EL, Leech SL, David D, Lamballerie X de, Fooks AR. 2013. Next generation sequencing of viral RNA genomes. BMC Genomics 14:444.

Mavian C, Paisie TK, Alam MT, Browne C, Beau De Rochars VM, Nembrini S, Cash MN, Nelson EJ, Azarian T, Ali A, et al. 2020. Toxigenic Vibrio cholerae evolution and establishment of reservoirs in aquatic ecosystems. Proc. Natl. Acad. Sci. 117:7897–7904.

Morel B, Barbera P, Czech L, Bettisworth B, Hübner L, Lutteropp S, Serdari D, Kostaki E-G, Mamais I, Kozlov AM, et al. 2021. Phylogenetic Analysis of SARS-CoV-2 Data Is Difficult.Malik H, editor. Mol. Biol. Evol. 38:1777–1791.

Morters MK, Restif O, Hampson K, Cleaveland S, Wood JLN, Conlan AJK. 2013. Evidence-based control of canine rabies: a critical review of population density reduction. Boots M, editor. J. Anim. Ecol. 82:6–14.

Müller NF, Dudas G, Stadler T. 2019. Inferring time-dependent migration and coalescence patterns from genetic sequence and predictor data in structured populations. Virus Evol. 5:1–10.

Müller NF, Rasmussen D, Stadler T. 2018. MASCOT: Parameter and state inference under the marginal structured coalescent approximation. Bioinformatics 34:3843–3848.

Müller NF, Rasmussen DA, Stadler T. 2017. The structured coalescent and its approximations. Mol. Biol. Evol. 34:2970–2981.

Müller NF, Wagner C, Frazar CD, Roychoudhury P, Lee J, Moncla LH, Pelle B, Richardson M, Ryke E, Xie H, et al. 2021. Viral genomes reveal patterns of the SARS-CoV-2 outbreak in Washington State. Sci. Transl. Med. 13:202.

Perez LJ, Orf GS, Berg MG, Rodgers MA, Meyer T V., Mohaimani A, Olivo A, Harris B, Mowerman I, Padane A, et al. 2022. The early SARS-CoV-2 epidemic in Senegal was driven by the local emergence of B.1.416 and the introduction of B.1.1.420 from Europe. Virus Evol. 8:1–12.

Pipes L, Wang H, Huelsenbeck JP, Nielsen R. 2021. Assessing Uncertainty in the Rooting of the SARS-CoV-2 Phylogeny.Malik H, editor. Mol. Biol. Evol. 38:1537–1543.

Rambaut A, Drummond AJ, Xie D, Baele G, Suchard MA. 2018. Posterior summarization in Bayesian phylogenetics using Tracer 1.7. Syst. Biol. 67:901–904.

Richardson EJ, Bacigalupe R, Harrison EM, Weinert LA, Lycett S, Vrieling M, Robb K, Hoskisson PA, Holden MTG, Feil EJ, et al. 2018. Gene exchange drives the ecological success of a multi-host bacterial pathogen. Nat. Ecol. Evol. 2:1468–1478.

Ritchie H, Mathieu E, Rodés-Guirao L, Appel C, Giattino C, Ortiz-Ospina E, Hasell J, Macdonald B, Beltekian D, Roser M. 2020. Coronavirus Pandemic (COVID-19). Our World Data.

Saito M, Oshitani H, Orbina JRC, Tohma K, de Guzman AS, Kamigaki T, Demetria CS, Manalo DL, Noguchi A, Inoue S, et al. 2013. Genetic Diversity and Geographic Distribution of Genetically Distinct Rabies Viruses in the Philippines.Rupprecht CE, editor. PLoS Negl. Trop. Dis. 7:e2144.

Simini F, González MC, Maritan A, Barabási AL. 2012. A universal model for mobility and migration patterns. Nature 484:96–100.

Suchard MA, Lemey P, Baele G, Ayres DL, Drummond AJ, Rambaut A. 2018. Bayesian phylogenetic and phylodynamic data integration using BEAST 1.10. Virus Evol. 4:1–5.

Sukumaran J, Holder MT. 2010. DendroPy: a Python library for phylogenetic computing. Bioinformatics 26:1569–1571.

Talbi C, Lemey P, Suchard MA, Abdelatif E, Elharrak M, Jalal N, Faouzi A, Echevarría JE, Morón SV, Rambaut A, et al. 2010. Phylodynamics and Human-mediated dispersal of a zoonotic virus. PLoS Pathog. 6.

Taylor LH, Hampson K, Fahrion A, Abela-Ridder B, Nel LH. 2017. Difficulties in estimating the human burden of canine rabies. Acta Trop. 165:133–140.

Tohma K, Saito M, Demetria CS, Manalo DL, Quiambao BP, Kamigaki T, Oshitani H. 2016. Molecular and mathematical modeling analyses of inter-island transmission of rabies into a previously rabies-free island in the Philippines. Infect. Genet. Evol. 38:22–28.

Tohma K, Saito M, Kamigaki T, Tuason LT, Demetria CS, Orbina JRC, Manalo DL, Miranda ME, Noguchi A, Inoue S, et al. 2014. Phylogeographic analysis of rabies viruses in the Philippines. Infect. Genet. Evol. 23:86–94.

Townsend SE, Sumantra IP, Pudjiatmoko, Bagus GN, Brum E, Cleaveland S, Crafter S, Dewi APM, Dharma DMN, Dushoff J, et al. 2013. Designing Programs for Eliminating Canine Rabies from Islands: Bali, Indonesia as a Case Study. PLoS Negl. Trop. Dis. 7.

Vaughan TG, Kühnert D, Popinga A, Welch D, Drummond AJ. 2014. Efficient Bayesian inference under the structured coalescent. Bioinformatics 30:2272–2279.

Vrancken B, Zhao B, Li X, Han X, Liu H, Zhao J, Zhong P, Lin Y, Zai J, Liu M, et al. 2020. Comparative Circulation Dynamics of the Five Main HIV Types in China.Silvestri G, editor. J. Virol. 94:683–703.

Wickham H. 2016. ggplot2: Elegant Graphics for Data Analysis. Springer-Verlag New York Available from: https://ggplot2.tidyverse.org

World Health Organization (WHO). Covid-19 cases and deaths by continent. Available from: https://portal.who.int/report/eios-covid19-counts/#display=Continents&nrow=2&ncol=3&arr=row&pg=1&labels=view_countries&sort=cur_case_who;desc&filter=&sidebar=-1&fv=

World Health Organization (WHO). 2018. WHO Expert Consultation on Rabies. Third report. Geneva WorldPop. WorldPop project. Available from: http://worldpop.org.uk/

Yang J, Müller NF, Bouckaert R, Xu B, Drummond AJ. 2019. Bayesian phylodynamics of avian influenza A virus H9N2 in Asia with time-dependent predictors of migration. PLOS Comput. Biol. 15:e1007189.

Yang Z. 1994. Maximum likelihood phylogenetic estimation from DNA sequences with variable rates over sites: Approximate methods. J. Mol. Evol. 39:306–314.

