## Supplementary Materials for "Impact and mitigation of sampling bias to determine viral spread: evaluating discrete phylogeography through CTMC modeling and structured coalescent model approximations"

#### Table of Contents

**Impact and mitigation of bias in the seven demes framework** **2**
Estimation of genetic parameters 2
Total migration counts 8
Lineage migration counts 9

**Impact and mitigation of bias in the three demes framework** **11**
Estimation of genetic parameters 11
Lineage migration counts 18
Total migration counts 19
Introduction dates 20
Root location 21

**RABV spread in the Philippines** **22**

**SARS-CoV-2 early spread** **23**

**Simulation framework of RABV epidemics** **32**

**Bayesian inference** **34**
Simulation study 34
Analysis of the rabies dataset 36
Analysis of the SARS-CoV-2 dataset 37

**References** **39**

### 1. Impact and mitigation of bias in the seven demes framework

#### a. Estimation of genetic parameters

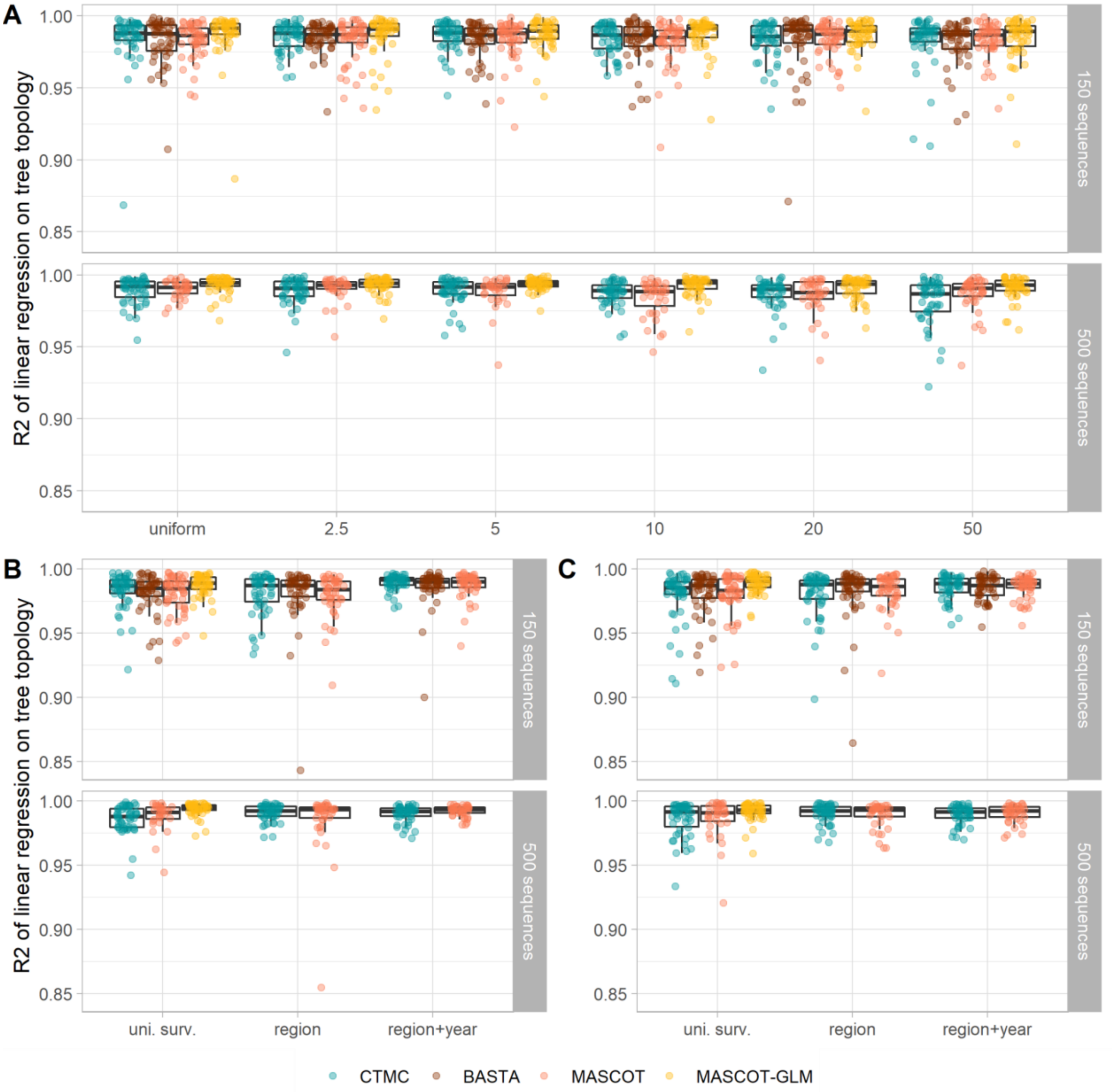


**Figure S1. Comparison of the simulated and estimated tree topologies for all sampling conditions and the four algorithms.** Pearson's determination coefficient of the pairwise divergence time between the simulated transmission chain and the MCC tree for the systematic bias (**A**), surveillance bias 10 (**B**), and surveillance bias 20 (**C**) sampling conditions.

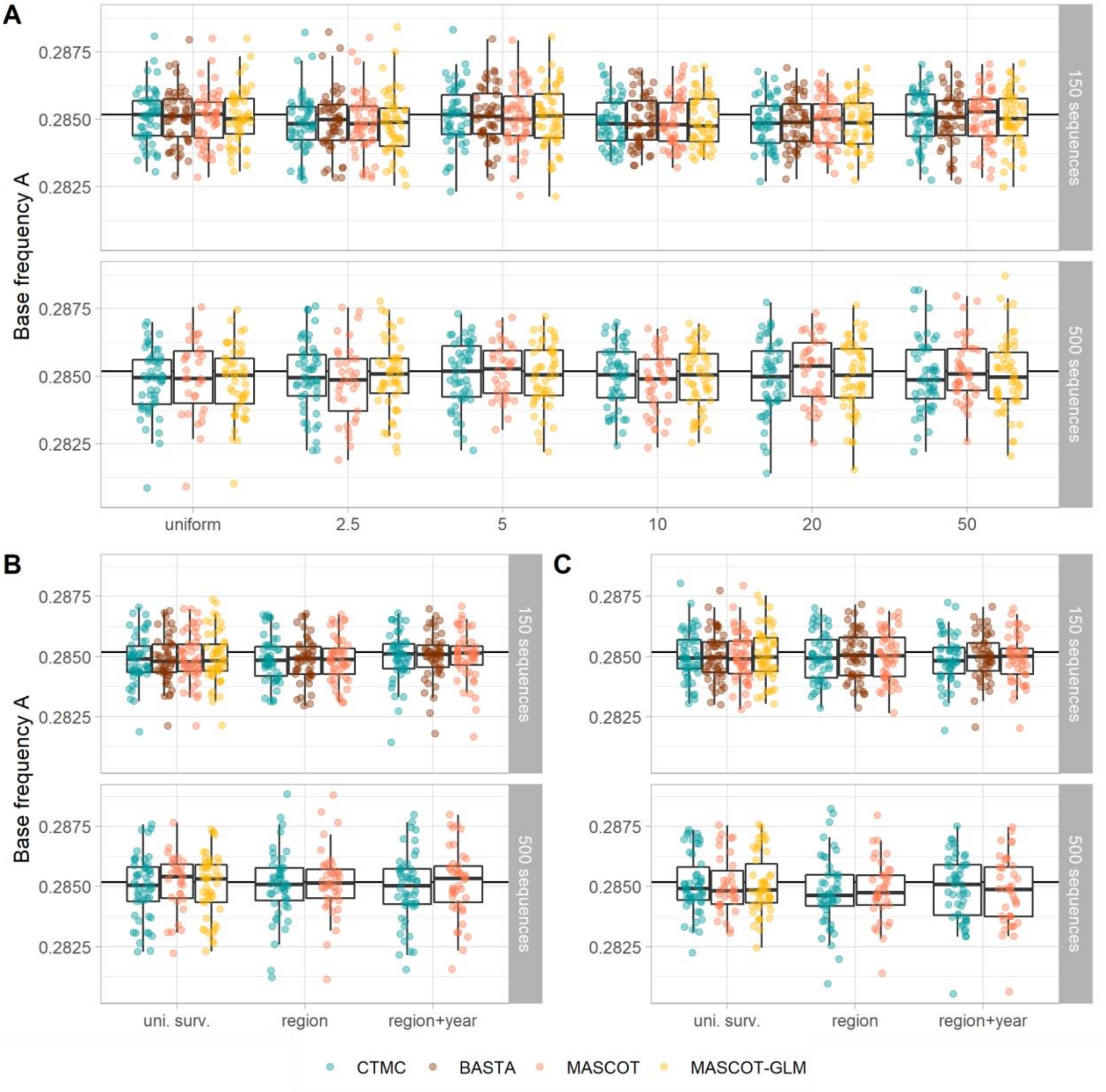


**Figure S2. Median estimates of base frequency A for all sampling conditions and the four algorithms.** Median estimate of the base frequency of A for the systematic bias (**A**), surveillance bias 10 (**B**), and surveillance bias 20 (**C**) sampling conditions. The true value of the parameter is represented as the horizontal black line.

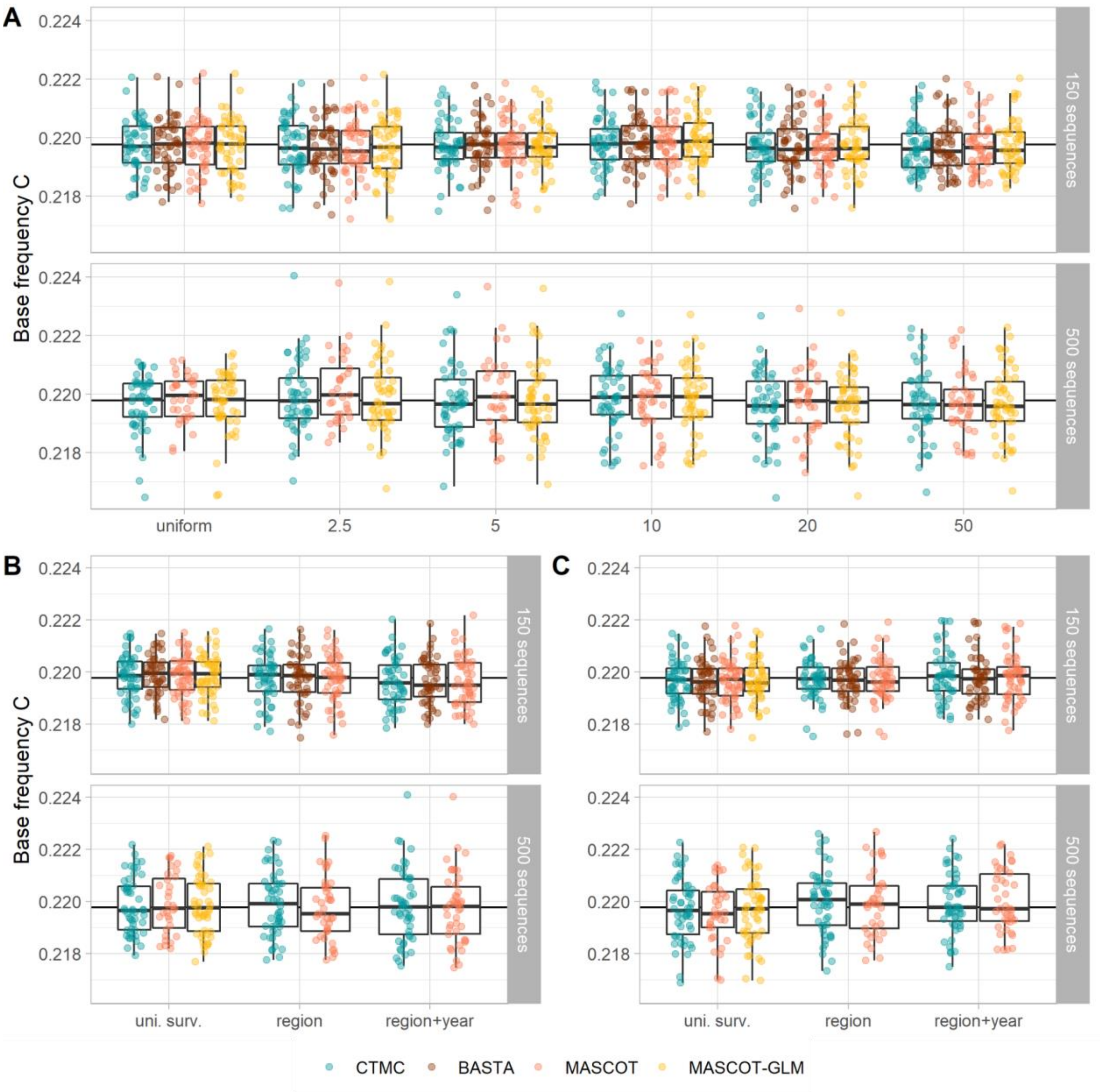


**Figure S3. Median estimates of base frequency C for all sampling conditions and the four algorithms.** Median estimate of the base frequency of C for the systematic bias (**A**), surveillance bias 10 (**B**), and surveillance bias 20 (**C**) sampling conditions. The true value of the parameter is represented as the horizontal black line.

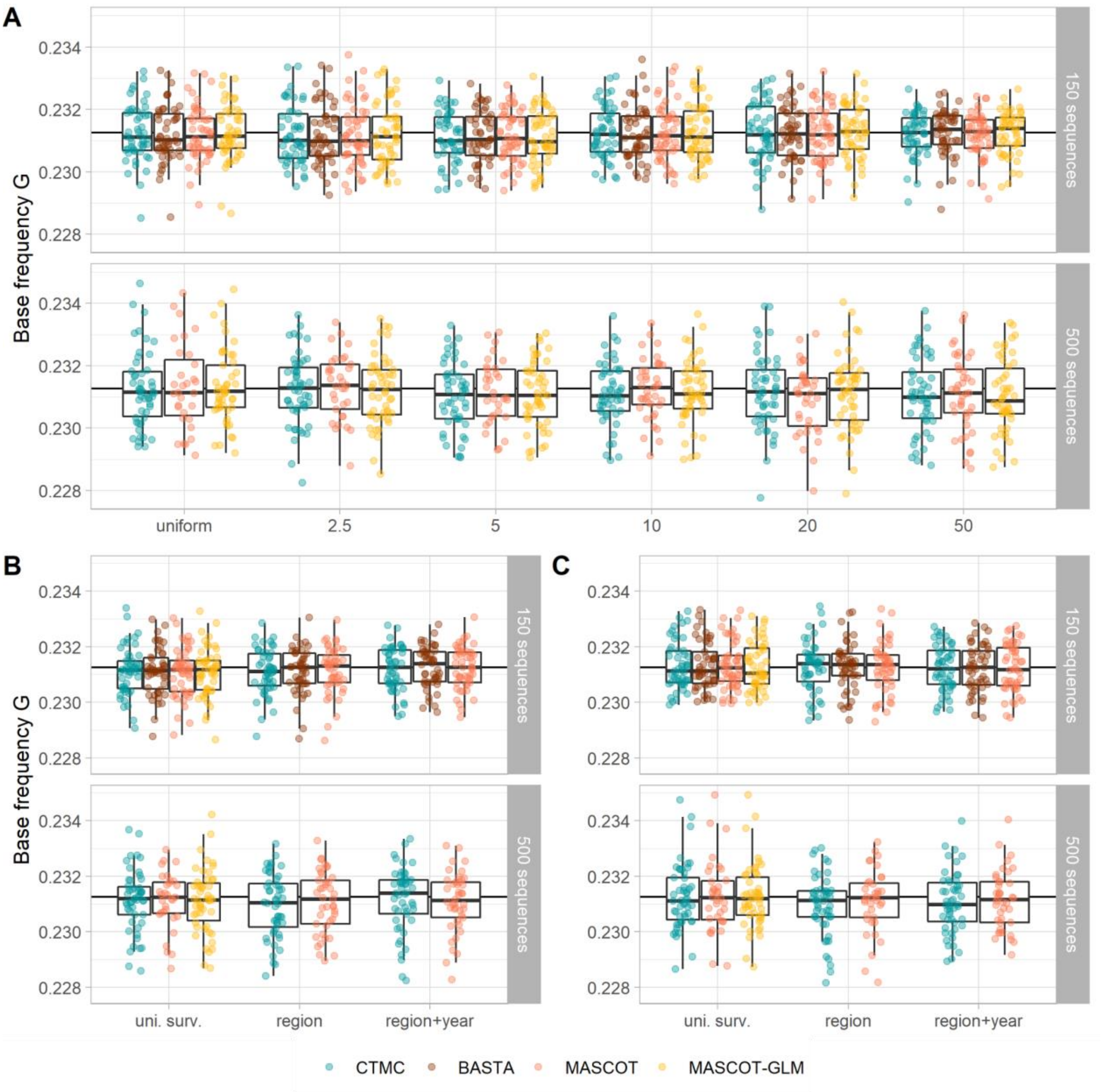


**Figure S4. Median estimates of base frequency G for all sampling conditions and the four algorithms.** Median estimate of the base frequency of G for the systematic bias (**A**), surveillance bias 10 (**B**), and surveillance bias 20 (**C**) sampling conditions. The true value of the parameter is represented as the horizontal black line.

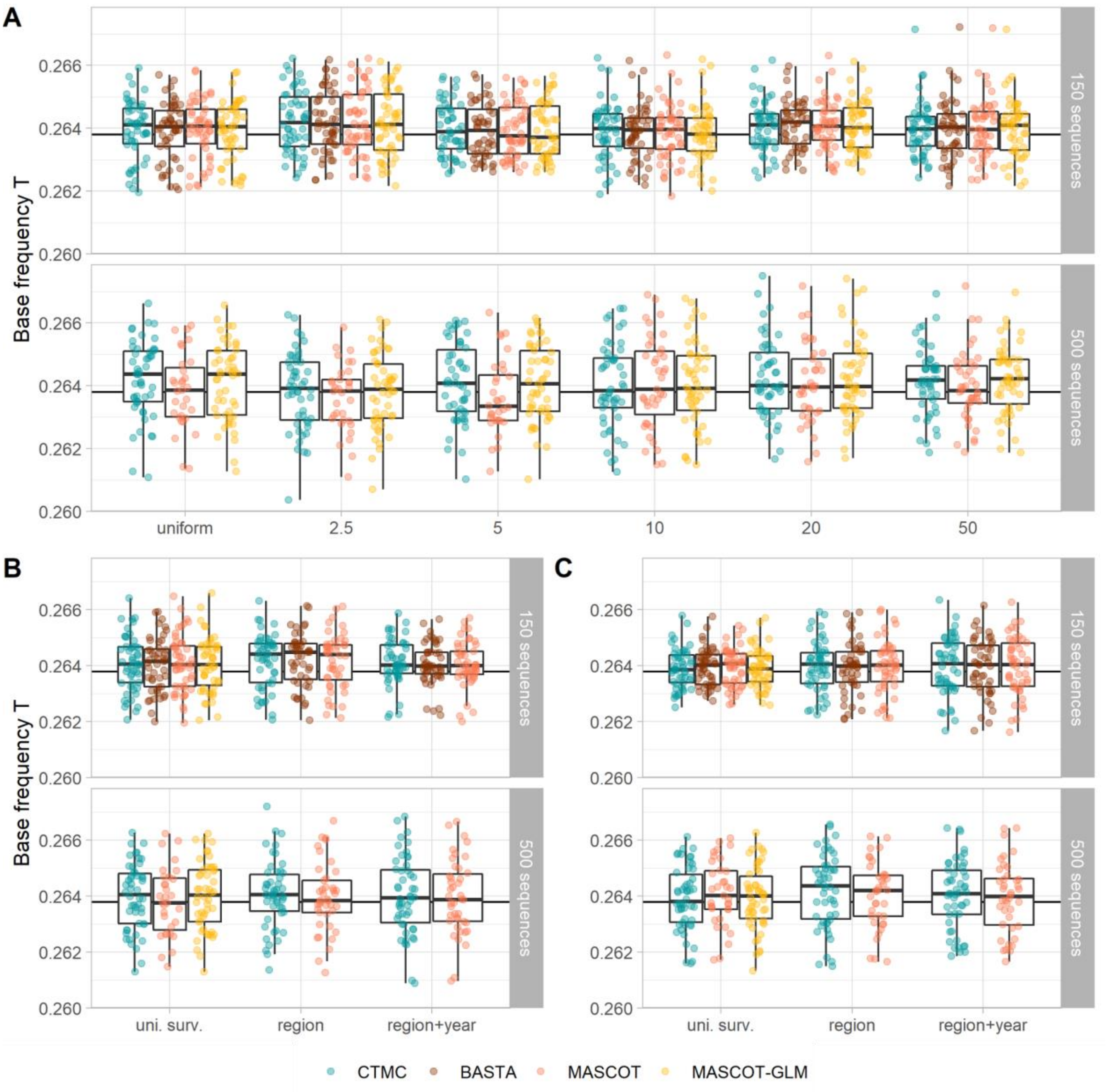


**Figure S5. Median estimates of base frequency T for all sampling conditions and the four algorithms.** Median estimate of the base frequency of T for the systematic bias (**A**), surveillance bias 10 (**B**), and surveillance bias 20 (**C**) sampling conditions. The true value of the parameter is represented as the horizontal black line.

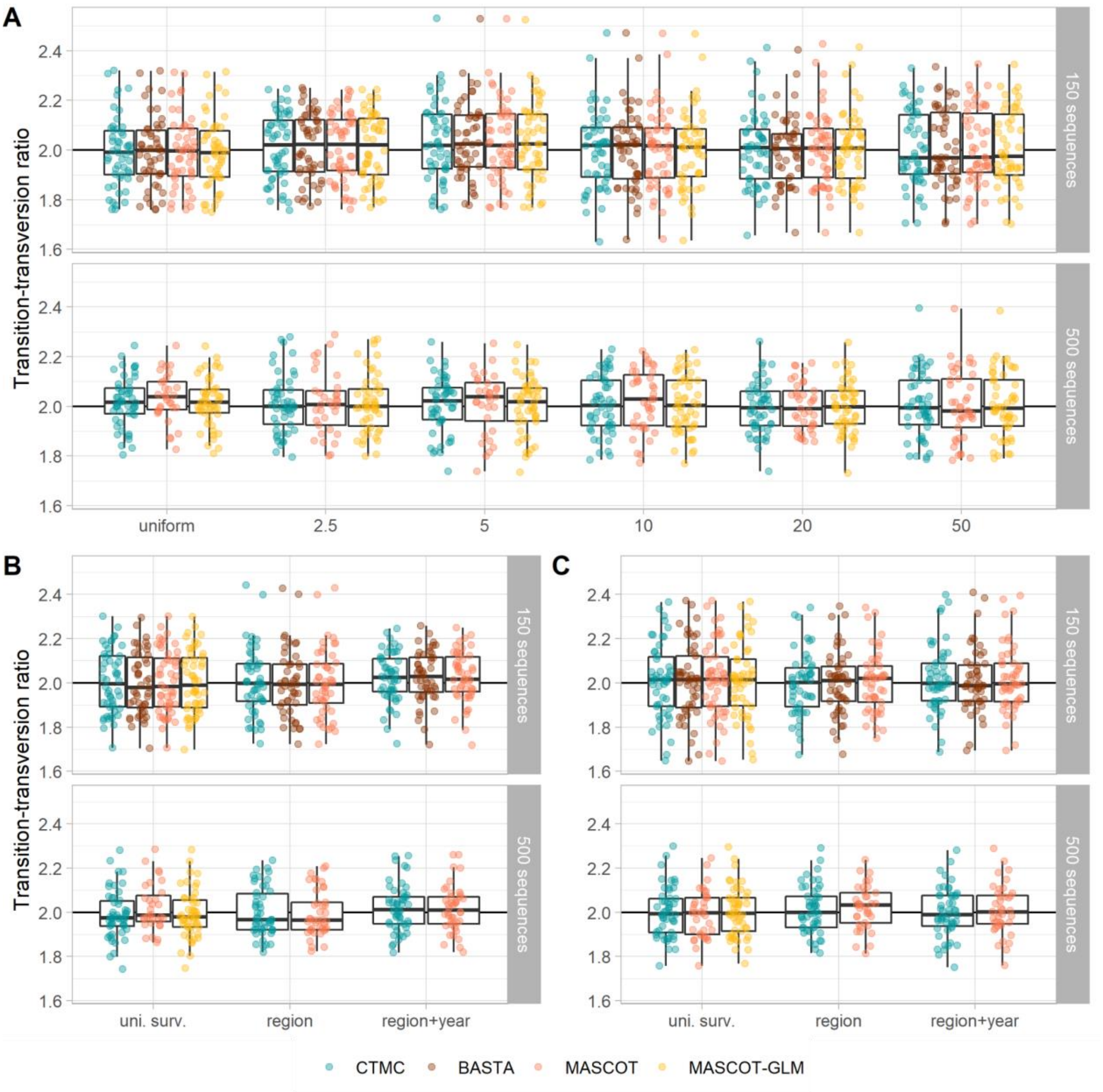


**Figure S6. Median estimates of the transition-transversion ratio for all sampling conditions and the four algorithms.** Median estimate of the transition-transversion ratio $\kappa$ for the systematic bias (**A**), surveillance bias 10 (**B**), and surveillance bias 20 (**C**) sampling conditions. The true value of the parameter is represented as the horizontal black line.

**b. Total migration counts**

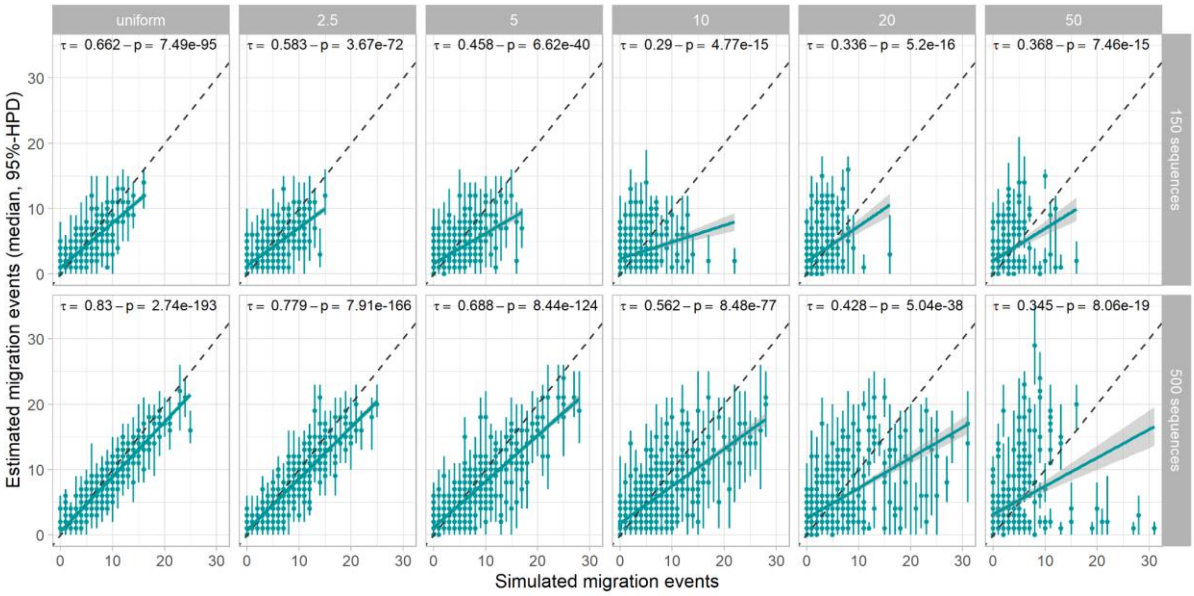


**Figure S7. Impact of bias on the estimation of the total migration counts by CTMC.** Median estimate of the base frequency of A for the systematic bias (**A**), surveillance bias 10 (**B**), and surveillance bias 20 (**C**) sampling conditions. The true value of the parameter is represented as the horizontal black line.

##### c. Lineage migration counts

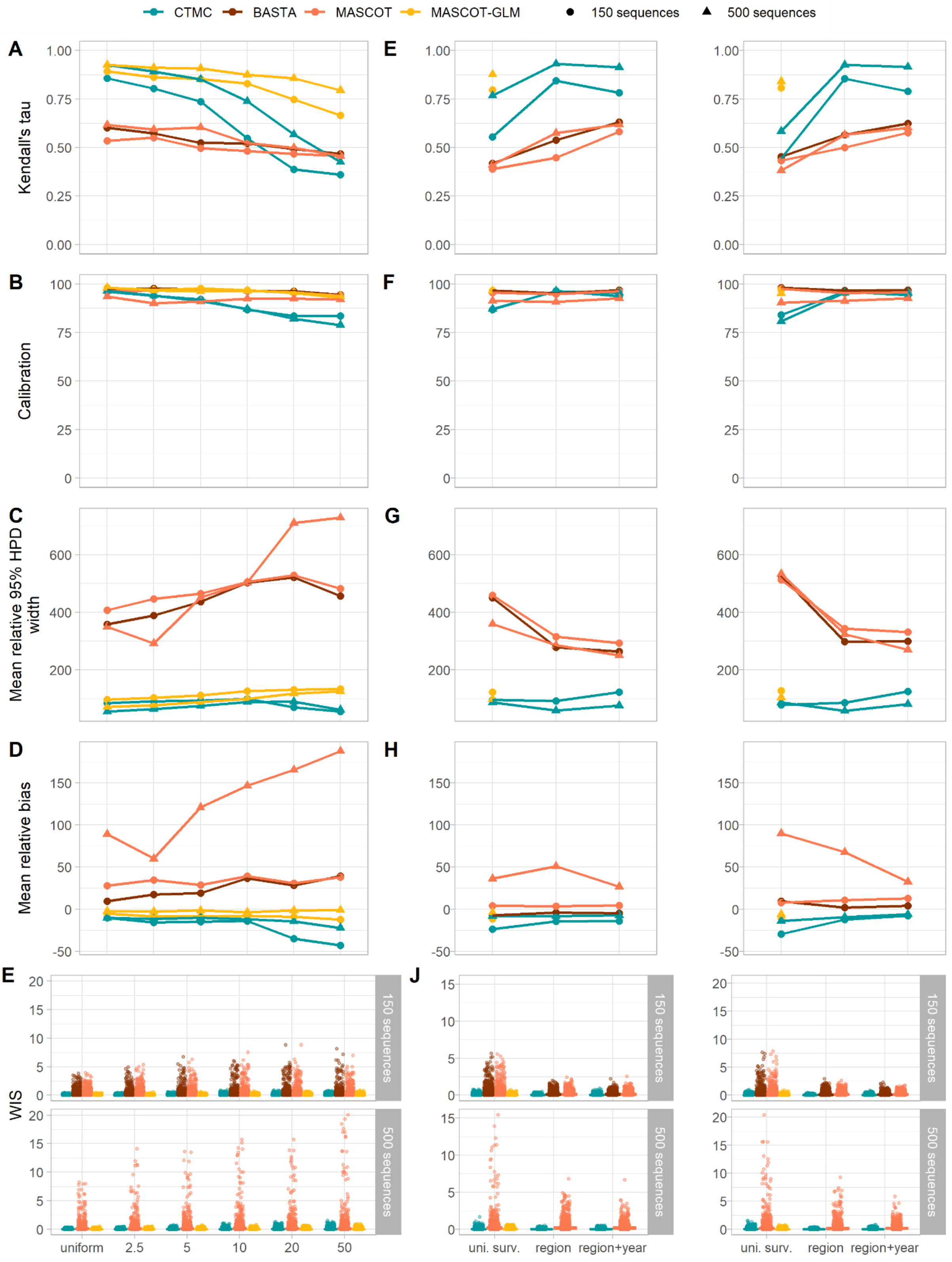


**Figure S8. Impact and mitigation of spatial bias on the estimation of the lineage migration counts. A-E**: Impact of the increasing levels of spatial bias on the correlation, the calibration, the mean relative 95% HPD width, the mean relative bias, and the WIS between the simulated and the estimated lineage migration counts. **F-J**: Mitigation of the impact of spatial bias on the correlation, the calibration, the mean relative 95% HPD width, the mean relative bias, and the WIS between the simulated and estimated lineage migration counts by using alternative sampling strategies. The mean relative bias and the mean relative 95% HPD width are not defined when the true value is null. We removed 65,399 out of 91,434 and 27,629 out of 41,916 simulated migration events in the small and large samples, respectively, due to true null values.

#### 2. Impact and mitigation of bias in the three demes framework

##### a. Estimation of genetic parameters

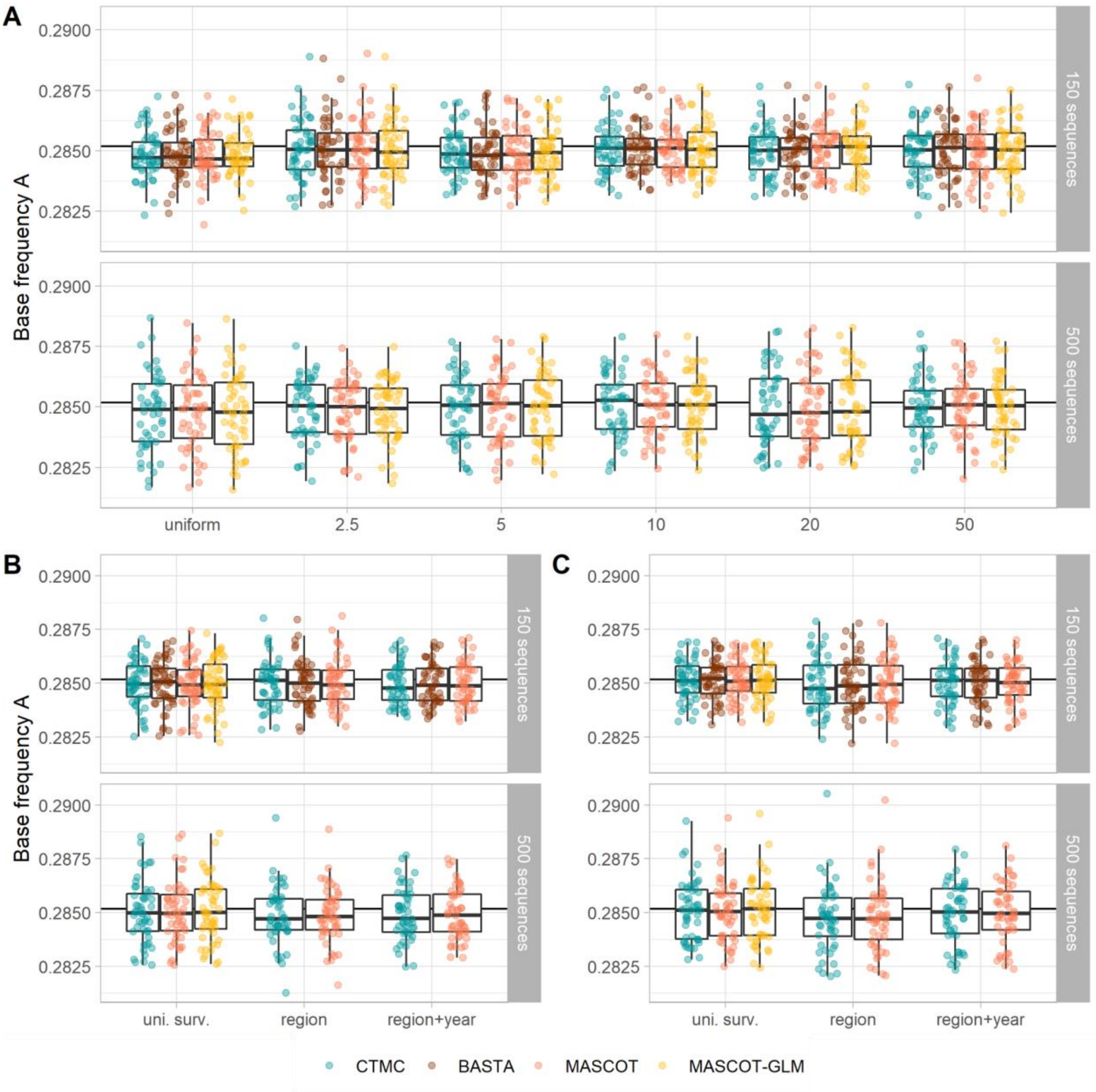


**Figure S9. Median estimates of base frequency A for all sampling conditions and the four algorithms.** Median estimate of the base frequency of A for the systematic bias (**A**), surveillance bias 10 (**B**), and surveillance bias 20 (**C**) sampling conditions. The true value of the parameter is represented as the horizontal black line.

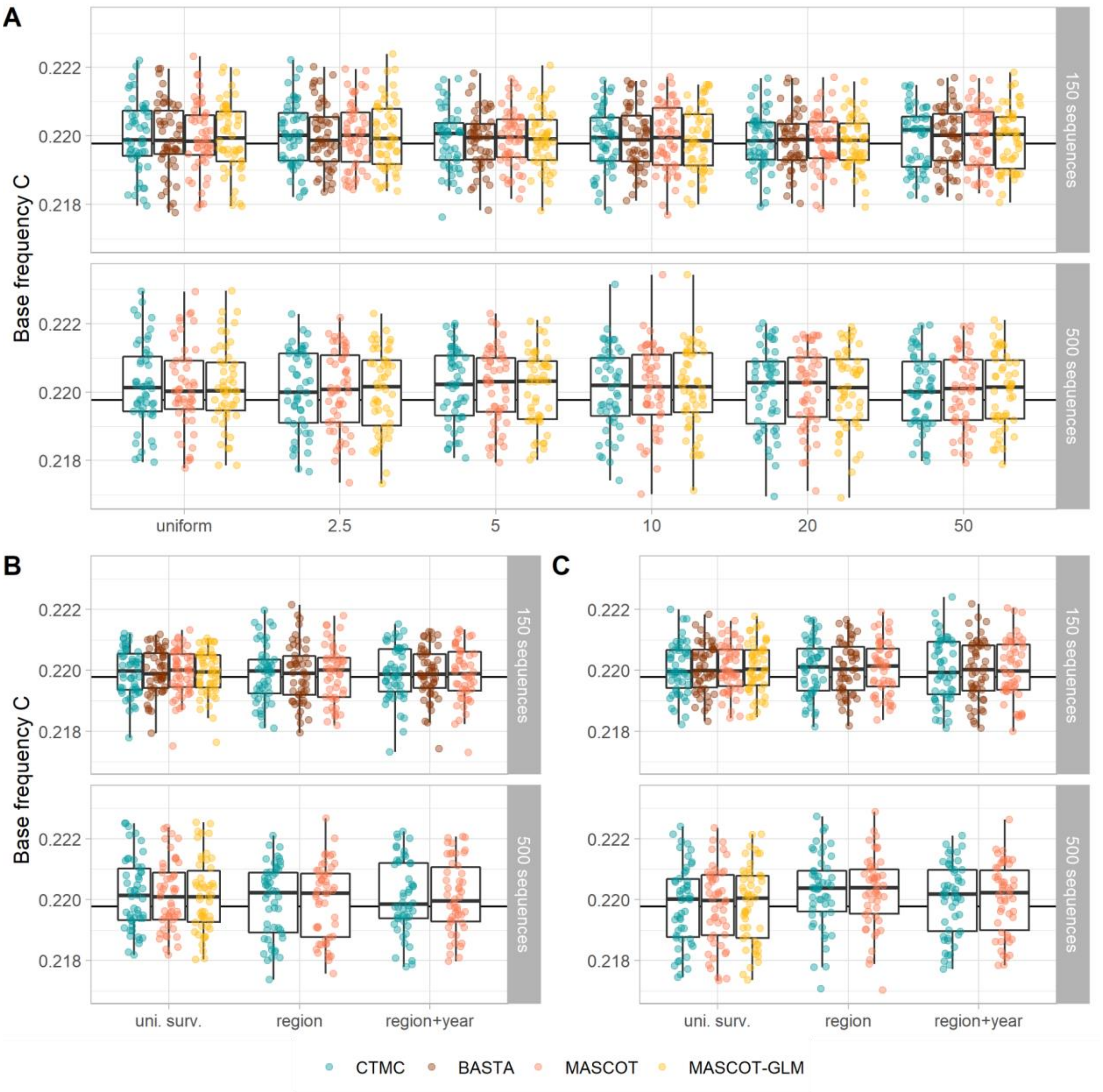


**Figure S10. Median estimates of base frequency C for all sampling conditions and the four algorithms.** Median estimate of the base frequency of C for the systematic bias (**A**), surveillance bias 10 (**B**), and surveillance bias 20 (**C**) sampling conditions. The true value of the parameter is represented as the horizontal black line.

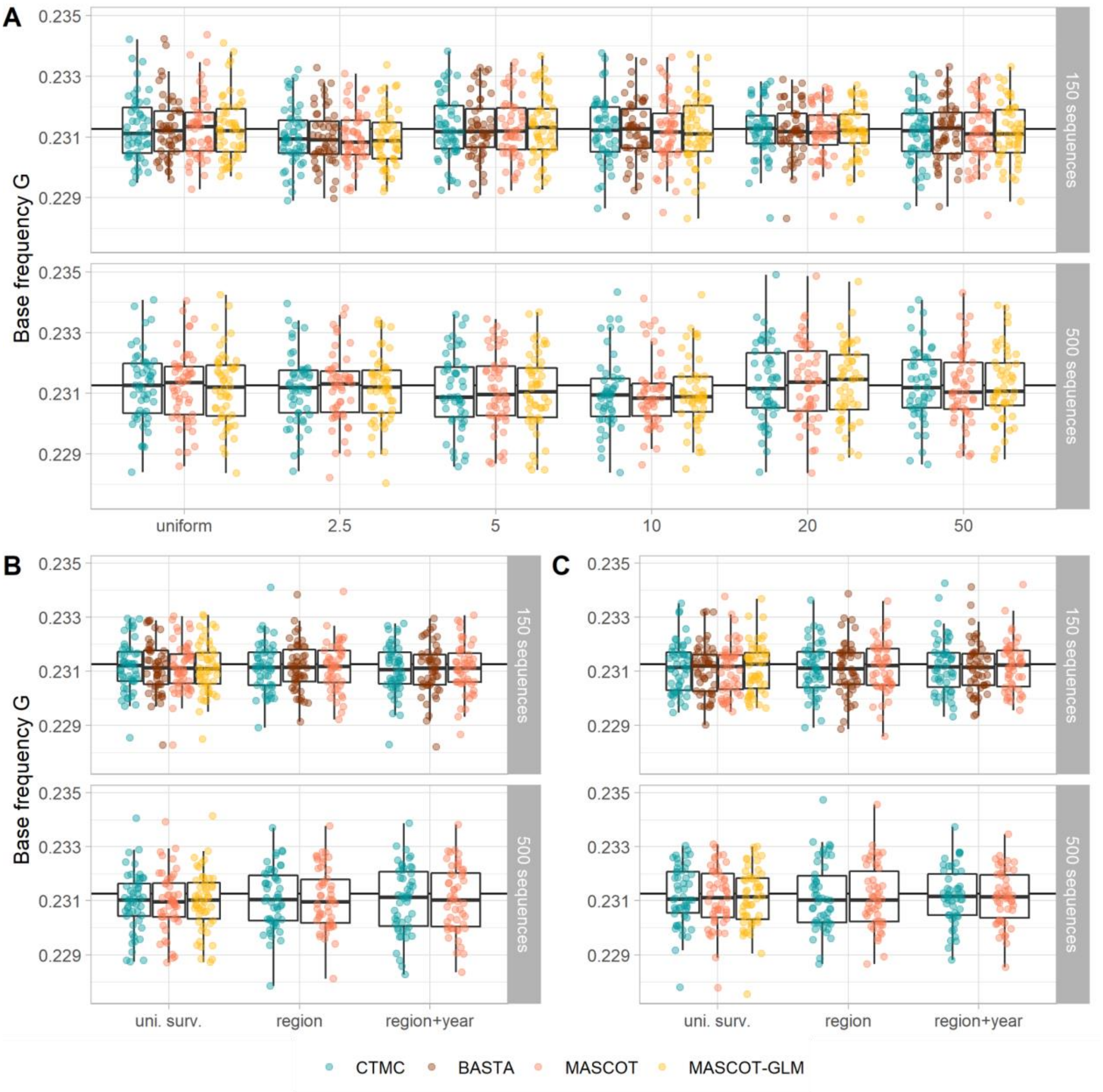


**Figure S11. Median estimates of base frequency G for all sampling conditions and the four algorithms.** Median estimate of the base frequency of G for the systematic bias (**A**), surveillance bias 10 (**B**), and surveillance bias 20 (**C**) sampling conditions. The true value of the parameter is represented as the horizontal black line.

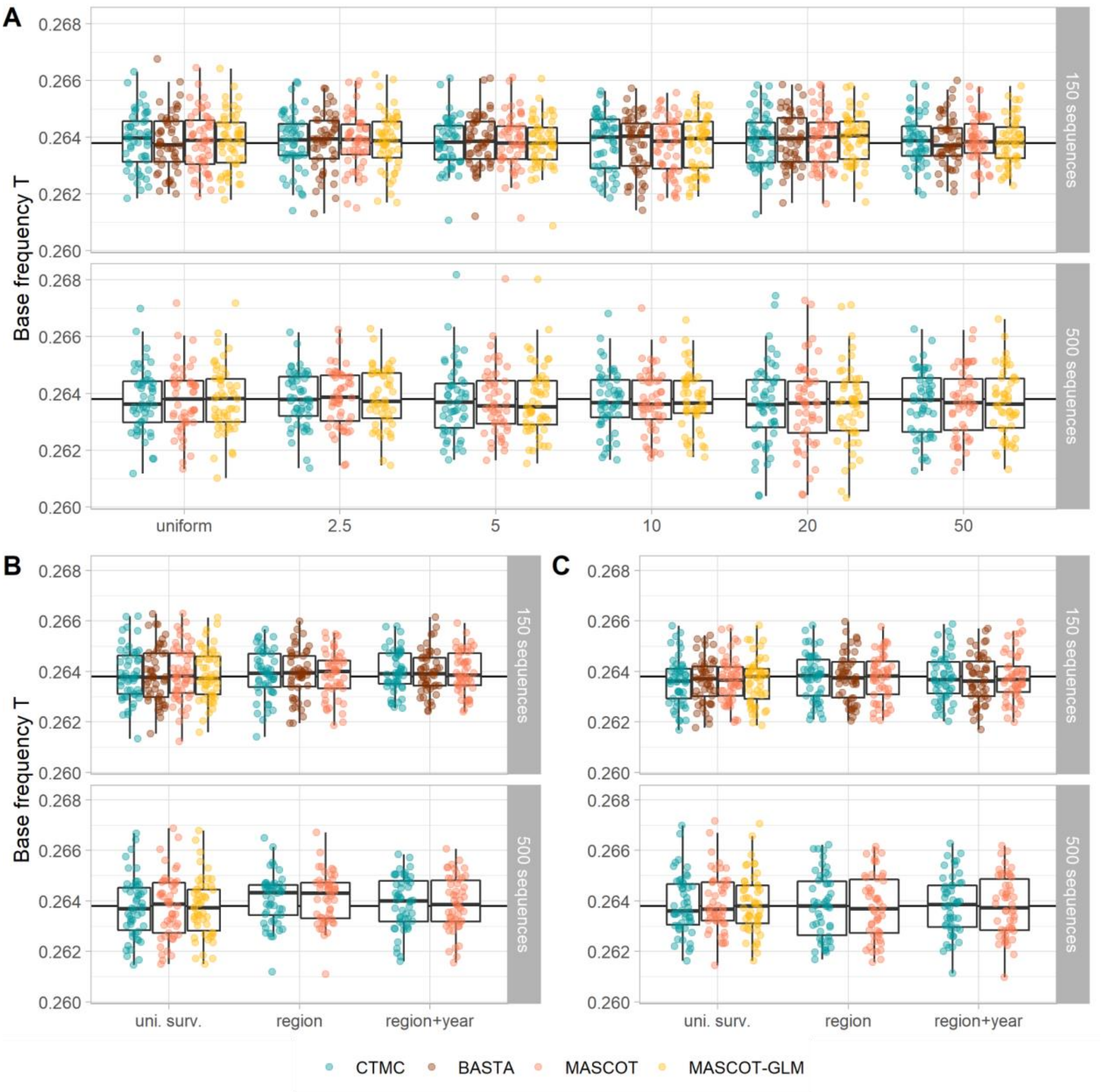


**Figure S12. Median estimates of base frequency T for all sampling conditions and the four algorithms.** Median estimate of the base frequency of T for the systematic bias (**A**), surveillance bias 10 (**B**), and surveillance bias 20 (**C**) sampling conditions. The true value of the parameter is represented as the horizontal black line.

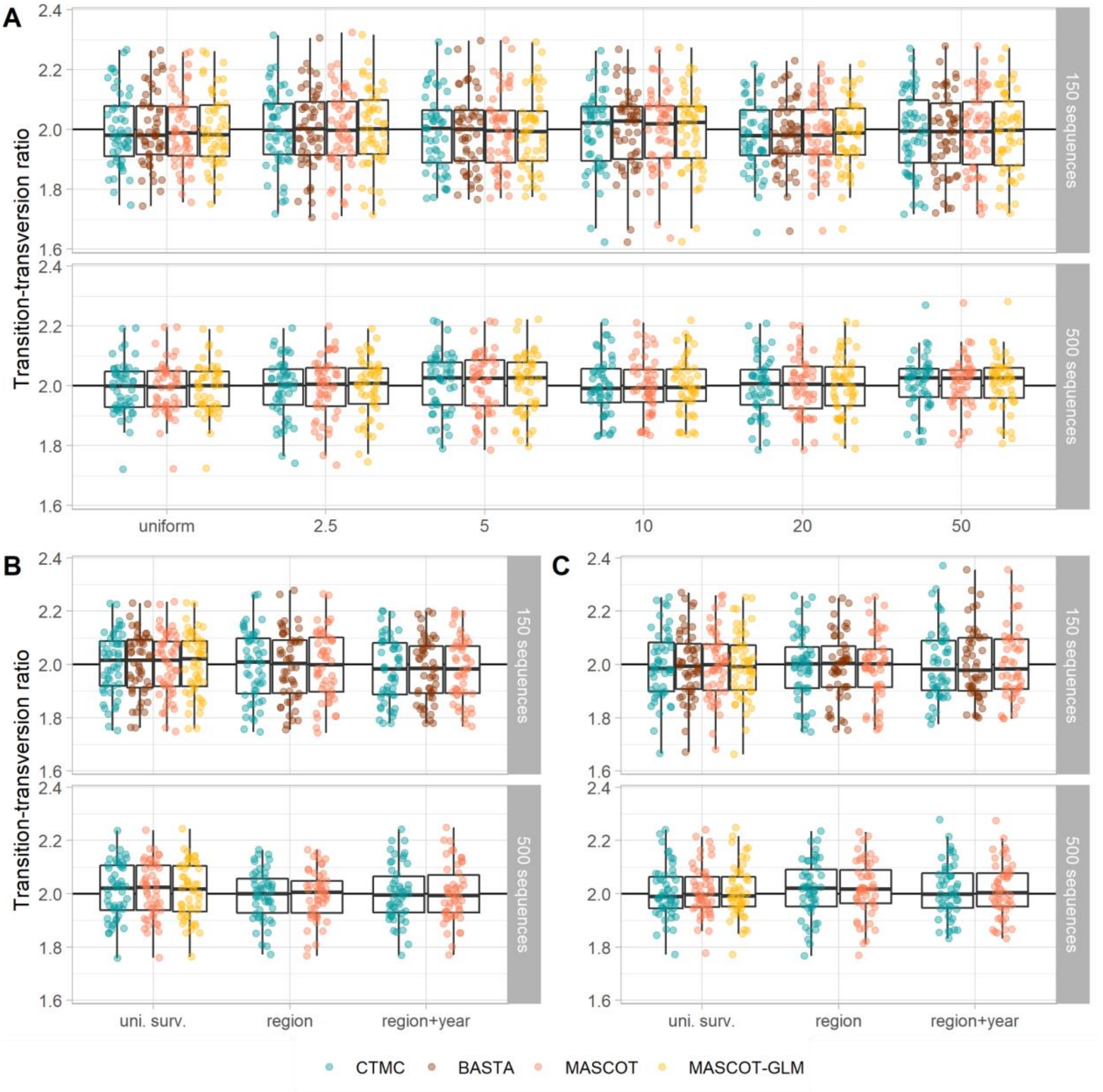


**Figure S13. Median estimates of the transition-transversion ratio for all sampling conditions and the four algorithms.** Median estimate of the transition-transversion ratio $\kappa$ for the systematic bias (**A**), surveillance bias 10 (**B**), and surveillance bias 20 (**C**) sampling conditions. The true value of the parameter is represented as the horizontal black line.

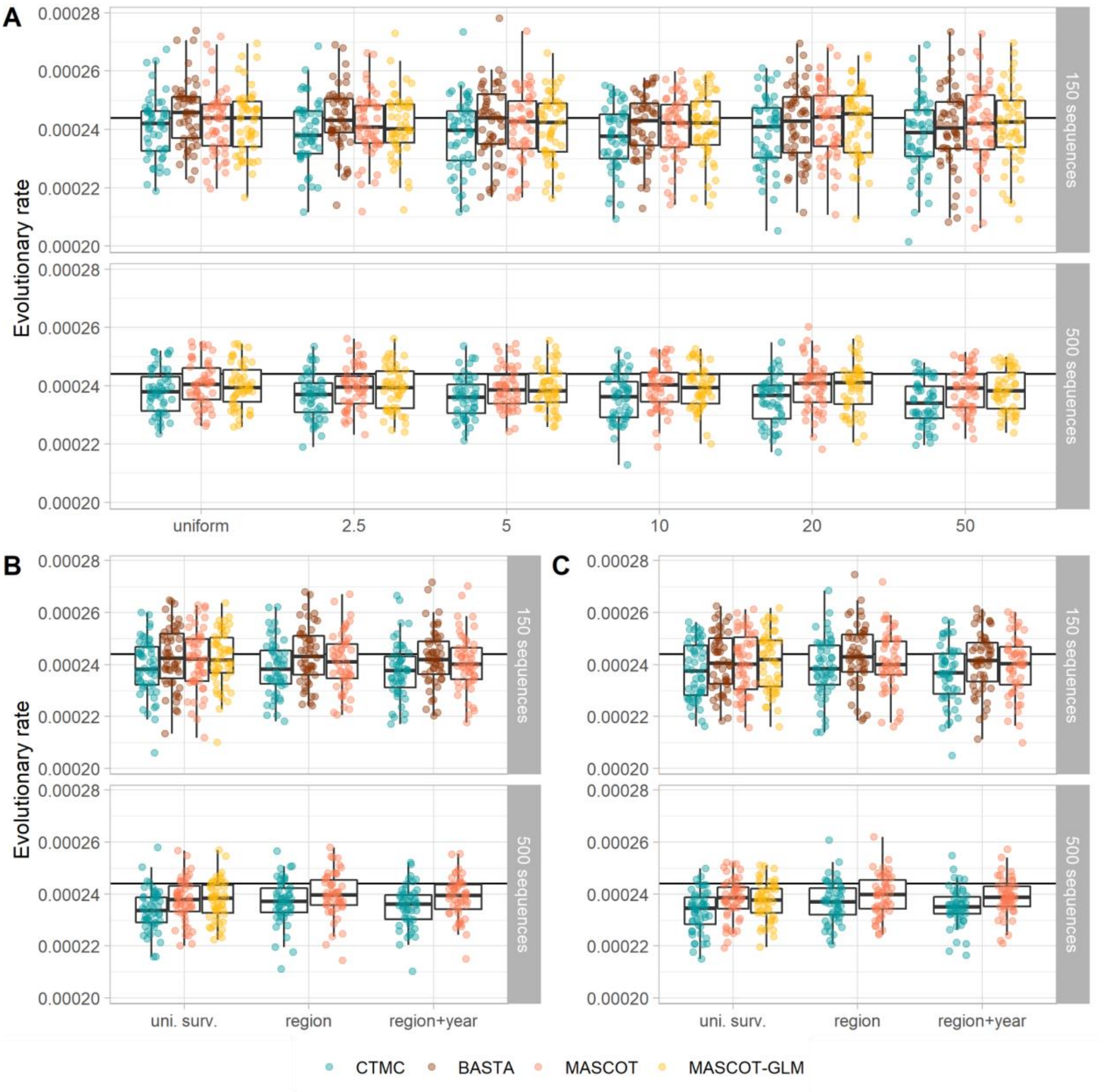


**Figure S14. Median estimates of the evolutionary rate for all sampling conditions and the four algorithms.** Median estimate of the evolutionary rate for the systematic bias (**A**), surveillance bias 10 (**B**), and surveillance bias 20 (**C**) sampling conditions. The true value of the parameter is represented as the horizontal black line.

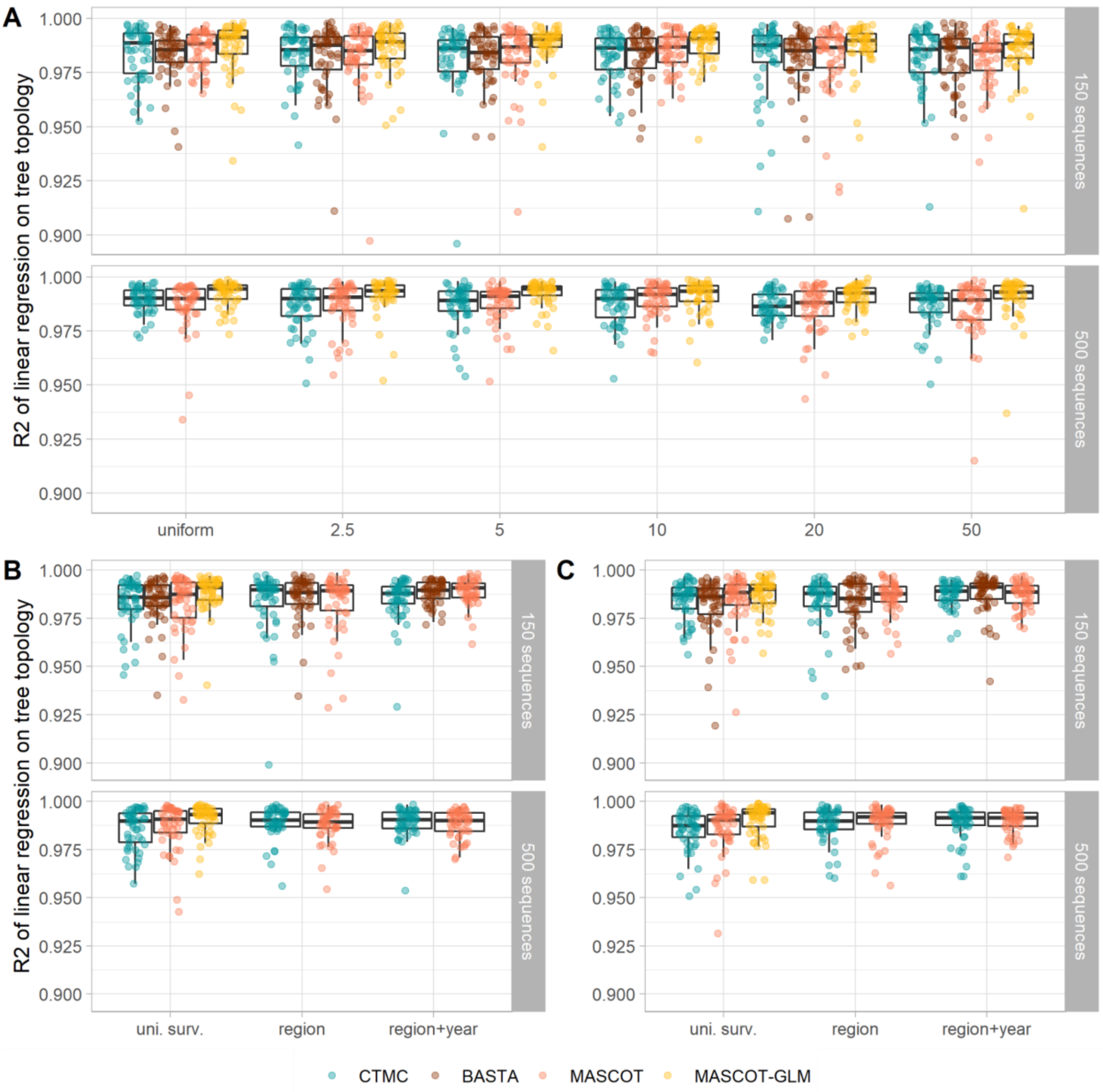


**Figure S15. Comparison of the simulated and estimated tree topologies for all sampling conditions and the four algorithms.** Pearson's determination coefficient of the pairwise divergence time between the simulated transmission chain and the MCC tree for the systematic bias (**A**), surveillance bias 10 (**B**), and surveillance bias 20 (**C**) sampling conditions.

**b. Lineage migration counts**

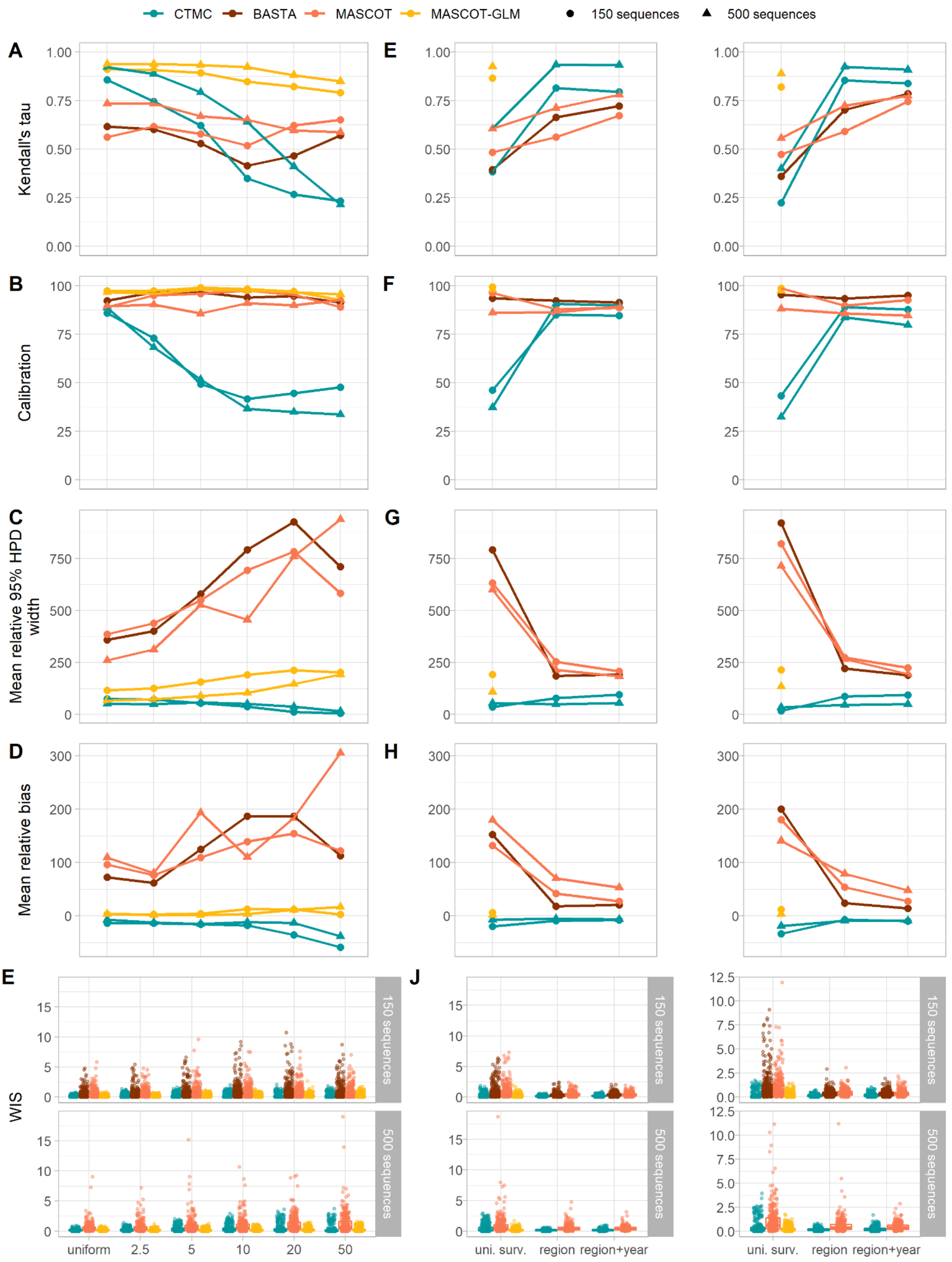


**Figure S16. Impact and mitigation of spatial bias on the estimation of the lineage migration counts. A-E**: Impact of the increasing levels of spatial bias on the correlation, the calibration, the mean relative 95% HPD width, the mean relative bias, and the WIS between the simulated and the estimated lineage migration counts. **F-J**: Mitigation of the impact of spatial bias on the correlation, the calibration, the mean relative 95% HPD width, the mean relative bias, and the WIS between the simulated and estimated lineage migration counts by using alternative sampling strategies. The mean relative bias and the mean relative 95% HPD width are not defined when the true value is null. We removed 3,126 out of 13,200 and 1,410 out of 9,588 simulated migration events in the small and large samples, respectively, due to null true values.

**c. Total migration counts**

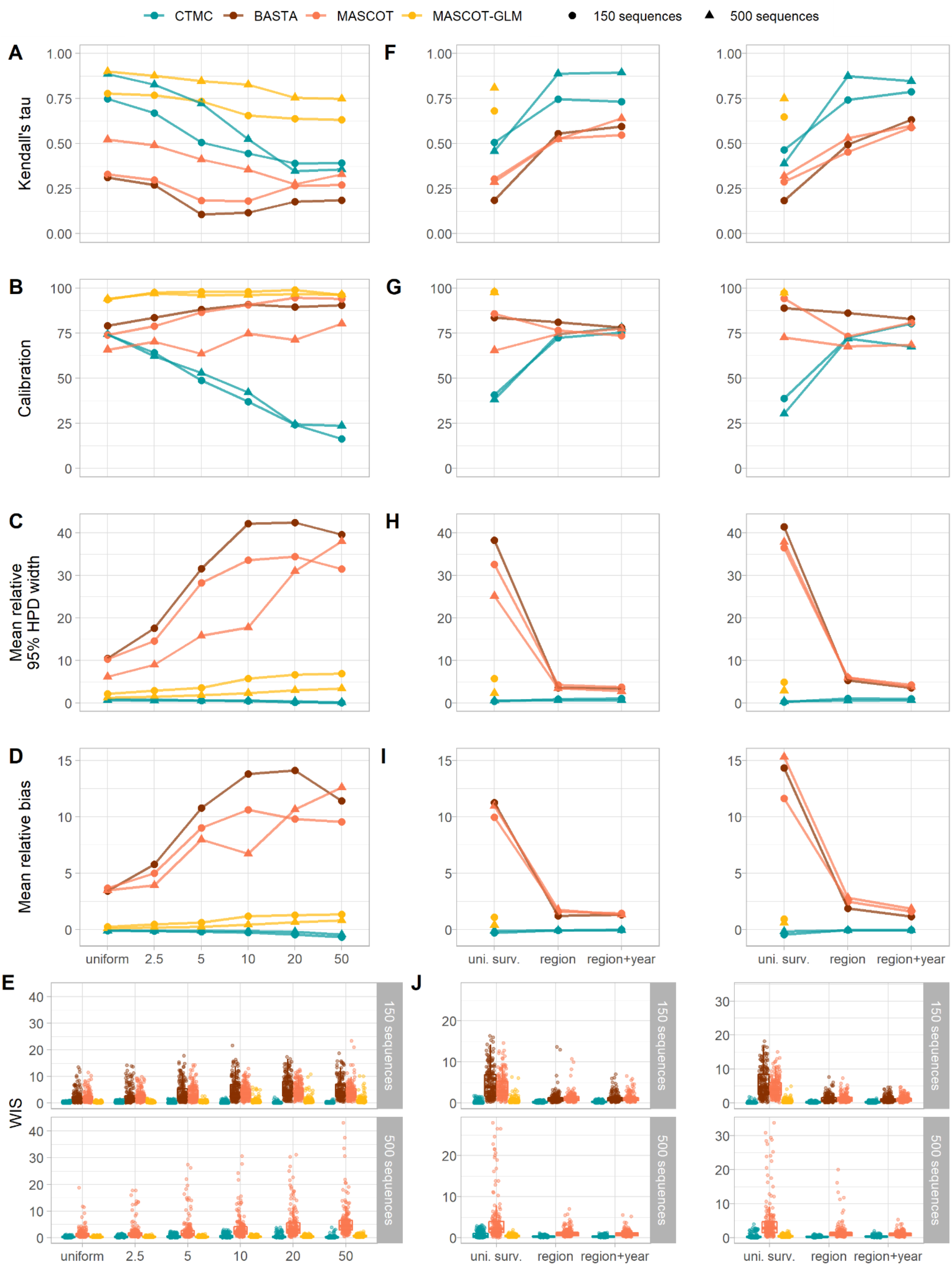

**Figure S17. Impact and mitigation of spatial bias on the estimation of the total migration counts. A-E**: Impact of the increasing levels of spatial bias on the correlation, the calibration, the mean relative 95% HPD width, the mean relative bias, and the WIS between the simulated and the estimated total migration counts. **F-J**: Mitigation of the impact of spatial bias on the correlation, the calibration, the mean relative 95% HPD width, the mean relative bias, and the WIS between the simulated and estimated total migration counts by using alternative sampling strategies. The mean relative bias and the mean relative 95% HPD width are not defined when the true value is null. We removed 612 out of 3,600 and 380 out of 3,600 simulated migration events in the small and large samples, respectively, due to true null values.

**d. Introduction dates**

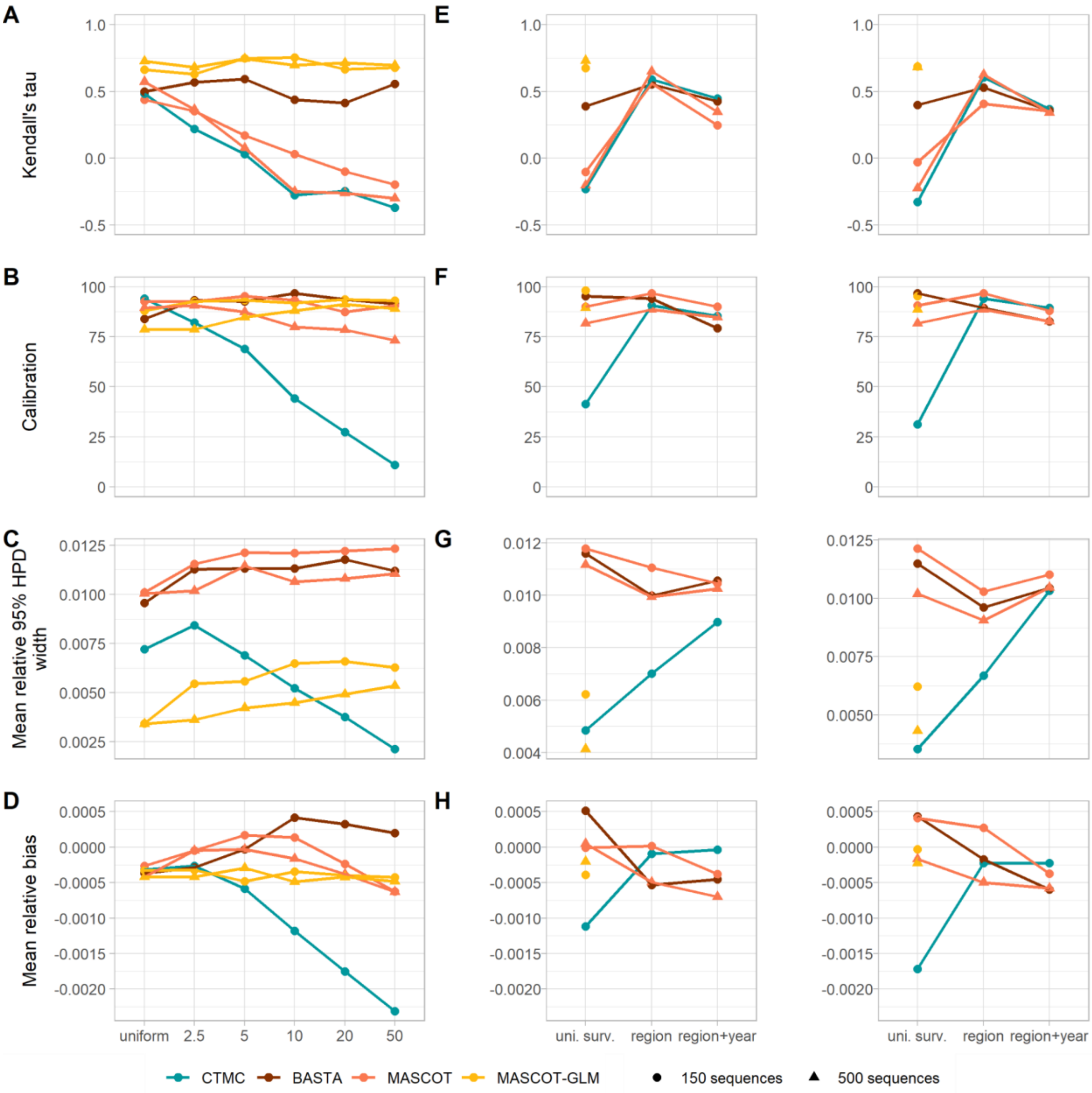


**Figure S18. Impact and mitigation of spatial bias on the estimation of the introduction dates. A-D**: Impact of the increasing levels of spatial bias on correlation, calibration, mean relative 95% HPD width, and average relative error between the simulated and estimated introduction dates. **E-H**: Mitigation of the impact of spatial bias on correlation, calibration, mean relative 95% HPD width, and average relative error between the simulated and estimated introduction dates by using alternative sampling strategies.

e. **Root location**

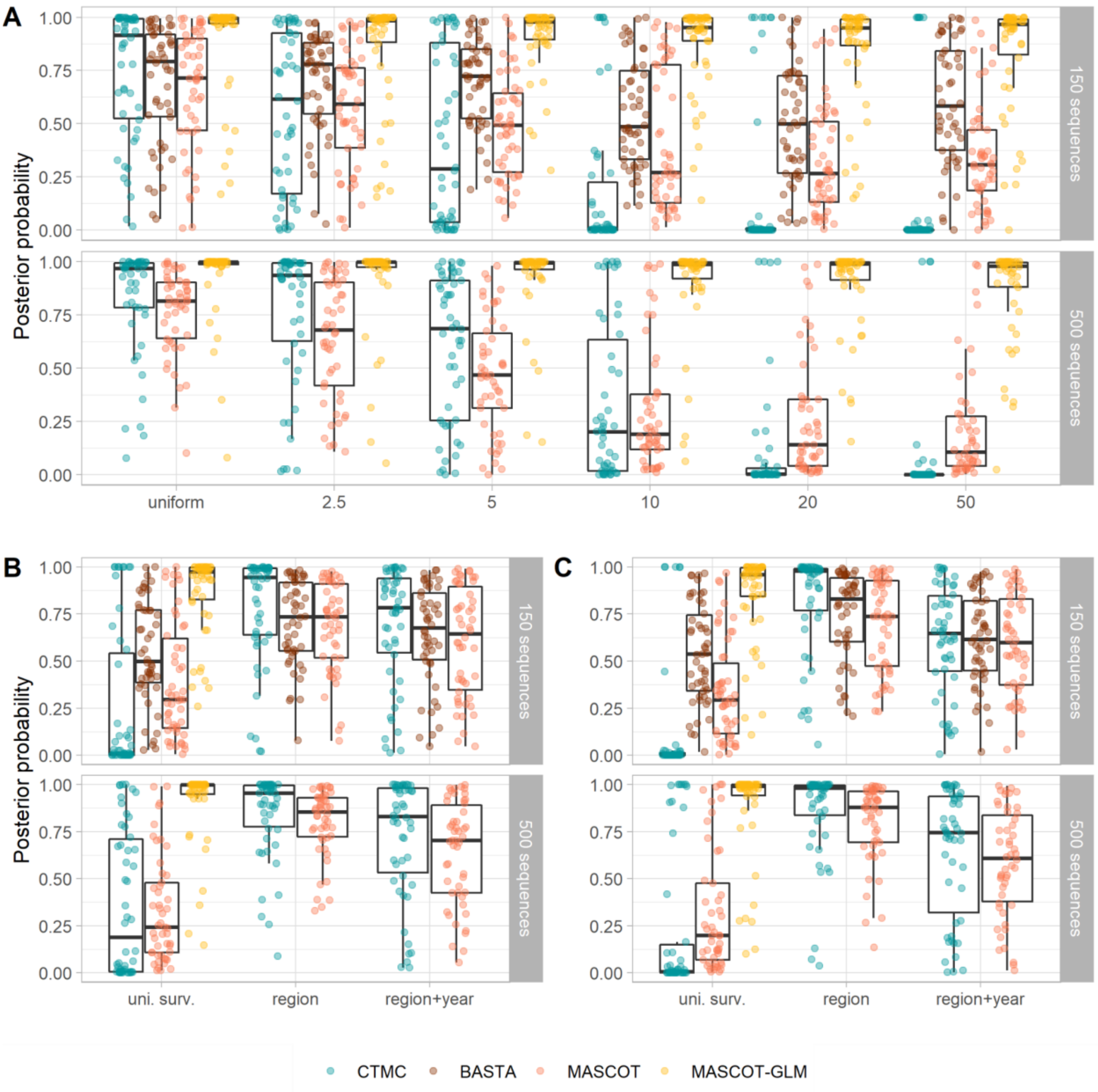


**Figure S19. Impact and mitigation of spatial bias on the estimation of the root location. A**: Decreasing root state posterior probability with an increasing biais. **B-C**: Mitigation of the effects of spatial biais using alternative sampling strategies with an underlying bias of 10 and 20, respectively. Each dot corresponds to the median root state posterior probability in one simulation (n = 50 per sampling protocol and sample size).

#### 3. RABV spread in the Philippines

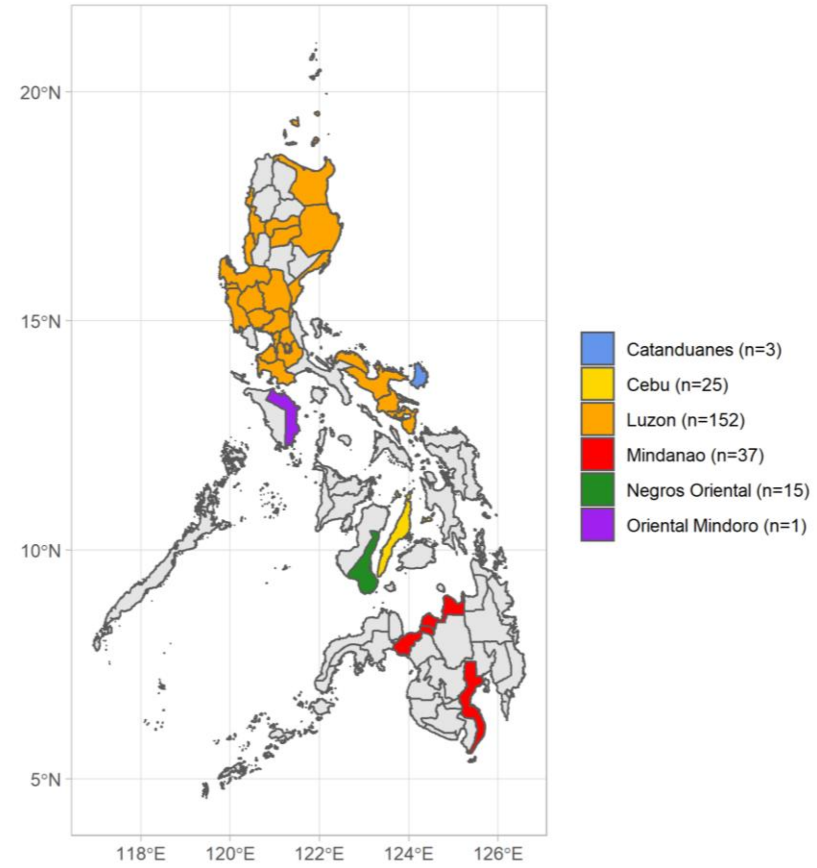


**Figure S20. Sampling location and intensity of the RABV sequences in the Philippines, 2004-2010.**

#### 4. SARS-CoV-2 early spread

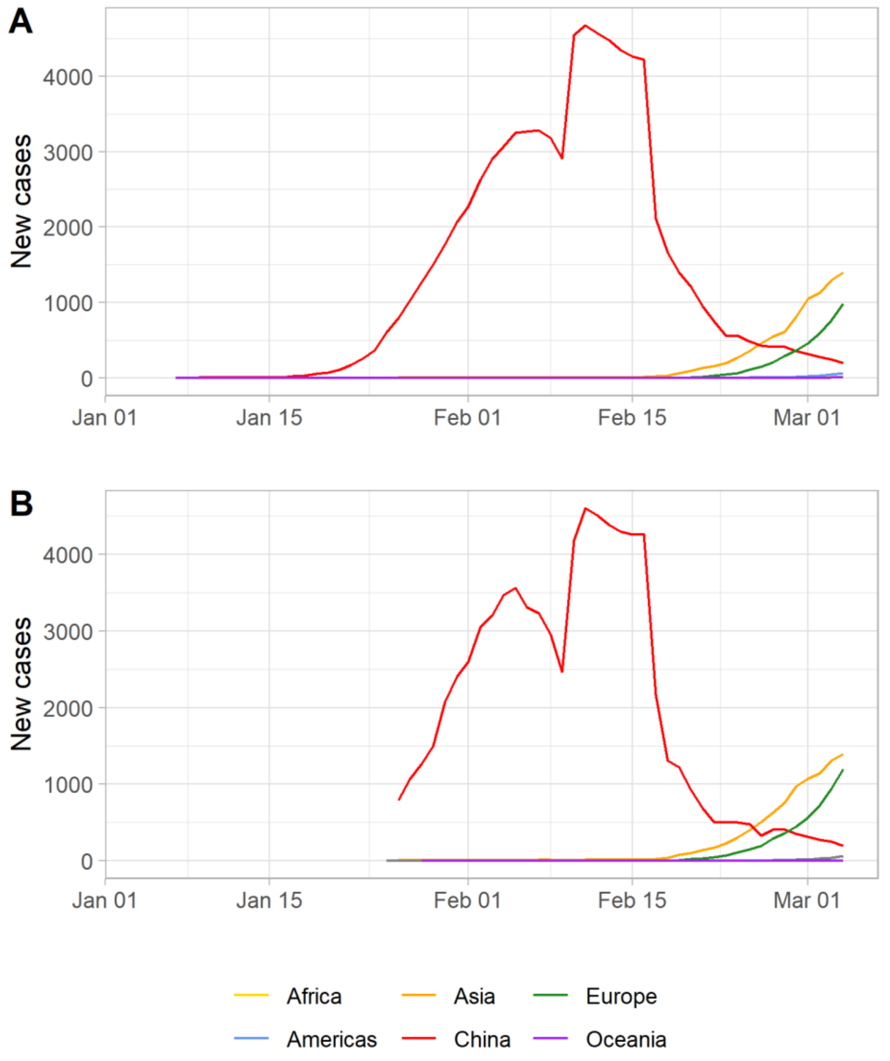


**Figure S21. Case count data by continent used to inform MASCOT. A-B**: Case count data from the World Health Organization and Our World In Data, respectively. We report the moving average over a seven-day window.

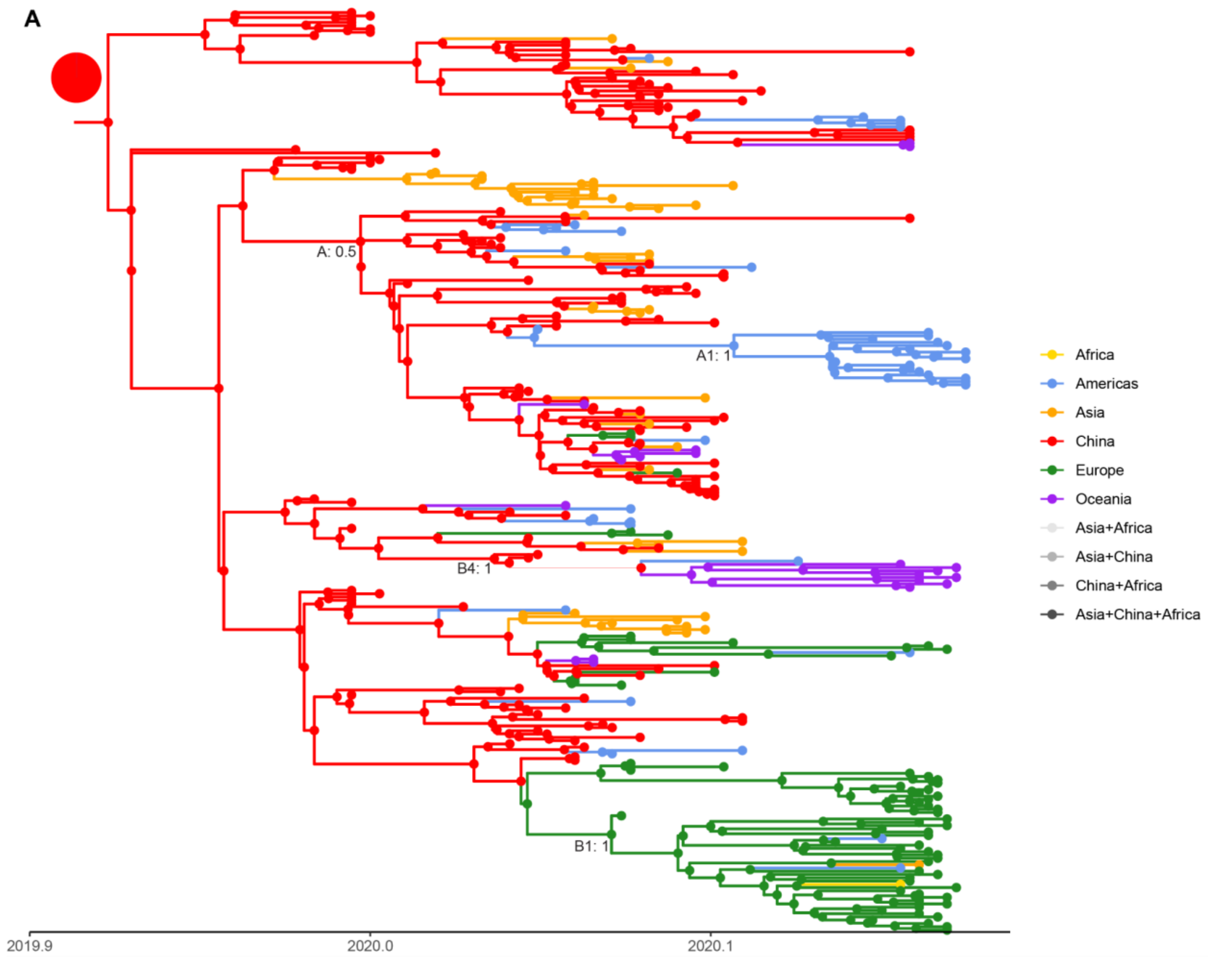
A
A: 0.5
A1: 1
B4: 1
B1: 1
2019.9
2020.0
2020.1
Africa
Americas
Asia
China
Europe
Oceania
Asia+Africa
Asia+China
China+Africa
Asia+China+Africa

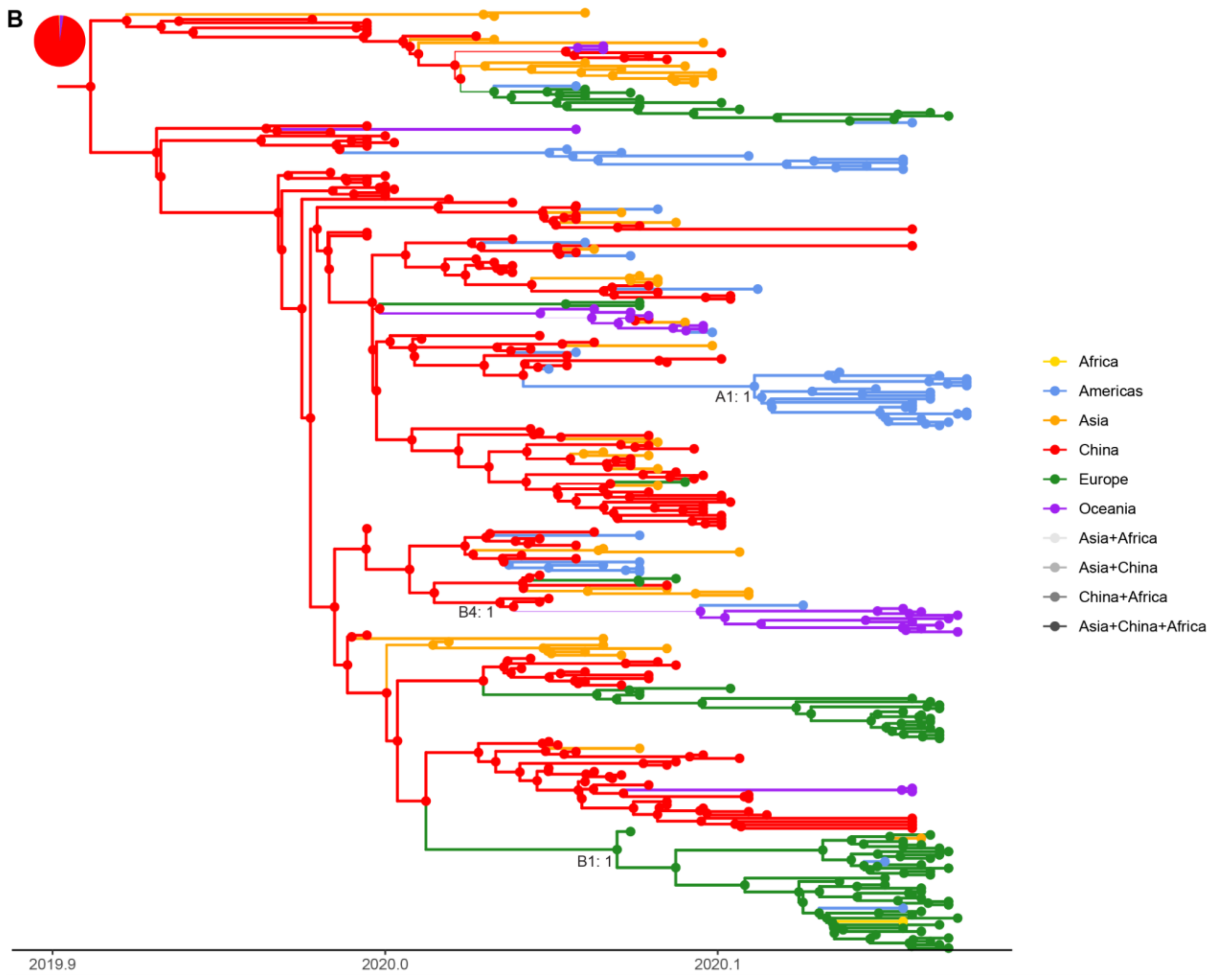
B
A1: 1
B4: 1
B1: 1
2019.9
2020.0
2020.1
Africa
Americas
Asia
China
Europe
Oceania
Asia+Africa
Asia+China
China+Africa
Asia+China+Africa

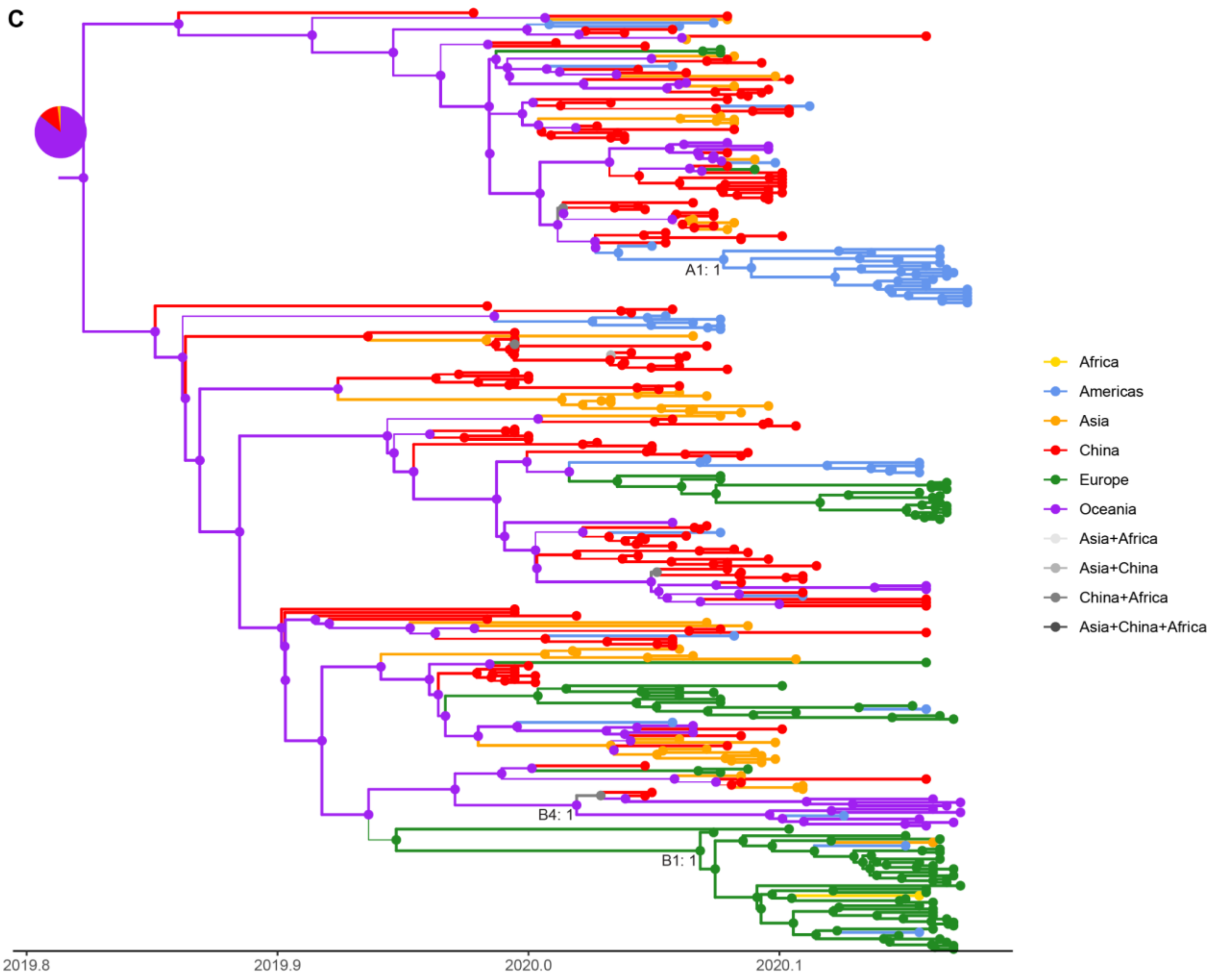
C
A1: 1
B4: 1
B1: 1
Africa
Americas
Asia
China
Europe
Oceania
Asia+Africa
Asia+China
China+Africa
Asia+China+Africa
2019.8
2019.9
2020.0
2020.1

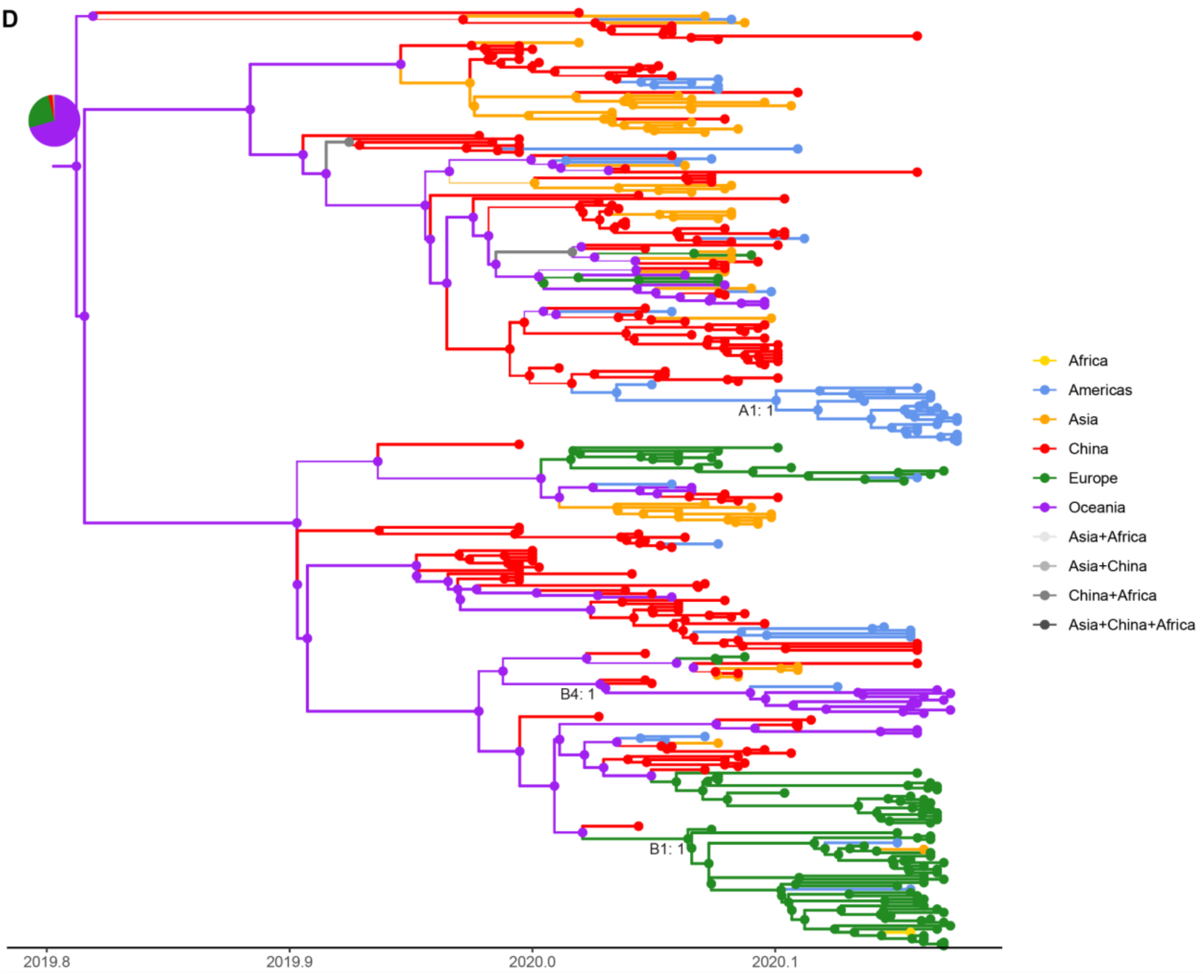

D
A1: 1
B4: 1
B1: 1
2019.8
2019.9
2020.0
2020.1
Africa
Americas
Asia
China
Europe
Oceania
Asia+Africa
Asia+China
China+Africa
Asia+China+Africa

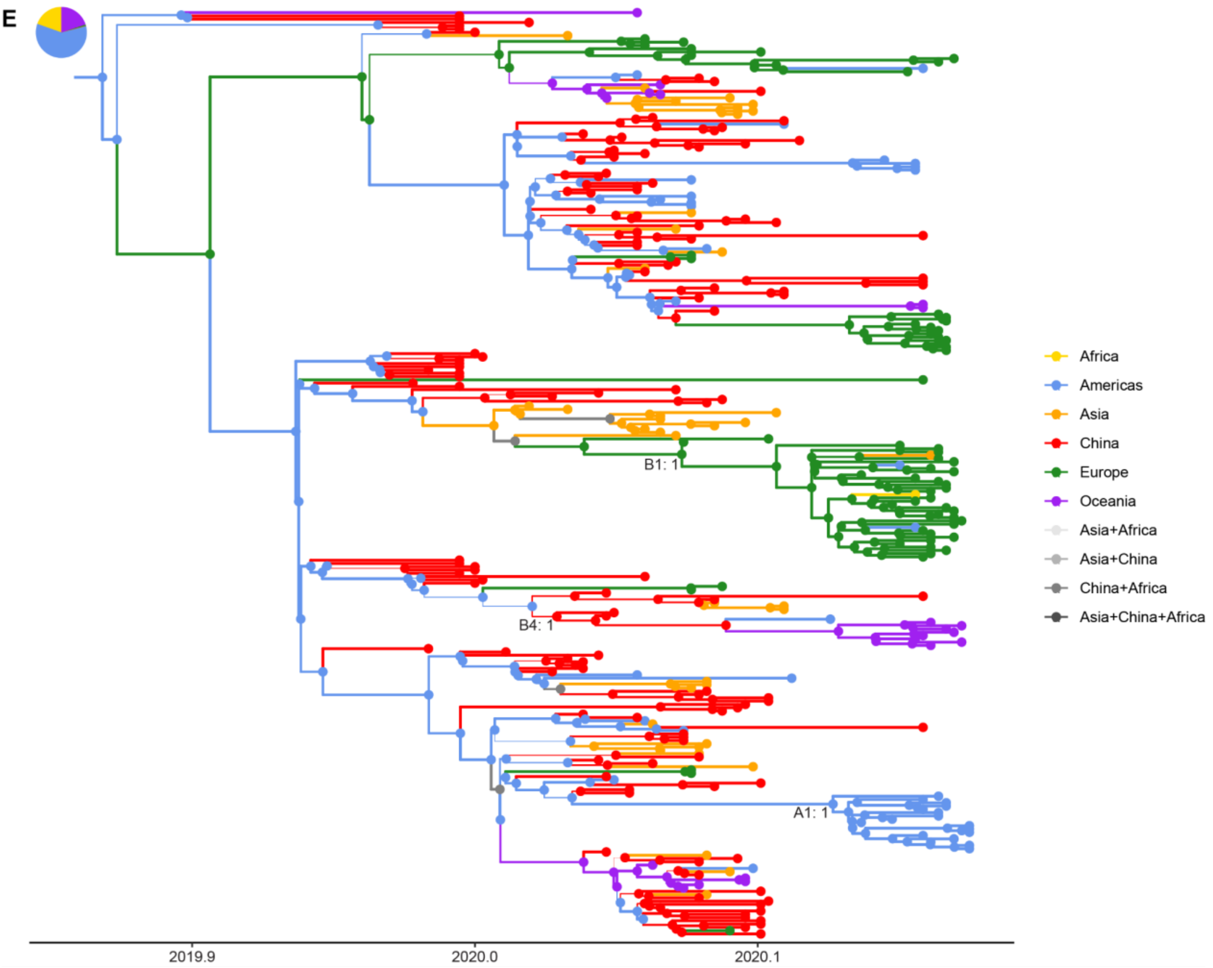
E
B1: 1
B4: 1
A1: 1
2019.9
2020.0
2020.1
Africa
Americas
Asia
China
Europe
Oceania
Asia+Africa
Asia+China
China+Africa
Asia+China+Africa

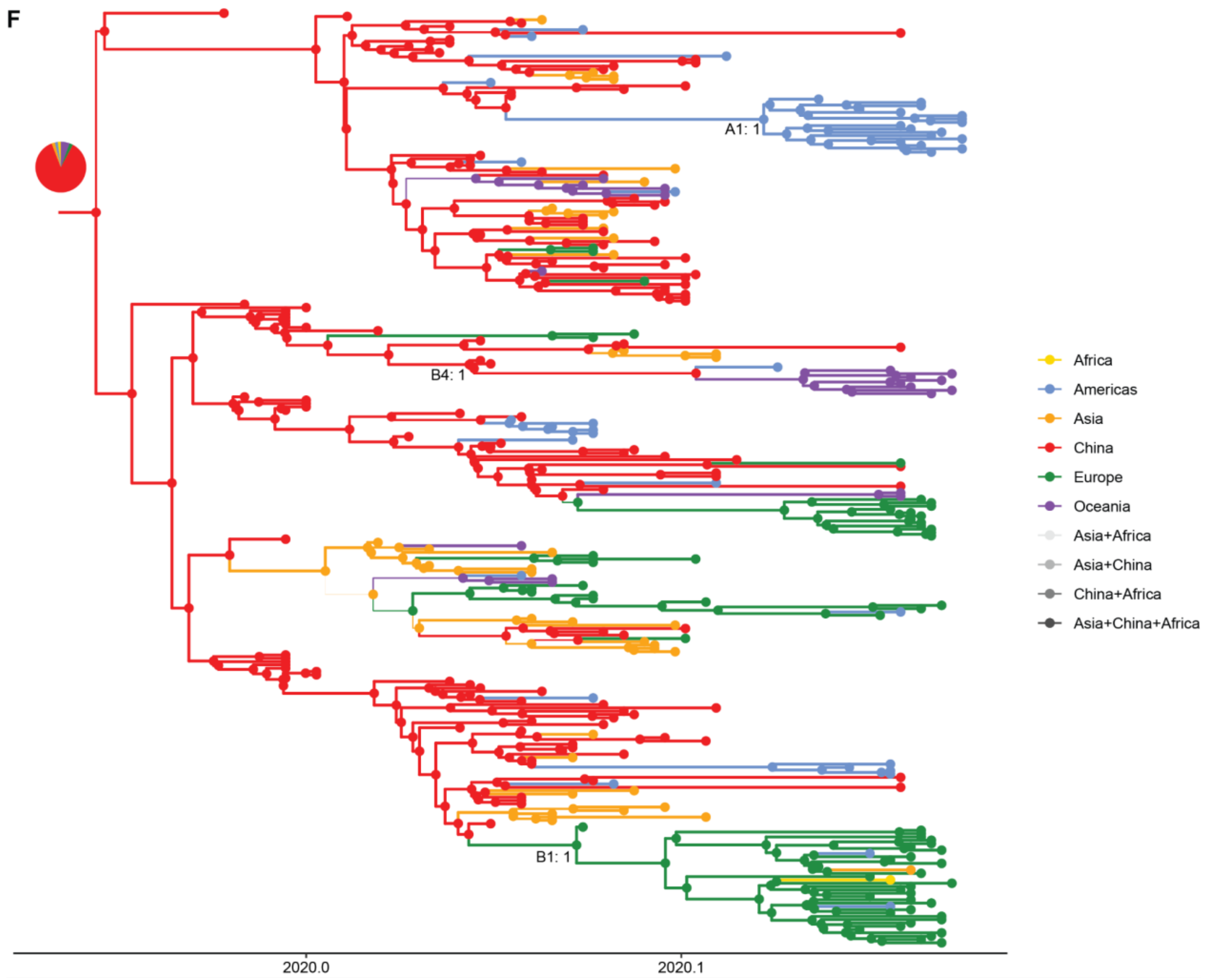


**Figure S22. Maximum clade credibility (MCC) tree of SARS-CoV-2 genomes from the early stages of the pandemic for all the algorithms. A-F**: MCC trees for CTMC, BASTA (1st mode of evolutionary rate), BASTA (2nd mode of evolutionary rate), MASCOT, MASCOT-WID, and MASCOT-WHO respectively. The posterior support of lineages A, A1, B1, and B4 that were identified by Lemey et al. (1) are reported at the corresponding nodes when they gather the same sequences as in the original analysis. Lineages A1, B1, and B4 were estimated to be monophyletic with a high posterior support by all algorithms. However, BASTA, MASCOT, MASCOT-WID, and MASCOT-WHO did not infer monophyly for lineage A that is why we did not report its posterior supports on the corresponding MCC trees. Branch width is proportional to the maximal location probability of the parent node. Nodes and branches are colored by location with maximal probability. The root location probability distribution is reported in the pie chart.

**Table S1. Predicted lineage location on the SARS-CoV-2 data.** We report the predicted locations with maximal probability of the four SARS-CoV-2 lineages estimated by all algorithms along with their posterior probability. We did not report the location of lineage A predicted by BASTA, MASCOT, MASCOT-WID, and MASCOT-WHO because these algorithms did not infer a monophyletic lineage. Location of lineage A1 and B1 is estimated in a similar fashion by all algorithms. However, lineage B4 was estimated to be located in Oceania by MASCOT and the 2nd mode of BASTA, whereas it is estimated to be located in China by CTMC-TRAVEL and CTMC. When we add case count data from Our World In Data and the WHO, the estimated lineage location is China but the posterior probability for MASCOT-WID is lower than for CTMC-TRAVEL and CTMC.

| | **Lineage A1** | **Lineage A** | **Lineage B1** | **Lineage B4** |
|---|---|---|---|---|
| **CTMC-TRAVEL** | Americas (1) | China (1) | Europe (0.761) | China (1) |
| **CTMC** | Americas (0.995) | China (1) | Europe (0.998) | China (1) |
| **BASTA - 1st mode** | Americas (0.958) | - | Europe (0.985) | China (0.983) |
| **BASTA - 2nd mode** | Americas (0.97) | - | Europe (0.977) | Oceania (0.851) |
| **MASCOT** | Americas (0.994) | - | Europe (0.993) | Oceania (0.781) |
| **MASCOT-WID** | Americas (0.937) | - | Europe (0.993) | China (0.446) |
| **MASCOT-WHO** | Americas (0.959) | - | Europe (0.977) | China (0.975) |

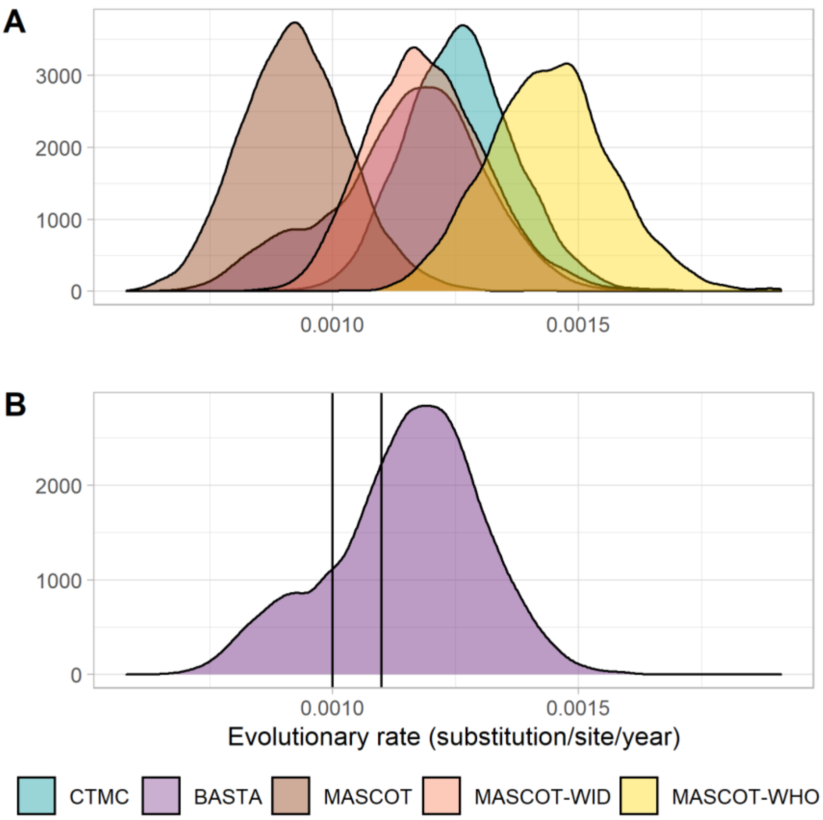


**Figure S23. Posterior kernel density distribution of the evolutionary rate estimated on the SARS-CoV-2 data set. A**: Posterior kernel density distribution of the evolutionary rate estimated by all algorithms. BASTA posterior distribution is bimodal with its major mode close to the estimates of CTMC and MASCOT-WID, and the second mode close to the estimate of MASCOT. **B**: Focus on the posterior kernel density distribution of BASTA. Due to the bimodality of the posterior density distribution, we split in two the tree posterior distribution according to the value of the evolutionary rate. The major mode corresponds to posterior samples with an evolutionary rate higher than 11e-4 substitution.site$^{-1}$.year$^{-1}$ and the minor mode to posterior samples with an evolutionary rate lower than 10e-4 substitution.site$^{-1}$.year$^{-1}$.

#### 5. Simulation framework of RABV epidemics

**Table S2. Values of simulation parameters.**

| Notation | Parameter description | Value | | Source |
|---|---|---|---|---|
| $b$ | **Dog birth rate per day** | 1/365 | | Assumption |
| $d$ | **Dog death rate per day** | 1/365 | | Assumption |
| $\beta$ | **Rabies transmission rate** | 3.2 | | Assumption |
| $H_i$ | **Human population size per region** | 7 demes<br>Region 1: 4,917,672<br>Region 2: 2,208,003<br>Region 3: 11,913,790<br>Region 4: 5,773,588<br>Region 5: 3,431,383<br>Region 6: 5,023,878<br>Region 7: 672,319 | 3 demes<br>Region 1: 10,557,059<br>Region 2: 17,687,379<br>Region 3: 5,696,197 | WorldPop (2) |
| $r_d$ | **Dog:human ratio** | 0.1 | | Assumption |
| $C_s$ | **Scaling factor** | 1.00E-08 | | Assumption |
| $v_{i \to j}$ | **Contact matrix** | - | | Radiation model (3,4) |
| $\gamma$ | **Infectious period** | Discretized gamma distribution from 1 to 15 days<br>$\Gamma(3, 1.1)$ | | Hampson et al., 2009 (5) |
| $\varepsilon$ | **Incubation period** | $\Gamma(2, 11.055)$ | | Hampson et al., 2009 (5) |
| $\mu$ | **Mutation rate** | 2.44e-4 $subs.site^{-1}.yr^{-1}$ | | Troupin et al., 2016 (6) |
| $\kappa$ | **Transition/transversion ratio** | 2 | | Assumption |
| $\pi_A$ | **Base A frequency in the reference genome** | 0.2852 | | Marston et al., 2013 (7) |
| $\pi_C$ | **Base C frequency** | 0.2198 | | Marston et al., 2013 (7) |
| $\pi_G$ | **Base G frequency** | 0.2313 | | Marston et al., 2013 (7) |
| $\pi_T$ | **Base T frequency** | 0.2638 | | Marston et al., 2013 (7) |

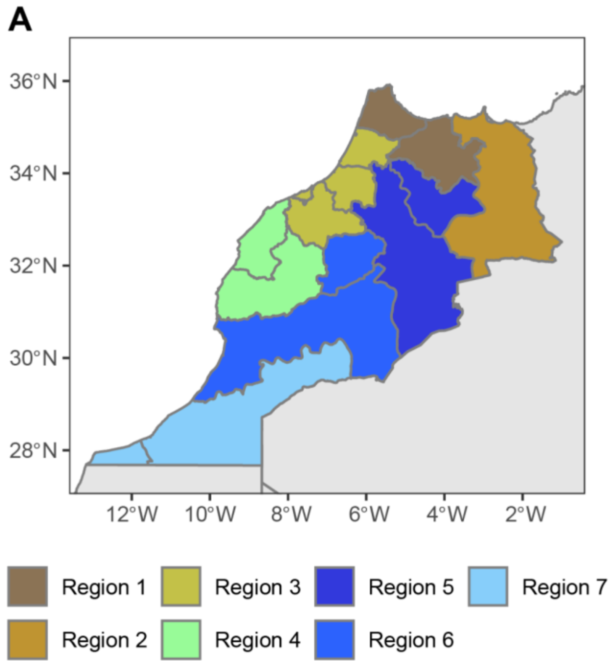


**Figure S24. Partition of Morocco into three and seven arbitrary locations. A**: Partition of Morocco into seven arbitrary locations. The major results corresponding to this framework are presented in the main text. All epidemics started by the introduction of a single index case in Region 2. Regions 3 and 4 correspond to over-sampled locations. **B**: Partition of Morocco into three arbitrary locations. The main results corresponding to this framework are presented exclusively in the Supplementary Materials. All epidemics started by the introduction of a single index case in Region 1. Region 2 corresponds to the over-sampled location. The gray outlines delineate the official regions downloaded from GADM and the colors indicate the arbitrary locations.

#### 6. Bayesian inference

##### a. Simulation study

**Table S3. Prior distributions used in the simulation study for the CTMC, BASTA, MASCOT, and MASCOT-GLM.**

| | Parameter | CTMC[a] | BASTA | MASCOT | MASCOT-GLM |
|---|---|---|---|---|---|
| **HKY substitution model** | **Transition-transversion ratio** | Lognormal(1, 1.25) | Lognormal(1, 1.25) | Lognormal(1, 1.25) | Lognormal(1, 1.25) |
| | **Base frequencies** | Dirichlet(alpha=1, sum=1) | Uniform([0:1]) | Uniform([0:1]) | Uniform([0:1]) |
| **Molecular clock** | **Clock rate** | CTCM Rate reference (8) | Lognormal(0, 4) | Lognormal(0, 4) | Lognormal(0, 4) |
| **Spatial model** | **Migration rate** | Exponential(1) | Exponential(1) | Exponential(1) | Exponential(1) |
| | **Migration clock** | CTMC Rate reference (8) | | Set to 1 | Exponential(1) |
| | **Coefficient of migration predictor (GLM)** | | | | Normal(0,1) |
| | **Root location frequency (CTMC)** | Uniform([0:1]) | | | |
| | **Deme population size (constant over time)** | 1/X[a] | Exponential(1) | Exponential(1) | |
| | **Deme size clock** | | | | Exponential(1) |
| | **Coefficient of deme size predictor (CTMC)** | | | | Normal(0,1) |
| | **Equal deme population sizes** | | Yes | Yes | |
| **BSSVS** | **Sum of non-zero migration rates** | Poisson($n_{demes}$-1) | Poisson($n_{demes}$-1) | Poisson($n_{demes}$-1) | |
| | **Sum of included predictors on migration rates** | | | | P(1) |
| | **Sum of included predictors on deme sizes** | | | | P(1) |

[a]We used the default priors from BEAUTi v1.10.4.

Abbreviations: BSSVS, Bayesian stochastic search variable selection; GLM, Generalized linear mode; HKY model, Hasegawa, Kishino, and Yano model.

**Table S4. Indicative number of iterations per hour for the four discrete phylogeographic approaches according to the number of demes and genomes. The number of iterations is expressed in millions.**

| No. of demes | No. of genomes | CTMC | BASTA | MASCOT | MASCOT-GLM |
|---|---|---|---|---|---|
| **3** | **150** | 14.85 | 0.67 | 5.23 | 2.4 |
| | **500** | 2.22 | 0.21 | 0.93 | 0.69 |
| **7** | **150** | 14.08 | 0.70 | 2.4 | 1.3 |
| | **500** | 1.85 | 0.12 | 0.32 | 0.23 |

**Table S5. Number of chains excluded per algorithm for the main analysis on seven demes.**

| No. of genomes | CTMC | BASTA | MASCOT | MASCOT-GLM |
|---|---|---|---|---|
| **150** | 0 | 17 | 6 | 0 |
| **500** | 0 | NA | 161 | 2 |

Abbreviations: NA, Not applicable.

**Table S6. Number of chains excluded per algorithm for the supplementary analysis on three demes.**

| No. of genomes | CTMC | BASTA | MASCOT | MASCOT-GLM |
|---|---|---|---|---|
| **150** | 0 | 0 | 0 | 0 |
| **500** | 0 | NA | 2 | 0 |

Abbreviations: NA, Not applicable.

b. **Analysis of the rabies dataset**

**Table S7. List of priors used for each discrete phylogeographic approach on the RABV data set.**

| | Parameter | CTMC | BASTA | MASCOT |
|---|---|---|---|---|
| **HKY substitution model** | **Transition/transversion ratio** | Lognormal(1, 1.25) | Lognormal(1, 1.25) | Lognormal(1, 1.25) |
| | **Base frequencies** | Empirical | Empirical | Empirical |
| | **Shape of the gamma rate of heterogeneity** | Exponential(0.5) | Exponential(0.5) | Exponential(0.5) |
| **Lognormal relaxed molecular clock** | **Mean** | CTCM Rate reference | Lognormal(0.001, 1000) | Lognormal(0.001, 1000) |
| | **Standard deviation** | Exponential(1/3) | Exponential(1/3) | Exponential(1/3) |
| **Spatial model** | **Migration rate** | Exponential(1) | Exponential(1) | Exponential(1) |
| | **Migration clock** | CTMC Rate reference | | |
| | **Region frequency (CTMC)** | Uniform([0:1]) | | |
| | **Deme population size (constant over time)** | Gamma(shape=0.001, scale=1000) | Exponential(1) | Exponential(1) |
| | **Regions root frequencies** | Uniform([0:1]) | | |
| | **Equal deme population sizes** | | Yes | Yes |
| **BSSVS** | **Sum of non-zero migration rates** | Poisson(5) | Poisson(5) | Poisson(5) |

Abbreviations: BSSVS, Bayesian stochastic search variable selection; HKY model, Hasegawa, Kishino, and Yano model.

##### c. Analysis of the SARS-CoV-2 dataset

**Table S8. List of priors used for each discrete phylogeography algorithm on the SARS-CoV-2 data set.**

| | Parameter | CTMC | BASTA | MASCOT | MASCOT-WID MASCOT-WHO |
|---|---|---|---|---|---|
| **HKY substitution model** | **Transition/transversion ratio** | Lognormal(1, 1.25) | Lognormal(1, 1.25) | Lognormal(1, 1.25) | Lognormal(1, 1.25) |
| | **Base frequencies** | Empirical | Empirical | Empirical | Empirical |
| | **Proportion of invariant** | Uniform([0,1]) | Uniform([0,1]) | Uniform([0,1]) | Uniform([0,1]) |
| | **Shape of the gamma rate of heterogeneity** | Exponential(0.5) | Exponential(0.5) | Exponential(0.5) | Exponential(0.5) |
| **Strict molecular clock** | **Evolutionary rate** | CTCM Rate reference | Lognormal(0, 4) | Lognormal(0,4) | Lognormal(0,4) |
| **Exponential growth coalescent model** | **Deme size** | Gamma(shape=0.001, scale=1000) | | | |
| | **Exponential growth rate** | Laplace(mean=0, scale=1) | | | |
| **Spatial model** | **Migration rate** | Exponential(1) | Exponential(1) | Exponential(1) | |
| | **Migration clock** | CTMC Rate reference | | | |
| | **Region frequency (CTMC)** | Uniform([0:1]) | | | |
| | **Regions root frequencies** | Uniform([0:1]) | | | |
| | **Deme size** | | Exponential(1) | Exponential(1) | |
| | **Equal deme population sizes** | | Yes | Yes | |
| | **Migration clock** | | | | Exponential(1) |

| | | | | | |
|---|---|---|---|---|---|
| **GLM spatial model** | **Migration predictors scaler** | | | | Normal(0,1) |
| | **Deme size clock** | | | | Exponential(1) |
| | **Deme size predictors scaler** | | | | Normal(0,1) |
| **BSSVS** | **Sum of non-zero migration rates** | Poisson(5) | Poisson(5) | Poisson(5) | |
| | **Sum of non-zero deme size predictors** | | | | Poisson(1) |
| | **Sum of non-zero migration rate predictors** | | | | Poisson(1) |

Abbreviations: BSSVS, Bayesian stochastic search variable selection; GLM, Generalized linear model; HKY model, Hasegawa, Kishino, and Yano model.

#### References

1. Lemey P, Hong SL, Hill V, et al. Accommodating individual travel history and unsampled diversity in Bayesian phylogeographic inference of SARS-CoV-2. *Nat. Commun.* 2020;11(1):5110.

2. WorldPop. WorldPop project. (http://worldpop.org.uk/)

3. Simini F, González MC, Maritan A, et al. A universal model for mobility and migration patterns. *Nature*. 2012;484(7392):96–100.

4. Golding N, Schofield A, Kraemer MUG. Movement: Functions for the analysis of movement data in disease modelling and mapping. *R Packag. version 0.2*. 2015.

5. Hampson K, Dushoff J, Cleaveland S, et al. Transmission dynamics and prospects for the elimination of canine Rabies. *PLoS Biol.* 2009;7(3):0462–0471.

6. Troupin C, Dacheux L, Tanguy M, et al. Large-Scale Phylogenomic Analysis Reveals the Complex Evolutionary History of Rabies Virus in Multiple Carnivore Hosts. *PLoS Pathog.* 2016;12(12):e1006041.

7. Marston DA, McElhinney LM, Ellis RJ, et al. Next generation sequencing of viral RNA genomes. *BMC Genomics*. 2013;14(1):444.

8. Ferreira MAR, Suchard MA. Bayesian analysis of elapsed times in continuous-time Markov chains. *Can. J. Stat.* 2008;36(3):355–368.
